# Actin cross-linking organizes basal body patterning through anomalous diffusion transitions

**DOI:** 10.64898/2026.05.14.725088

**Authors:** Raghavan Thiagarajan, Younes Farhangi Barooji, Poul-Martin Bendix, Mandar M. Inamdar, Jakub Sedzinski

## Abstract

Subcellular protein complexes and organelles exhibit diverse dynamic behaviors that reflect the mechanical constraints and organization of the intracellular environment. Although some structures follow classical Brownian motion, many display anomalous dynamics, including subdiffusion and superdiffusion, driven by viscoelasticity, molecular crowding, and cytoskeletal interactions. Transitions between these regimes are increasingly recognized as critical for subcellular organization, yet how they are regulated and influence pattern formation remains unclear. Here, we investigate the spatial arrangement of cilia on the apical surface of multiciliated cells (MCCs) in developing *Xenopus laevis* embryos, where coordinated ciliary beating depends on the precise organization of hundreds of centriole-derived basal bodies (BBs). Using quantitative confocal, high-resolution and high-speed TIRF imaging together with theoretical modeling, we show that BB trajectories undergo time-resolved transitions between diffusive and anomalous motion, with distinct regimes that correlate with apical surface expansion. During the early stages, actin remodeling facilitates the dispersal of BBs by providing a permissive, low-confinement environment. As development progresses, the actin network becomes increasingly cross-linked, forming a dense meshwork that constrains BB movement and promotes uniform spacing across the apical domain. Disruption of *α*-actinin-1, a major actin cross-linking protein, impairs the integrity of the apical actin meshwork, weakens BB confinement, and disrupts regular spatial patterning, ultimately compromising the spatial arrangement of BBs required for proper cilia alignment. Together, we show that progressive apical actin cross-linking coordinates BB positioning and regulates their dynamic state, guiding the shift from diffusive to confined motion. This transition in dynamics enables the emergence of a uniform BB pattern, which in turn ensures the aligned deployment of motile cilia necessary for effective directional fluid flow.

## Introduction

The subcellular environment is a dynamic and heterogeneous medium in which particles, ranging from membraneless macromolecular assemblies to membrane-bound organelles, exhibit motion that often deviates from classical Brownian behavior^1–3^. Such anomalous dynamics, including subdiffusion and superdiffusion, can arise from molecular crowding, spatial heterogeneity, or active forces generated by cytoskeletal networks ^4–8^. These non-Brownian modes of motion are functionally important, contributing to subcellular organization, cargo trafficking, and spatial patterning within cells^9–11^.

The actin cytoskeleton, a dynamic and viscoelastic network, plays a central role in the shaping of intracellular dynamics by providing mechanical constraints and generating contractile and extensile forces^12–16^. Remodeling of actin filaments, cross-linking, and interactions with molecular motors enable transitions between diffusive and anomalous motion^6,17,18^. Among the many subcellular structures regulated by actin mechanics, centrioles and their mature form, basal bodies (BBs), serve as a powerful model for investigating how cytoskeletal remodeling governs organelle positioning and collective pattern formation.

Centrioles are conserved microtubule-based organelles that function across diverse cell types to organize microtubule arrays, facilitate mitotic spindle formation, and establish planar cell polarity and directional migration ^19–28^. In ciliated cells, the mother centriole matures into a BB that anchors and nucleates a cilium^29–32^. Multiciliated cells (MCCs), a specialized epithelial cell type, generate hundreds of such BB and associated motile cilia on their apical surface^33–35^. These cilia drive directional fluid flow in various tissues: enabling mucociliary clearance in the airway^36,37^, transporting ova through the oviduct^38,39^, and circulating cerebrospinal fluid in the brain ventricles^40^.

The precise positioning of the BBs on the apical surface is essential for coordinated ciliary beating and effective fluid flow generation^41–44^. Disruption of BB patterning alters mucociliary clearance and contributes to ciliopathies that affect the respiratory, reproductive, and nervous systems^45–50^. Understanding how BBs are spatially organized during MCC development remains a central question in cell and developmental biology.

In particular, the BB arrangements within the apical domain of MCC are tissue specific: unipolar in the brain ependyma^51^, linear in the mammalian airway epithelium^52,53^, and uniformly distributed in the *Xenopus* embryonic epidermis^54,55^. These diverse spatial patterns reflect an emergent organization governed by tissue-specific molecular and mechanical cues. BB positioning has been associated with planar cell polarity pathways, cytoskeletal dynamics, and structures associated with BB, such as the basal feet^47,52,54,56^. In both *Xenopus* embryonic MCCs and mammalian airway MCCs, the number of centrioles scales with the apical surface area, indicating a conserved coupling between organelle biogenesis and cell geometry^57,58^. However, the dynamic principles by which BBs self-organize within the apical domain remain poorly understood.

Emerging evidence suggests that the apical actin cytoskeleton plays a crucial role in regulating BB positioning. In *Xenopus* MCCs, the assembly of the apical actin after centriole amplification drives the expansion of the apical surface and provides a mechanical and structural scaffold for the BB docking^59–61^. Proteins such as WDR5 and Filamin-A promote actin polymerization and cross-linking around BBs^62,63^, while disruption of the apical actin meshwork disrupts the BB docking and spatial organization^54,64^. These observations suggest that the apical actin network not only scaffolds BBs but also imposes mechanical constraints that influence their motility and final distribution.

Viewed as active particles embedded in an evolving viscoelastic medium, BBs offer a unique model for exploring how the cytoskeletal architecture shapes organelle positioning. Their behavior resembles that of active colloids, where spatial order arises from local interactions and mechanical feedback with the surrounding matrix^65–67^. Modeling studies have shown that BBs can self-organize through cytoskeleton-dependent interactions^52,68^, yet how these processes are regulated by the dynamic and cross-linked architecture of the apical actin network remains largely unexplored.

Here, we demonstrate that remodeling of the apical actin cytoskeleton, particularly through cross-linking, governs the dynamic behavior and spatial organization of BB during the development of *Xenopus laevis* MCCs. By combining quantitative confocal imaging with high-speed TIRF microscopy, we show that BBs exhibit in-plane motility characterized by anomalous diffusion, transitioning from free to confined dynamics as the apical domain expands. The perturbation of actin cross-linking through *α*-actinin-1 KD disrupts both the confinement of the BB and regular spatial patterning by altering the actin meshwork, which normally forms pocket-like structures that embed and position the BBs within the apical domain. Theoretical modeling reveals that actin-mediated spatial bias and repulsive interactions imposed by the cross-linked network are required to reproduce the emergent BB distribution. Together, these findings establish a direct mechanistic link between the architecture of apical actin and BB dynamics, highlighting how cytoskeletal remodeling influences the positioning and patterning of subcellular organelles.

## Results

### Basal body patterning is synchronized with apical surface expansion

The formation of MCCs in the *X. laevis* embryonic epidermis involves a coordinated morphogenetic program known as radial intercalation^69,70^, during which the progenitors of MCCs insert into the outer epithelial layer and expand their apical domain in a process called apical emergence^60^. This apical surface is essential for the generation of motile cilia, which are nucleated from the BB after their ascent, docking, and distribution across the apical domain (Fig. 1a).

**Figure 1.**
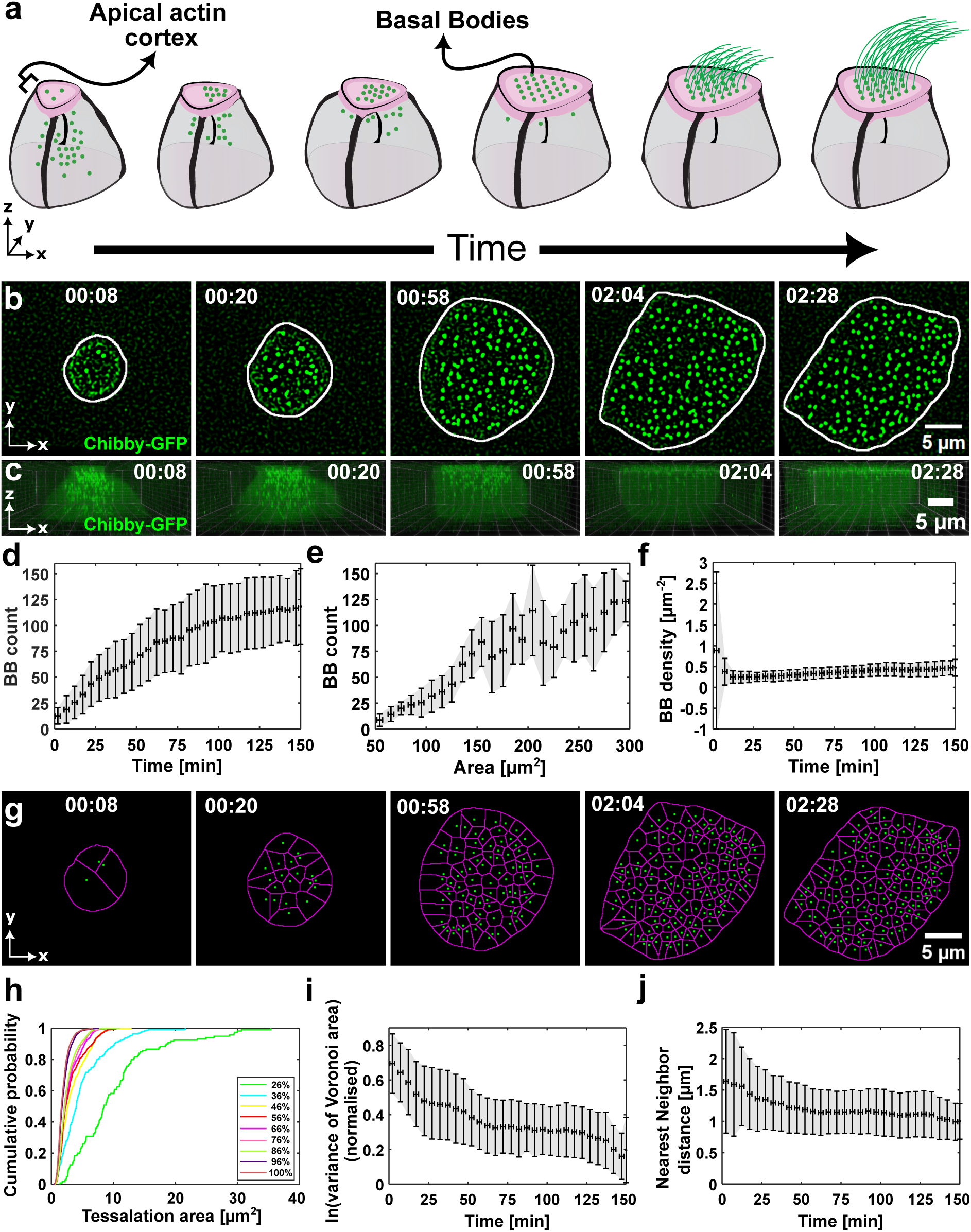
Basal body patterning is synchronized with apical surface expansion. All data correspond to coarse-timed imaging (frame interval = 30–60 s; total duration = 2.5 h). (**a**) Schematic showing the sequence of BB dynamics during multiciliated cell (MCC) morphogenesis. BBs ascend from the basal to the apical side, followed by distribution and patterning within the apical plane. BBs are shown in green, and the apical actin cortex is shown in pink. (**b**) Time-lapse sequence of BBs (Chibby-GFP) within the apical domain, with the apical periphery outlined by a white contour (see Movie 1). (**c**) Corresponding sequence showing the ascent of the BB from the basal to apical side of the MCC (see Movie 2). (**d**) Number of BBs plotted against time. (**e**) Number of BBs plotted against the apical area. (**f**) BB density over the apical area plotted against time. (**g**) Voronoi tessellations of BBs for the time-lapse sequence shown in (b) (see Movie 3). (**h**) Cumulative distribution functions (CDFs) of Voronoi tessellation areas for every 10% increase in the apical area, for the data shown in (g). (**i**) Variance of Voronoi tessellation areas plotted against time. Variance values are natural log–transformed (ln) and min–max normalized. (**j**) The Nearest-neighbor distance of the BBs plotted against time. In (**d**), (**e**), (**f**), (**i**) and (**j**), error bars represent mean ± standard deviation (SD) of data averaged over 9 cells from 5 experiments, using 5 min (time) or 10 µm^2^ (area) bins. In (**b**), (**c**), and (**g**), time is in hh:mm.

To investigate BB dynamics during apical emergence, we performed early-stage injections in *X. laevis* embryos to drive the MCC-specific expression of fluorescently tagged proteins. Chibby, a centriolar and BB-associated protein required for BB docking and ciliogenesis^71^, was used to label BB, while LifeAct^72^, an F-actin marker, was used to visualize the apical expansion. Both constructs were expressed under the control of the MCC-specific *α*-tubulin promoter^73^ (Fig. 1b, 1c, S1a, Movie 1). Confocal microscopy enabled 3D time-lapse imaging of MCCs throughout the entire BB ascent, docking, and spatial organization process (Fig. 1b, 1c, Movie 2). These “coarse-timed” recordings were acquired at intervals of 30 or 60 s.

We focus our analysis on the 2.5-hour window during which apical emergence occurs, a phase previously characterized by a near-linear increase in the apical area followed by a plateau (Fig. S1b)^60^. During this period, BBs exhibited directed migration from the basal to the apical side of the cell (Fig. 1c) and progressively docked at the apical membrane (Fig. 1b). Quantification of the appearance of the apical BBs revealed a linear accumulation over time (Fig. 1d, 1e), while the density of the BBs remained constant throughout the expansion phase (Fig. 1f). This indicated that BB delivery was closely coordinated with apical surface growth. Following docking, the BBs were redistributed throughout the apical domain (Fig. 1b), suggesting an additional level of spatial organization beyond the initial insertion on the apical side, prompting further investigation of their spatial distribution patterns.

Previous studies examining the final spatial organization of BBs in *X. laevis* embryos have shown that they adopt a characteristic spacing of approximately ∼1 µm^54,74^. To understand how this pattern emerges over time, we quantified the dynamics of the BB distribution using Voronoi tessellation (Fig. 1g, Movie 3). At early time points, the tessellation pattern was heterogeneous, featuring a mixture of large and small areas. As the apical surface expanded and became progressively populated with BBs, the pattern transitioned to a more homogeneous configuration (Fig. 1g, 1h). This transition was quantitatively reflected in a decreasing variance of the tessellation areas over time (Fig. 1i). When analyzed as a function of apical surface area, the variance of the tesselation areas decreased during the early expansion phase (<150 µm^2^) and reached a plateau during later stages of expansion (>150 µm^2^) (Fig. S1c).

To further characterize the spatial arrangement of BBs, we measured inter-BB distances using two complementary metrics: *minimum distance*, which captures local packing precision, and *pairwise distance*, which reflects overall spacing tendencies (*see methods section 7.3.4 in SI*) (Fig. S1d). The minimum distance (or nearest neighbor) showed a consistent decline over time (Fig. 1j). When plotted against the apical area, it followed a similar trend: decreasing during the initial expansion phase and stabilizing beyond 150 µm^2^ (Fig. S1e). This trend mirrored the behavior observed in the tessellation area variance. In particular, in the final stages of expansion, the nearest-neighbor distances converged to ∼1 µm (Fig. 1j, S1e), aligning with previous observations of the mature BB spacing. To confirm this, we computed the relative probability distribution of all nearest-neighbor distances across the entire apical expansion timeline. A dominant peak at ∼1 µm was evident, reinforcing this value as the preferred spacing between BBs (Fig. S1f).

Next, we analyzed pairwise distances to gain insight into how the global BB spacing evolved in relation to apical growth. This analysis revealed a gradual increase in the pairwise distance with the apical area, suggesting that the BBs became more widely distributed as the surface expanded (Fig. S1g). Despite this general spread, the dominant pairwise distance remained centered around ∼6 µm (Fig. S1h), indicating that the most likely distance between any two BBs was consistently around ∼6 µm.

Together, these findings demonstrate that the ascent, docking, and distribution of BB occur simultaneously with the apical expansion. The observation that BB density remains constant while both the tessellation area variance and nearest-neighbor distance decline from the onset suggests that MCCs initiate BB positioning with high spatial precision early in apical emergence. The plateauing of these parameters mid-way through expansion further implies that the final BB distribution is established well before the apical surface reaches its full size. Collectively, these results point to tight coordination between the mechanisms governing apical expansion and BB distribution.

### Passive and active mechanisms shape basal body distribution

To understand the mechanisms governing the distribution of BBs and their relationship to apical expansion, we characterized BB in-plane dynamics at the apical domain. Using time-lapse imaging, we tracked the movements of BB during apical surface expansion at coarse-timed intervals of 30 or 60 seconds (Fig. 2a, Movie 4). This approach enabled us to examine the spatiotemporal properties of the BB entry and how their trajectories evolve toward a uniform distribution. BBs were found to insert into the apical domain at a steady rate of approximately 2 BBs per minute, with only minor temporal fluctuations (Fig. S2a).

**Figure 2.**
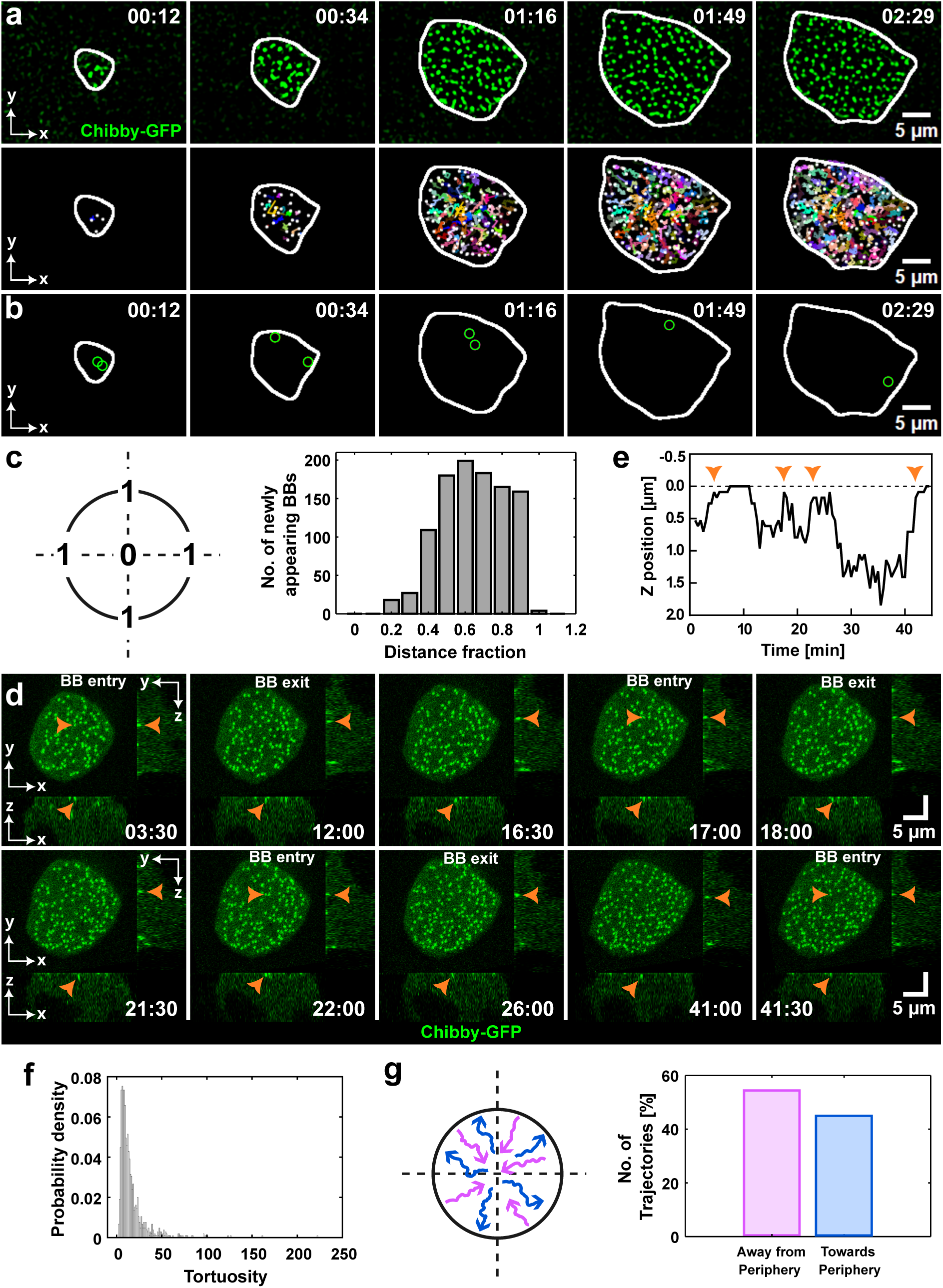
Passive and active mechanisms shape BB distribution. All data correspond to coarse-timed imaging (frame interval = 30–60 s; total duration = 2.5 h). (**a**) First row: time-lapse sequence of BBs (Chibby-GFP) in the apical domain. Second row: trajectories of BBs for the sequence above (see Movie 4). (**b**) Appearance of BBs near the apical periphery. The circles mark the BBs, and the white outline marks the apical boundary (see Movie 5). (**a**) and (**b**) share the same time-lapse sequence where the apical periphery is outlined by a white contour. (**c**) The schematic shows the distance fraction where 0 corresponds to the center and 1 to the periphery of the apical domain. Plot shows the distribution of newly appearing BBs across distance fractions within the apical domain. (**d**) Time-lapse sequence showing the entry and exit of a BB in the apical domain. Each panel shows views in xy (top), xz (bottom), and yz (right); the orange arrow highlights the tracked BB and the time corresponds to the plot in (e) and Movie 6. (**e**) Plot showing the change in axial Z position of the BB shown in (d) over time. The dotted line at zero marks the apical surface. Arrows indicate the positions at which the BB approaches or reaches the apical surface during transient entry events before subsequently exiting. The final arrow marks the entry event after which the BB remains stably docked at the apical surface. (**f**) Probability density of trajectory tortuosity with a peak at 9.6. (**g**) The schematic illustrates BB trajectories directed towards (blue) and away (pink) from the periphery. The plot shows the percentage of trajectories moving in each direction. Data in (**c**), (**f**), and (**g**) are pooled from 9 cells across 5 experiments. In (**f**) and (**g**), a total of 1052 BB trajectories were analyzed across these cells. In (**a**) and (**b**), time is in hh:mm; in (**d**), time is in mm:ss.

Next, we asked whether there was a spatial bias in BB docking. To address this, we extracted the initial coordinates of the newly appearing BBs and classified them based on their position within the apical domain - either centrally or peripherally located (Fig. S2b; *see methods section 7.3.6 in SI*). Interestingly, new BBs appeared preferentially in the periphery of the apical domain (Fig. 2b, 2c, Movie 5). Given that the apical periphery undergoes continuous expansion outward, this spatial preference suggests that the insertion of the BB is coupled to the extension of the apical domain. Thus, BB appearance and docking proceed at a constant rate, but occur preferentially within newly extended regions of the apical surface.

To further explore the spatial regulation of BB insertion, we quantified the positions of new BBs relative to: (i) BBs that had appeared in previous time points (Fig. S2c) and (ii) pre-existing BBs present at the same time point (Fig. S3a), using both minimum and pairwise distance metrics. For the first case, we found that the minimum and pair-wise distances between new BBs and those from preceding time points peaked at 5-6 µm (Fig. S2d, S2f). Moreover, these distances increased progressively during apical expansion (Fig. S2e, S2g), indicating that BBs tend to appear in newly expanded, spatially unoccupied regions. For the latter case, the minimum distance between newly inserted BBs and the pre-existing BB population at the same time point, peaked at ∼1 µm (Fig. S3b) and plateaued at a similar value at the end of the expansion process (Fig. S3c). These findings are consistent with nearest-neighbor distance measurements among all BBs, which also peaked around ∼1 µm (Fig. 1j, S1e). In contrast, pairwise distances peaked at ∼6 µm and exhibited a steady increase over time (Fig. S3d, S3e).

Together, these data suggest that BBs maintain a minimum inter-BB spacing of 1 µm, while the global spacing increases toward a preferred distance of 6 µm as the apical domain expands. This pattern is also evident in the increasing pairwise distances between all subpopulations of BBs, including pre-existing BBs (Fig. S1g, S1h), newly inserted BBs from successive time points (Fig. S2f, S2g), and new vs. pre-existing BBs (Fig. S3d, S3e), suggesting an emergent spatial pattern in the organization of BBs. At the same time, the steady increase in pairwise distances supports the role of apical expansion as a driving force in the redistribution of BBs.

To understand how the BB distribution evolves over time, we quantified the characteristics of the BB trajectories. The contour lengths varied from approximately 5 to 150 µm (Fig. S3f), with longer tracks reflecting complex, convoluted paths and shorter tracks suggesting limited mobility or transient presence. However, we ensured data integrity by implementing a strict tracking and filtering pipeline, that excluded spurious short trajectories thereby restricting the remaining short-lived events to genuine transient presence (*see methods section 7.2.4 in SI*). Manual inspection further revealed that some BBs exited and re-entered the apical surface during expansion (Fig. 2d, 2e, Movie 6). These exit–entry events occurred multiple times though with substantial temporal variance (Fig. S3i). In particular, exit behavior was observed only after the apical domain exceeded an area of ∼140 µm^2^ and persisted thereafter, suggesting that the retention of BB becomes less stable beyond a certain domain size.

Interestingly, exiting BBs were often confined to specific subregions and exhibited minimal displacement (Movie 6). This observation prompted us to ask whether some BBs remained stationary throughout the expansion phase, potentially contributing to the population of trajectories with shorter end-to-end lengths (Fig. S3g). In fact, analysis of step size distributions revealed a strong probability peak at ∼0 µm (Fig. S3h), which supports the presence of stationary BBs. At the same time, the observed distribution of tortuosity values was consistently greater than 1 (Fig. 2f), indicating that BB paths were non-linear, i.e., the contour lengths (Fig. S3f) consistently exceeded their end-to-end displacements (Fig. S3g). To test for directional bias in the movement of the BBs, we computed the dot product of the displacement and position vectors, which revealed no preferred orientation of movement towards or away from the expanding periphery (Fig. 2g, S3j).

In summary, not all BBs are retained stably within the apical domain during expansion. They can display prolonged stationary behavior, undergo exit and re-entry cycles, or follow highly convoluted and non-directional trajectories. Consistent with these observations, the dynamics of BB exhibited mixed characteristics of both advective and diffusive behavior, as inferred from the alignment of the trajectory and the average of drift (Fig. S3k, S3l) and the analysis of the mean square displacement (Fig. S3m), respectively. These results collectively indicate that, although apical expansion influences BB movements, it is not the sole determinant; The distribution of BB emerges through a combination of passive spreading and additional active regulatory mechanisms.

### Basal body dynamics shift with apical domain expansion

Apical expansion in MCCs is driven by the assembly of the actin network^60^. Simultaneously, the BBs ascend from the cell base, reach the apical surface, and become embedded within the actin cortex^56^. At the end of the apical expansion, the BBs occupy only 20% of the apical area, while the remaining 80% is predominantly composed of actin (Fig. S1a, S4a). Based on this spatial distribution, we hypothesized that actin may play an active role in the regulation of BB dynamics and patterning.

Since actin remodeling occurs on timescales from seconds to tens of seconds^75–77^, we reasoned that key features of BB movement may be missed at the coarse-imaging intervals (30–60 s) used previously. To resolve BB behavior at finer temporal resolution, we used a TIRF setup equipped with a high-speed camera to acquire 30,000-frame movies (10.5 minutes duration) at 21 ms per frame (Fig. 3a, Movie 7).

**Figure 3.**
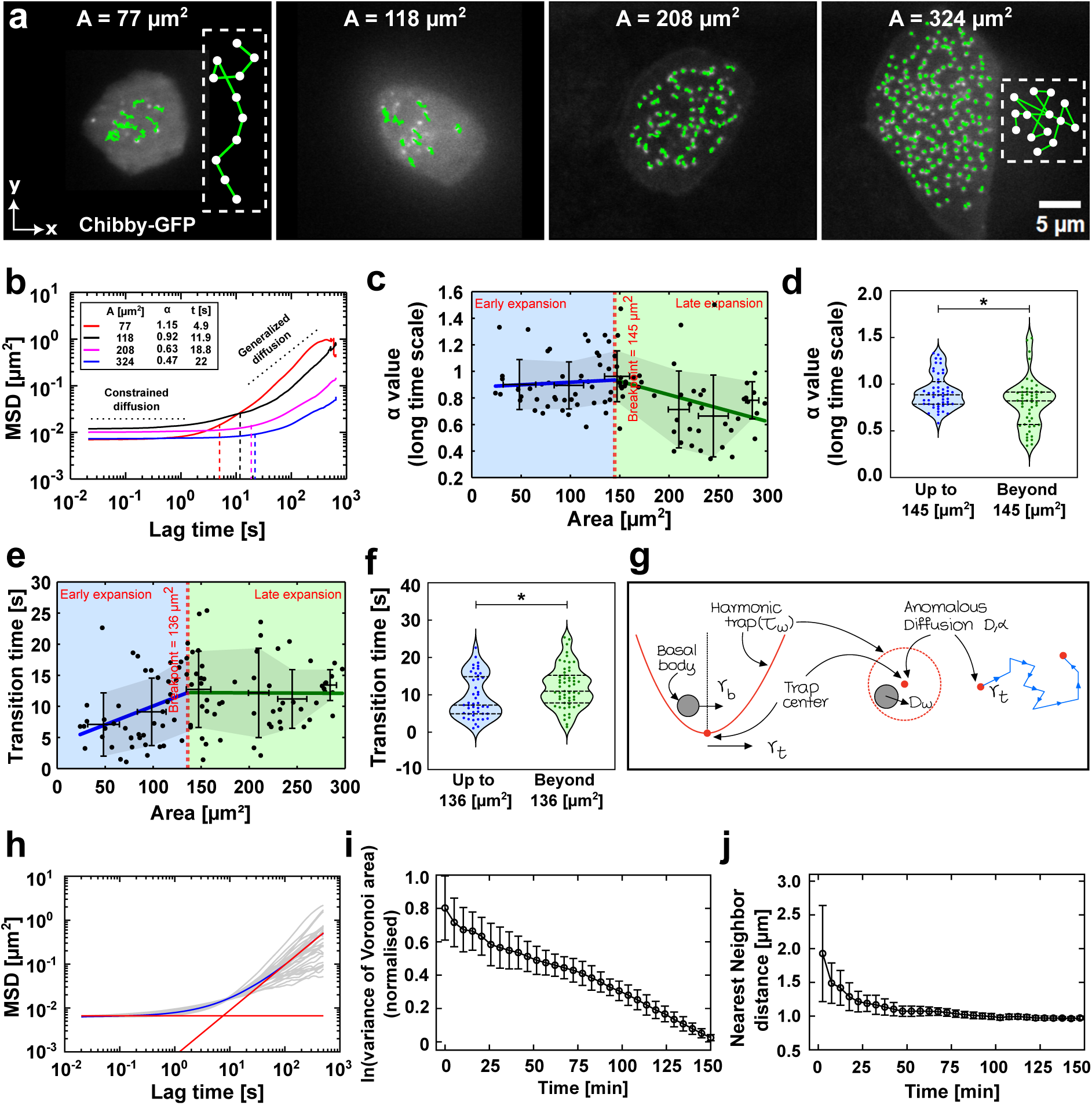
Basal body dynamics shift with apical domain expansion. Data in (**a-f**) correspond to fine-timed imaging (frame interval = 21 ms; 30,000 frames; total duration = 10.5 min). (**a**) Images of four apical domains of different sizes, shown at the last time point of a 10.5 minute fine-timed image sequence. BBs (Chibby-GFP) are shown with their overlaid trajectories. The inset in the first image shows a schematic illustrating elongated trajectories in smaller apical domains, while the inset in the last image highlights convoluted trajectories in larger domains (see Movie 7). (**b**) MSDs of BBs from cells shown in (a), plotted against lag time. Each curve represents the ensemble-averaged MSD of trajectories within a single cell. The vertical dotted lines mark the transition times. Reported for each curve are the apical area, *α* value, and transition time. The black dotted lines above the curves indicate the phases of short- and long-timescales, depicting constrained diffusion at short times and a mix of sub-, diffusive, and superdiffusive behaviors at long times. (**c**) Long-term *α* values plotted against the apical area. (**d**) Comparison of *α* values before and after the area breakpoint in (c); exact P = 0.0144. (**e**) Transition time plotted against the apical area. (**f**) Comparison of transition times before and after the area breakpoint in (e); exact P = 0.0173. In (**c**) and (**e**), each dot represents the ensemble average per cell; error bars represent mean ± SD of averaged data, based on 4,604 trajectories from 100 cells across 17 experiments, using 50 µm^2^ bins. The red dotted line marks the breakpoint (*α* value - 145 µm^2^; transition time – 136 µm^2^) identified by piecewise linear regression (*see methods section 8.3.5 in SI*) while blue and green lines are regression fits before and after the breakpoints. The blue and green shaded regions denote data before and after the breakpoints and serve as visual guides for the early (<150 µm^2^) and late (>150 µm^2^) phases of apical expansion. Data for (**d**) include 837 trajectories from 50 cells (≤ 145 µm^2^) and 3767 from 50 cells (>145 µm^2^), and for (**f**), 675 trajectories from 45 cells (≤ 136 µm^2^) and 3929 from 55 cells (>136 µm^2^), all from 17 experiments. In (**d** and **f**), the central line represents the median, and the upper and lower lines represent the 75^th^ and 25^th^ percentiles, respectively. (**g-j**) Model simulations. (**g**) Schematic of the particle trap arrangement in simulations showing initial caging followed by anomalous diffusion. (**h**) MSDs of simulated BBs: grey - individual particles; blue - ensemble average; red lines - fitted regimes; their intersection defines the transition time. Data obtained from 20 independent runs of the single-particle simulation. (**i**) Variance of Voronoi tessellation areas plotted against time. Variance values are natural log–transformed (ln) and min–max normalized. (**j**) The distance from the Nearest-neighbor plotted against time. In (**i** and **j**), error bars represent mean ± SD of data averaged over 20 simulations, using 5 min bins.

These fast acquisition movies were collected across the entire range of apical expansion - from 50 µm^2^ to 300 µm^2^ - and used to calculate mean squared displacement (MSD) plots for each BB trajectory within each cell. To extract morphogenetically relevant features while minimizing pseudoreplication and intrinsic variability, we averaged BB trajectories by ensemble at the single cell level, resulting in one averaged MSD plot per cell (Fig. 3a; Fig. 3b, Fig. S4b).

Across all apical areas and cells, the averaged MSD curves displayed similar qualitative behavior: At short timescales, they exhibited low slope values (*α* ≪ 1), indicating constrained motion; at longer timescales, the slopes increased and shifted into a range of subdiffusive (*α <* 1), diffusive (*α* ≈ 1), or superdiffusive (*α >* 1) regimes (Fig. 3b, Fig. S4b). In particular, while short-timescale *α* values were invariant across apical expansion (Fig. 3b, Fig. S4d), long-timescale *α* values varied with apical area (Fig. 3b, 3c), despite diffusion strength remaining constant (Fig. S4e).

Focusing on the long-timescale *α* values, we observed that BBs in smaller apical areas exhibited behavior close to free diffusion (*α* ≈ 1) (Fig. 3c). As the apical area increased, *α* gradually decreased to ∼ 0.6, indicating a transition to more restricted subdiffusive behavior. Piecewise linear regression revealed a breakpoint around 145 µm^2^ (Fig. 3c, Fig. S4f), indicating a change in slope between early (<150 µm^2^) and late (>150 µm^2^) phases of apical expansion (Fig. 3c, Fig. 3d). This behavioral transition was also evident in the morphology of the BB trajectories: in smaller areas, the trajectories appeared more elongated, while in larger areas they became increasingly convoluted (Fig. 3a, Movie 7). Consistently, end-to-end distances decreased with increasing area (Fig. S4g), while contour lengths remained constant (Fig. S4h), leading to increased tortuosity with apical expansion (Fig. S4i). Together, these data support a progressive change in the dynamics of the BB motility during the late phase of apical expansion.

Next, we examined the transition between short- and long-timescale MSD behaviors (Fig. 3b, Fig. S4c). At short timescales, *α* values were near zero (*α* ≪ 1), consistent with strong confinement or “trapping” of BBs. Over longer timescales, *α* values spanned a broader range, reflecting either weakening of this confinement (*α <* 1) or complete trap release (*α* ≈ 1 or *α >* 1). To verify that this initial trapping was not a measurement artifact, we tracked fluorescent beads of a size similar to BBs (0.5 µm) fixed on the coverslip. The MSD of immobilized bead controls, was approximately an order of magnitude lower than that of BBs, confirming that the observed BB trapping reflects genuine biological motion (Fig. S4j, *see methods section 8.3.6 in SI*).

We then quantified the time it took for the BBs to transition from trapped to released state (Fig. S4c) and found that this transition time was positively correlated with the apical area (Fig. 3b, Fig. 3e). Specifically, cells in the late phase of expansion showed significantly longer transition times (Fig. 3e, Fig. S4k, Fig. 3f), suggesting that BBs remain confined for longer periods in larger apical domains. These findings imply that the interaction of BB with its local environment becomes increasingly restrictive as the apical domain expands.

Taken together, our results demonstrate that both the behavior of the trajectory of BB (as quantified by *α*) and confinement dynamics (as measured by the transition time) evolve in parallel with the apical expansion. The consistent shift in both parameters near ∼140 µm^2^ (Fig. 3c, Fig. 3e) suggests a shared underlying mechanism driving this transition. The reduction in end-to-end displacement and the increase in tortuosity imply that BBs have greater spatial freedom during early expansion. In contrast, their movement becomes increasingly constrained in later stages. Moreover, the lengthening of the trap-release transition times supports the notion that BBs are held within confined regions for longer durations as expansion progresses.

### In silico modeling of BB dynamics

To gain further insight into the mechanisms driving the observed transitions in the dynamics of BBs and to explore the potential role of the actin network in this process, we next sought to replicate the behavior of BBs using theoretical modeling.

First, to mechanistically explain the observed two-regime behavior in BB dynamics in the previous section, we implemented a stochastic model in which BBs diffuse within harmonic traps that themselves undergo anomalous motion (Fig. 3g). In this framework, each BB is confined by a harmonic potential with diffusivity *D_w_*and trap relaxation time *τ_w_*, while the trap center follows fractional Brownian motion with diffusion strength *D* and anomalous exponent *α* ^78^. The model equations were integrated using an Euler-Maruyama algorithm. The Davis-Harte process corresponding to fractional Brownian motion (fBM)^79^ was used to obtain the increments for anomalous diffusion of the trap (*see Supplementary Information (SI) for complete mathematical formulation*). This framework successfully reproduced the experimentally observed MSD profiles: initial caging at short timescales followed by anomalous diffusion at longer timescales (Fig. 3h). The physical basis for this model aligns with our experimental observations that BBs are embedded within dynamically remodeling traps that locally confine them while itself undergoing stochastic reorganization, thereby permitting long-range anomalous exploration on extended timescales.

The coarse-grained experimental imaging (30 s intervals) operates at timescales significantly longer than the trap-release transition time (*τ* ≈ 5 − 15 s), thereby capturing only the anomalous diffusion regime without resolving the initial caging dynamics. Consequently, our computational model for the full spatiotemporal evolution of BB organization during apical emergence focuses on anomalous diffusion coupled with apical surface expansion, while explicitly accounting for the initial caging effect.

The model simulates a dynamically expanding circular apical domain with area prescribed by experimentally measured growth kinetics (*see SI*). BBs are recruited at a constant rate matching experimental observations (Movie 8), with their initial positions sampled from spatially biased probability distributions that favor peripheral insertion, as observed experimentally (*see SI*). Following insertion, each BB undergoes anomalous diffusion characterized by an exponent *α* randomly sampled from a Gaussian distribution (*µ* = 0.85, *σ* = 0.2). Trajectories were generated using a piecewise fractional Brownian motion approximation based on the Davis-Harte algorithm: the full simulation interval of 150 min was divided into three consecutive segments, each generated with a fixed Hurst exponent *H* = *α/*2, with the exponent reduced by a factor of 3/4 between successive segments and lower bounded by *α* = 0.4. Segment-wise diffusion normalization was imposed using the same reference timescale *t*_ref_ = 30 s.

Critical to reproducing the observed BB spacing was the incorporation of mechanical interactions mediated by traps surrounding each BB as was implicated by the fine-grained model for BB. We model these traps as soft repulsive zones of thickness *r_s_* and stiffness *k_s_*surrounding a hard BB core of radius *r_b_*, with outer pocket radius *r_o_* = *r_b_* + *r_s_*(*see SI*). Let *d* denote the center-to-center distance between two BBs. Repulsion begins when *d <* 2*r_o_*: initially through elastic compression of the soft shells, and subsequently through a stiffer hard-core repulsion once the shells are fully compressed (*d <* 2*r_b_*). This two-stage interaction mechanism naturally prevents BB clustering beyond experimentally observed minimum spacings (∼ 1 *µ*m) while permitting dynamic spatial reorganization. BBs crossing the apical boundary were projected back into the apical domain at a small radial offset from the boundary.

Quantitative comparison of model outputs with experimental data confirmed successful recapitulation of key organizational metrics. The normalized variance of Voronoi tessellation areas decreased progressively over time (Fig. 3i), mirroring the experimental transition from scattered to organized BB distributions as density-dependent mechanical interactions drove spatial regularization. The average nearest-neighbor distance exhibited similar temporal evolution, decreasing steadily before plateauing at values consistent with experimental observations: ∼ 1 *µ*m (Fig. 3j). These results demonstrate that the combination of peripherally biased BB insertion, anomalous diffusion dynamics, and soft-trap mediated mechanical interactions is sufficient to account for the emergent spatial patterning of BBs during apical expansion.

### Progressive formation of an apical actin meshwork facilitates BB redistribution

The theoretical model predicts the presence of soft traps that confine individual BBs which are necessary to reproduce the experimentally observed spacing. This raises a central question: do such trapping structures exist *in vivo* and what molecular components give rise to them? Because BBs are embedded within a dense apical actin network, actin represents a strong candidate to provide the predicted trapping forces. To test this, we examined whether actin organization contributes to BB spatial patterning by acquiring high-resolution images of the apical surfaces of MCCs in embryos injected with Chibby-GFP, followed by fixation and actin (phalloidin) staining. These images revealed pronounced differences in actin organization as a function of apical domain size (Fig. 4a). In smaller apical domains, actin appeared disorganized and lacked a defined structure. In contrast, larger apical domains exhibited progressively more structured actin architectures, culminating in the emergence of a prominent meshwork organization (Fig. 4a).

**Figure 4.**
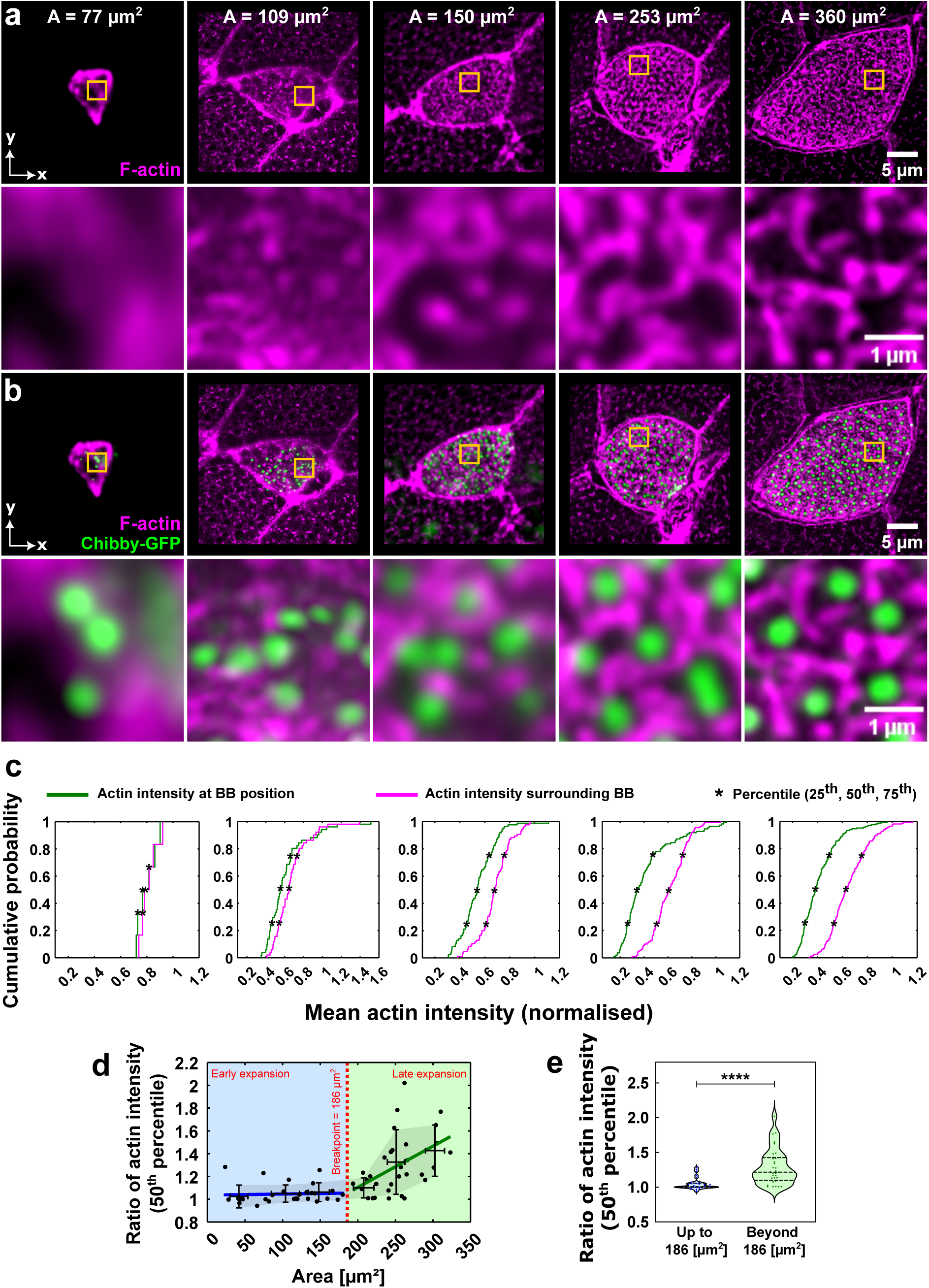
Progressive formation of an apical actin meshwork facilitates BB redistribution. All data from high-resolution Airyscan imaging. (**a–b**) Images of increasing apical domain areas showing the formation of an actin meshwork. (**a**) Actin (Phalloidin). (**b**) Actin and BBs (Chibby-GFP). The second rows in (a) and (b) show zoom-ins of the regions highlighted by orange squares. (**c**) CDFs of actin intensity at the BB positions (green) and in the surrounding regions (pink), corresponding to (a) and (b). Black markers indicate actin intensity at the 25^th^, 50^th^, and 75^th^ percentiles. (**d**) Ratio of actin intensity (surrounding / BB position) at the 50^th^ percentile of CDF, plotted against the apical area. Each dot represents a single cell, and the error bars represent mean ± SD of data averaged over 64 cells across 4 experiments, using 50 µm^2^ bins. The red dotted line marks the breakpoint (186 µm^2^) of the piecewise linear regression while blue and green lines represent the regression fits before and after the breakpoint. The blue and green shaded regions denote data before and after the breakpoint and serve as visual guides for the early (<150 µm^2^) and late (>150 µm^2^) phases of apical expansion. (**e**) Comparison of actin intensity ratios at 50^th^ percentile of CDF before and after the breakpoint in (d); P < 0.0001. The central line represents the median, and the upper and lower lines represent the 75^th^ and 25^th^ percentiles, respectively. Data: 32 cells for each category (≤186 µm^2^) and (>186 µm^2^), from 4 experiments.

During the early stages, when this actin meshwork was absent, BBs were arranged in clustered configurations (Fig. 4b). As the apical domain expanded, actin progressively assembled into a network composed of discrete pocket-like structures that surrounded individual BBs. The formation of interconnected actin pockets into a continuous meshwork across the apical surface correlated with the transition of BBs from a clustered to a more uniform spatial distribution (Fig. 4b).

To further characterize the dynamics of actin meshwork formation and its role in BB patterning, we quantified actin accumulation around the BBs. We compared the mean actin intensity at the BB positions (Fig. S5a) with that in the immediate surrounding regions (Fig. S5b). Cumulative Distribution Function (CDF) plots of actin intensity for these two regions revealed overlapping distributions in small apical domains, which progressively diverged in larger domains (Fig. 4c). This indicated an increase in actin accumulation around the BBs relative to their immediate position. To further quantify these differences, we computed the surrounding-to-BB intensity ratios at the 25^th^ (Fig. S5d), 50^th^ (Fig. 4d), and 75^th^ (Fig. S5e) percentiles of the CDF. Across all percentiles, the ratios increased with apical area, confirming enhanced actin enrichment in BB-surrounding regions in larger domains.

The CDF analysis compares how the full distribution of actin intensities differs between BB positions and surrounding regions, whereas frame averaging summarizes each frame by a single mean intensity value for each region. Thus, the CDF captures distribution-level shifts, while frame averaging provides a per-cell measure of average enrichment and reduces pseudoreplication. Consistent with the CDF based results, the ratio of frame-averaged actin intensity (surrounding / BB position) increased with apical area, further supporting enhanced actin accumulation around BBs in larger domains (Fig. S5f).

Notably, all CDF based measurements (Fig. 4d, S5c, S5d, S5e), as well as the frame-averaged intensity ratio (Fig. S5f), exhibited a common breakpoint around 170-185 µm^2^, beyond which actin enrichment increased markedly. Statistical analysis confirmed significant differences in these metrics before and after this threshold (Fig. 4e, S5g, S5h, S5i).

Importantly, the actin-related breakpoint (∼175 µm^2^) occurs close in developmental time to the transition in BB dynamics previously identified at ∼140 µm^2^ (Fig. 3c, 3e). Considering the continuous expansion of the apical surface over ∼2.5 h, the proximity of these values likely represent the shared underlying developmental transition observed through different quantitative readouts, suggesting a strong mechanistic link between the two processes. In small apical domains lacking an organized actin network (Fig. 4a, 4b), BBs exhibited elongated trajectories and short transition times (Fig. 3a, 3e). As the actin meshwork assembly progressed with increasing apical area, the BB trajectories became more convoluted and transition times increased, reflecting greater confinement (Fig. 3a, 3c, 3e). Thus, the formation of the actin meshwork appears to progressively coordinate the dynamics of the distribution of BBs and ultimately guides the organization of BBs into a defined pocket-like architecture as indicated by our theoretical model.

### Disruption of the apical actin cross-linking impairs BB organization

Actin meshwork formation results from the cross-linking of actin filaments by actin-binding proteins^12,80,81^. To examine how this cross-linked actin network contributes to the distribution and patterning of BB, we knockdown (KD) *α*-actinin-1, a key actin cross-linker^82–84^, using a translation-blocking morpholino oligonucleotide targeting *α*-actinin-1 mRNA. Embryos were injected at the 2–4 cell stage with 15 ng of morpholino, and apical expansion and BB behavior were imaged over 2.5 hours at 30–60 s intervals (Fig. 5a, S6a, Movie 9). Despite *α*-actinin-1 morpholino treatment, BBs were still able to migrate from the basal side to the apical surface (Fig. 5b, Movie 10), and the apical domain expanded (Fig. S6a, Movie 9), with BBs docking to the apical surface during this process (Fig. S6a, Movie 9). However, the apical domain only reached an area of ∼120 µm^2^ at the end of the imaging (Fig. 5c), representing a more than twofold reduction in the expansion rate compared to the wild-type (WT) (Fig. 5d). Furthermore, BB migration and docking were incomplete (Fig. 5b), with approximately half as many BBs reaching the apical domain compared to WT (Fig. 5e, 5f). These results underscore the importance of actin cross-linking for efficient apical expansion and BB docking.

**Figure 5.**
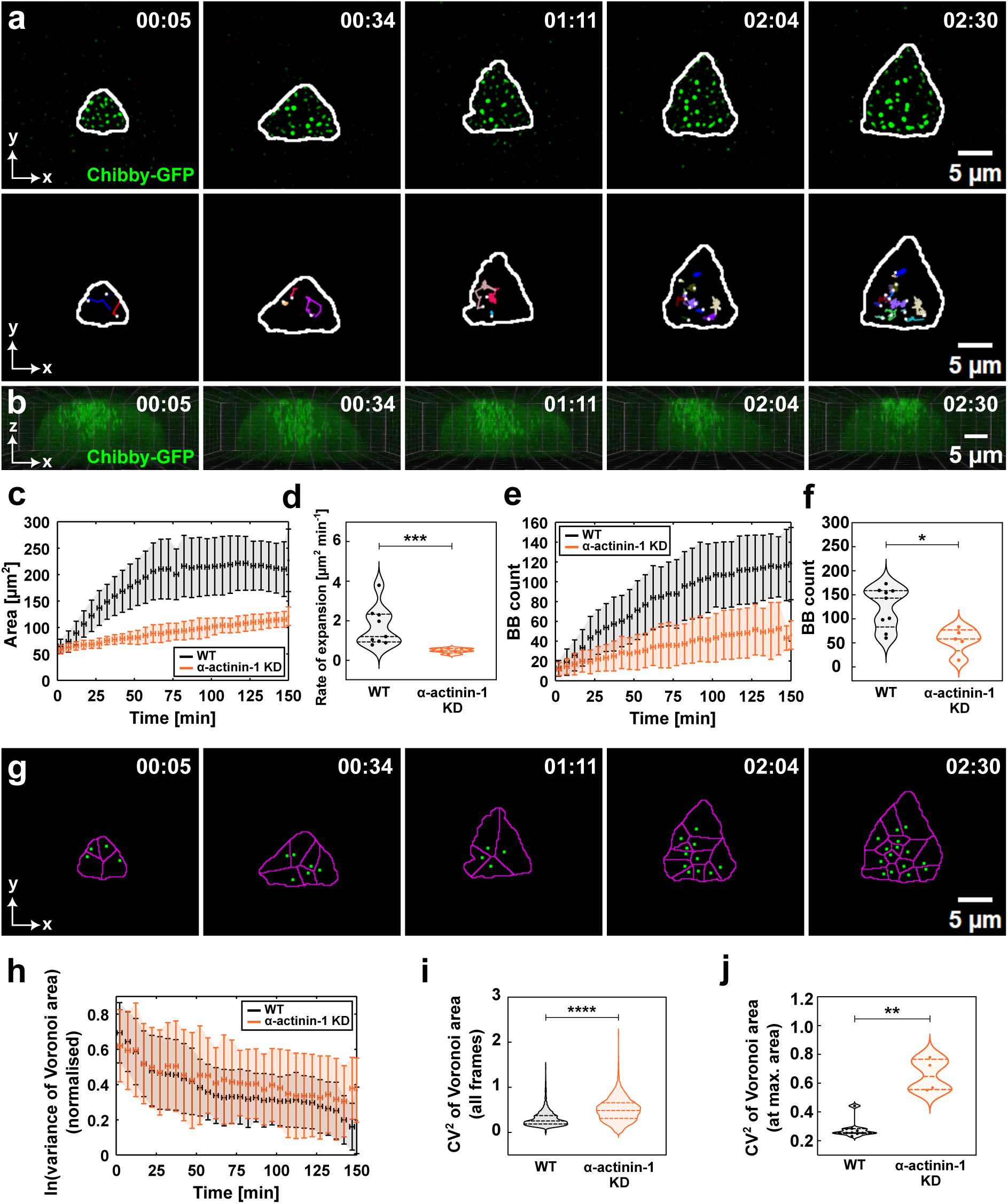
Disruption of apical actin cross-linking impairs BB organization. All data correspond to coarse-timed imaging (frame interval = 30–60 s; total duration = 2.5 h). All plots compare WT (black) and *α*-actinin-1 translation-blocking morpholino (orange) conditions. (**a,b,g**) Time-lapse sequences of an *α*-actinin-1 translation-blocking morpholino injected cell. (**a**) First row: BBs (Chibby-GFP) within the apical domain; second row: corresponding BB trajectories; with the apical periphery outlined by a white contour (see Movies 9 and 11). (**b**) Sequence showing BB ascent from the basal to the apical side of the cell (see Movie 10). (**c**) Apical area plotted against time. (**d**) Comparison of apical expansion rates; exact P = 0.001. (**e**) BB number plotted against time. (**f**) BB number at the maximum apical area reached within 2.5 h; exact P = 0.0028. (**g**) Voronoi tessellations of BBs in an *α*-actinin-1 KD MCC corresponding to (a) (see Movie 12). (**h**) Variance of voronoi tessellation areas plotted against time. Variance values are natural log–transformed (ln) and min–max normalized. (**i-j**) Comparison of variability in Voronoi tessellation areas: (**i**) pooled across all frames (2.5 h); P < 0.0001; (**j**) at the maximum apical area reached within 2.5 h; exact P = 0.0028. Variability is quantified as *CV* ^2^, the squared coefficient of variation (variance divided by the square of the mean). Data in (**c-f**) and (**h-j**) are from 9 WT cells across 5 experiments and from 5 KD cells across 4 experiments. For (**i**), data were pooled from 2611 frames (WT) and 1589 frames (KD). In (**c**), (**e**), and (**h**), error bars represent mean ± SD of binned data (5 min bins). In (**d**), (**f**), (**i**), and (**j**), the central line represents the median, and the upper and lower lines represent the 75^th^ and 25^th^ percentiles, respectively.

Interestingly, in *α*-actinin-1 knockdowns (KDs), apical expansion halted at a maximum of ∼140 µm^2^ (Fig. 5c), which corresponds closely to the area at which actin meshwork formation initiates in WT (Fig. 4), reinforcing the role of *α*-actinin-1 in the assembly of actin networks. Despite these defects, we observed partial preservation of WT characteristics: new BBs still tended to appear near the periphery (Fig. S6b), and BB trajectories exhibited no directional bias, showing equal probability of movement toward or away from the expanding edge (Fig. 5a, S6c, S6d, Movie 11). These observations suggest that, while both the apical expansion and the BB docking were slowed, the coupling between these two processes was maintained. In fact, the density of BB in the apical domain remained comparable to that of WT (Fig. S6e, S6f), indicating that expansion and BB insertion were proportionally delayed.

To determine whether *α*-actinin-1 KD induced delay simply postponed BB organization or fundamentally disrupted it, we examined the apical domains of MCC at later developmental stages (Nieuwkoop and Faber (NF) stage^85^ 26). Although some MCCs eventually reached WT apical sizes, the organization of BBs remained perturbed (Fig. S6g). These defects were confirmed using a splice-blocking morpholino, which yielded similar phenotypes (Fig. S6g, S6h). Together, these findings indicate that although actin cross-linking is not strictly required for apical expansion per se, it is essential for proper BB organization and spatial patterning.

Next, we assessed how loss of actin cross-linking affects the spatial distribution of BBs. The variance of the Voronoi tessellation areas, used here as a proxy of spatial regularity, was increased and more variable in the *α*-actinin-1 KDs throughout expansion (Fig. 5g, 5h, 5i), resulting in a visibly incomplete distribution of BBs (Fig. 5g, 5j, Movie 12). To quantify these effects, we analyzed the minimum and pairwise distance metrics in three populations of BBs: (i) all BBs at each frame (Fig. S7a–Fig. S7d); (ii) newly inserted BBs relative to those from the previous frame (Fig. S7e–Fig. S7h); and (iii) new BBs relative to pre-existing BBs within the same frame (Fig. S7i–Fig. S7l). In all comparisons, the minimum and pairwise distances were consistently lower in *α*-actinin-1 KDs than in WT (Fig. S7a–Fig. S7l), reflecting the visually apparent clumping of BB (Fig. 5a, S6a). These results confirm that actin cross-linking is required to maintain the appropriate spacing and prevent aggregation of BBs during their distribution.

In WT embryos, the apical area serves as a reliable proxy for morphogenetic progression. In contrast, in *α*-actinin-1 KDs, this relationship is disrupted: developmental time (as measured by area) no longer maps linearly onto real time. To evaluate BB organization independently of experimental time, we reanalyzed spatial metrics relative to morphogenetic time, specifically, apical domain area (Fig. S8a–Fig. S8n). Up to the maximum apical area achieved by the *α*-actinin-1 KDs (140 µm^2^), the variance of the tessellation (Fig. S8a, S8b) and the minimum and pairwise BB distances (Fig. S8c, S8e, S8g, S8i, S8k, S8m) showed qualitatively similar trends in both WT and *α*-actinin-1 KD conditions. However, in almost all comparisons, the distributions shifted significantly (Fig. S8d, S8f, S8h, S8j, S8l, S8n), indicating subtle but consistent increases in spatial heterogeneity.

In summary, these findings reveal that actin cross-linking mediated by *α*-actinin-1 is not only required for timely apical expansion and BB recruitment, but is also critical to establishing spatial precision in BB organization. Although some aspects of BB positioning may emerge passively as the apical domain grows, the meshwork formed by cross-linked actin filaments is necessary for refining and stabilizing the final pattern.

### α-actinin-1 perturbation impairs transition in BB dynamics and actin meshwork formation

The coarse-timed data (30-60 s interval) from the *α*-actinin-1 KDs provided insights into the spatial heterogeneity of the BB distribution, arising from disrupted actin cross-linking. These observations were collected over 2.5 hours of experimental time, covering apical areas up to 140 µm^2^, which corresponds to the maximum domain size reached in *α*-actinin-1 KDs during this window and allows direct comparison with the WT condition. However, high-resolution end point imaging revealed that, despite delays, *α*-actinin-1 KDs ultimately reached sizes of the apical domain comparable to those of WT at later developmental stages (Fig. S6g). Thus, the coarse-timed experiments captured the BB distribution and trajectory dynamics only during the early phase of apical expansion in the *α*-actinin-1 KDs.

To overcome this limitation and examine the behavior of the BBs throughout the full range of apical expansion, including the late phase, we employed fine-timed imaging (21 ms interval), which allows sampling of a wide range of apical areas by capturing multiple MCCs over a 10.5 minute window (Fig. 3a). We hypothesized that this approach would reveal additional signatures of disrupted dynamics in *α*-actinin-1 KDs (Fig. 6a, Movie 13).

**Figure 6.**
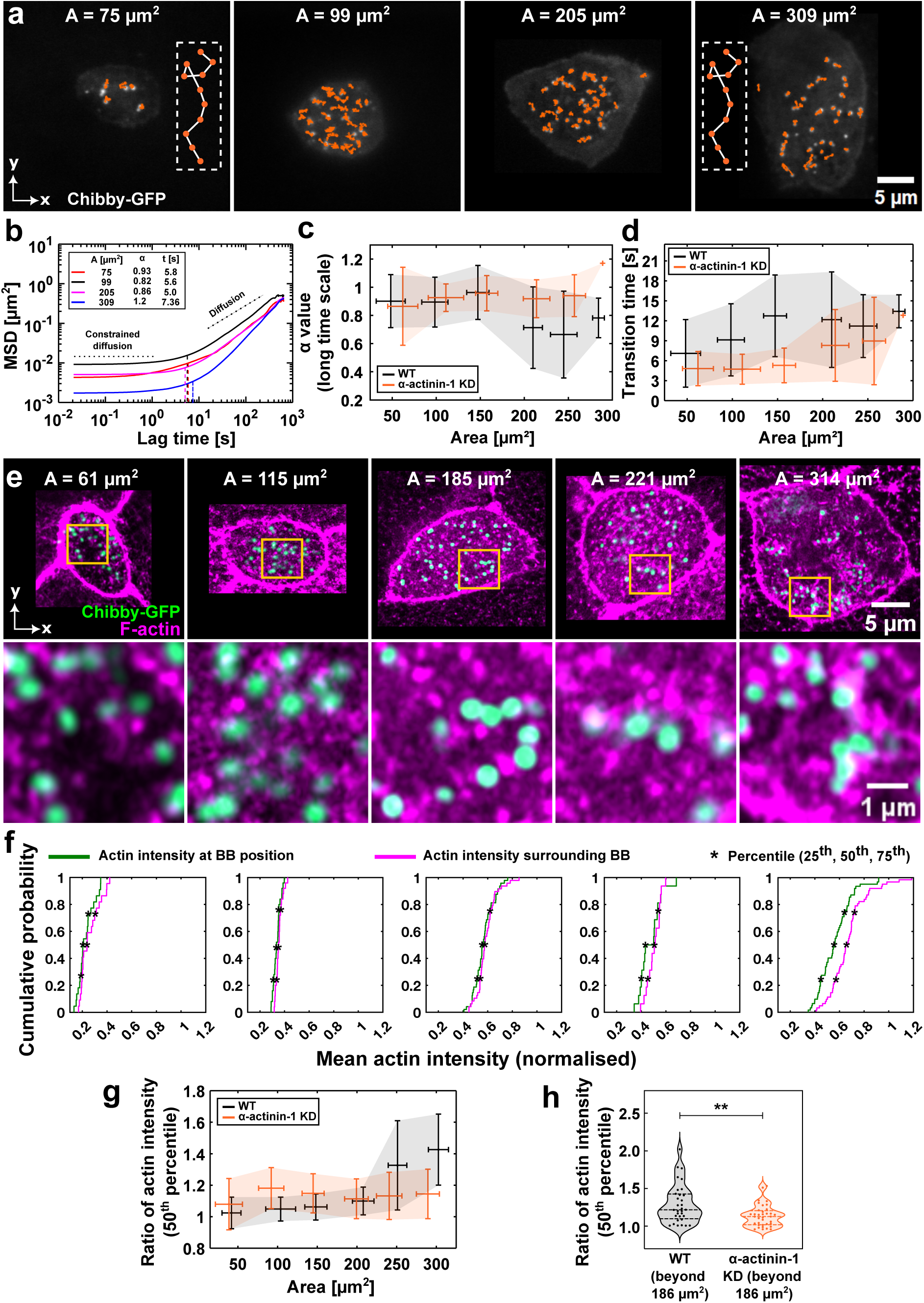
*α*-actinin-1 perturbation impairs transition in BB dynamics and actin meshwork formation. Data in (**a-b**) and (**e-f**) correspond to *α*-actinin-1 translation-blocking morpholino condition; (**c-d**, **g-h**) compare WT (black) and *α*-actinin-1 translation-blocking morpholino (orange) conditions. Data in (**a-d**) correspond to fine-timed imaging (frame interval = 21 ms; 30,000 frames; total duration = 10.5 min). (**a**) Images of four apical domains of different sizes, shown at the last time point of a 10.5 min sequence. BBs (Chibby-GFP) are shown with overlaid trajectories (see Movie 13). The insets in the first and last images show schematics illustrating elongated trajectories in smaller and larger apical domains, respectively. (**b**) MSDs of BBs from the cells in (a), plotted against lag time. Each curve represents the ensemble-averaged MSD of trajectories within a single cell. Vertical dotted lines mark transition times. Reported for each curve are the apical area, *α* value, and transition time. (**c**) Long-time *α* values plotted against apical area. (**d**) Transition times plotted against apical area. In (**c-d**), *α* values and transition times are ensemble averages of all BB trajectories per cell. Error bars represent mean ± SD of averaged data over 4604 trajectories from 100 cells across 17 experiments (WT) and 2029 trajectories from 46 cells across 5 experiments (KD), using 50 µm^2^ bins. Data in (**e-h**) correspond to high-resolution Airyscan imaging. (**e**) Images of apical domains of increasing size showing the lack of actin meshwork formation. Top row: actin (Phalloidin) and BBs (Chibby-GFP). Bottom row: zoom of regions highlighted in orange squares in the top row. (**f**) CDFs of actin intensity at BB positions (green) and surrounding regions (pink), corresponding to (e). Black markers indicate actin intensity at the 25^th^, 50^th^, and 75^th^ percentiles. (**g**) Ratio of actin intensity (surrounding / BB position) at the 50^th^ percentile of CDF, plotted against the apical area. Error bars represent mean ± SD of averaged data over 64 cells across 4 experiments (WT) and 70 cells across 3 experiments (KD), using 50 µm^2^ bins. (**h**) Comparison of actin intensity ratios at 50^th^ percentile of CDF between WT and KD, beyond 186 µm^2^; (exact P = 0.0059). Data from 32 cells across 4 experiments (WT) and 36 cells across 3 experiments (KD). The central line represents the median, and the upper and lower lines represent the 75^th^ and 25^th^ percentiles, respectively.

We first analyzed the averaged MSDs of BBs across apical areas ranging from 50 to 300 µm^2^ (Fig. 6b). In *α*-actinin-1 KDs, MSD plots showed constrained behavior (*α* ≪ 1) at short timescales, followed by a transition to primarily diffusive behavior (*α* ≈ 1) at longer timescales (Fig. 6b). This contrasts with WT trajectories, where BBs exhibit anomalous diffusion on long timescales (Fig. 3b), reflecting evolving BB motions due to progressive actin meshwork formation. In *α*-actinin-1 KDs, both short-timescale *α* values and diffusion strength remained constant across the range of apical areas (Fig. S9a, S9b), and the long-timescale *α* values similarly showed no area-dependent trend (Fig. 6c, S9e), unlike in the WT, where these values decreased progressively with expansion (Fig. 3c). The absence of any breakpoint or shift in dynamics indicates that BB trajectories in *α*-actinin-1 KDs are not remodeled during expansion. This is supported by the relatively constant end-to-end distances (Fig. S9c) and tortuosity values (Fig. S9d), consistent with the elongated and unconstrained trajectories observed in *α*-actinin-1 KDs (Fig. 6a).

We next focused on the transition in the *α* values between short and long timescales, which in WT reflects the shift from trapped to freely diffusing states. In *α*-actinin-1 KDs, the BB trajectories followed a similar trend: low *α* values at short timescales and diffusive values at longer timescales (Fig. 6b). This transition also became more pronounced with increasing apical area (Fig. 6d, S9f), as observed in WT (Fig. 3e). However, the transition in *α*-actinin-1 KDs was markedly delayed. In WT, transition times increased to ∼12.5 s at ∼150 µm^2^ (Fig. 3e), while in *α*-actinin-1 KDs, the same transition time was only reached at ∼300 µm^2^ (Fig. 6d). This indicates that BBs in *α*-actinin-1 KDs are released from confinement more rapidly and remain in a more dynamic state even in later stages of expansion, reflecting a failure of spatial refinement that normally occurs through actin cross-linking.

Altogether, these results show that in *α*-actinin-1 KDs, the BB trajectories (Fig. 6c) and the trap-release dynamics (Fig. 6d) remain largely unchanged in the apical area range. This suggests that *α*-actinin-1 mediated actin cross-linking is essential to regulate BB motion during apical expansion. In WT, this cross-linking begins around ∼170 µm^2^ (Fig. 4, Fig. S5), falling within the same apical area range as the transition in BB behavior. In *α*-actinin-1 KDs, due to the lack of cross-linking, BBs retain diffusive trajectories and exhibit only brief trapping events even in large apical domains (Fig. 6c, 6d). The elongated, unconstrained trajectories observed in the late stages (Fig. 6a, Movie 13) further support this. The delayed transition times in *α*-actinin-1 KDs reflect an inability to stably confine BBs, thereby impairing their progressive spatial organization. These findings highlight the importance of the gradual buildup of actin cross-linking in refining BB dynamics and achieving proper patterning. Importantly, while the number of BBs is known to scale with the apical area^57,58^, our *α*-actinin-1 KD experiments reveal that the BB docking also critically depends on the actin architecture. Thus actin cross-linking introduces an additional layer of regulation in the relationship between the apical area and the BB number.

To confirm the impact of *α*-actinin-1 KD treatment on actin organization, we performed high-resolution imaging of apical domains injected with Chibby-GFP and stained with phalloidin to visualize the actin network (Fig. S9g). Across the full range of apical areas, *α*-actinin-1 KDs displayed disorganized actin, lacking the meshwork architecture and characteristic actin pockets observed in WT cells, with BBs correspondingly arranged in clumped configurations (Fig. 6e). Quantitative analysis supported these observations: CDFs of actin intensity at BB positions and in their surrounding regions failed to diverge in large apical domains (Fig. 6f), in contrast to the behavior observed in WT cells, indicating an absence of actin enrichment around BBs. Consistently, the ratio of actin intensities (surrounding / BB position) at the 50^th^ percentile (Fig. 6g, 6h) did not increase with apical area in *α*-actinin-1 KDs.

Furthermore, frame-averaged actin intensities at BB positions and in the surrounding regions were consistently lower in *α*-actinin-1 KDs throughout apical expansion compared to WT (Fig. S9h, S9k, S9i, S9l), indicating a global reduction in actin accumulation in these regions. Accordingly, the ratio of frame-averaged actin intensity (surrounding / BB position) showed no increase in *α*-actinin-1 KDs (Fig. S9j, S9m), whereas in WT cells this ratio began to rise around 178 µm^2^ (Fig. S5f, S9j). Together, these results confirm the absence of actin meshwork formation in *α*-actinin-1 KDs and establish *α*-actinin-1 as a critical factor in actin meshwork assembly in MCCs.

Finally, we note that the absence of a breakpoint in actin intensity metrics (Fig. 6g, S9j) mirrors the lack of transition in the BB dynamics (Fig. 6c, 6d). Together, these data demonstrate that in the absence of actin cross-linking, the actin meshwork fails to develop, preventing BB confinement and resulting in clumped, unpatterned distributions. These findings establish *α*-actinin-1 dependent actin cross-linking as a key regulator of BB dynamics and spatial patterning in MCCs.

### Heterogeneous distribution of the apical actin network leads to BB clumping

The *α*-actinin-1 KDs showed clear changes in BB dynamics and patterning, underscoring the importance of actin cross-linking in this process. To further investigate how BB disorganization manifests under these conditions, we quantified the spatial heterogeneity of BB distributions. Coarse-timed imaging was used to capture the chronological evolution of BB disorganization during apical expansion. Visually, two distinct clustering patterns were observed: (i) BBs aggregating into a single large cluster localized to one end of the apical domain (global cluster), or (ii) BBs forming multiple smaller clusters scattered across the surface (local cluster) (Fig. 7a).

**Figure 7.**
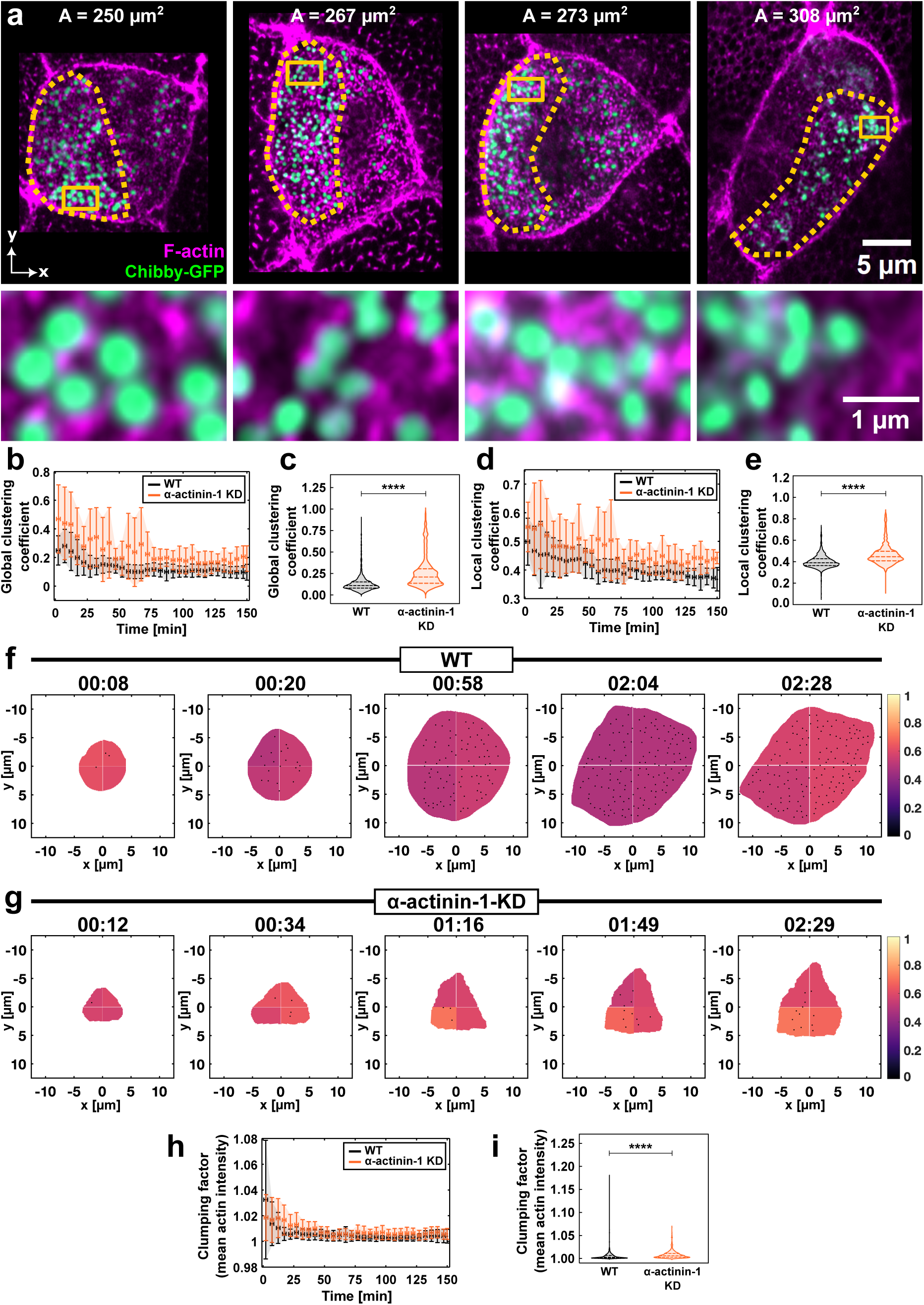
Heterogeneous distribution of the apical actin network leads to BB clumping. All data from coarse-timed imaging (frame interval = 30–60 s; total duration = 2.5 h). Data in (**b-e**) and (**h-i**) compare WT (black) and *α*-actinin-1 translation-blocking morpholino (orange) conditions. (**a**) Images showing the global and local clustering in *α*-actinin-1 translation-blocking morpholino condition. Areas outlined by dotted orange lines indicate global clustering, whereas the zoom-ins of regions outlined by solid lines show local clustering of BBs. (**b, d**) Time evolution of the global (**b**) and local (**d**) clustering coefficients from experiments. (**c, e**) Comparisons of global (**c**) and local (**e**) clustering coefficients between WT and KD. (**f-g**) Time-lapse sequences of apical domains discretized into four quadrants and color-coded for mean actin intensity: (**f**) WT; (**g**) KD (see Movie 16). (**h**) Time evolution of the clumping factor for actin. (**i**) Comparison of clumping factors between WT and KD for actin. In (**b**), (**d**) and (**h**)), error bars represent mean ± SD of binned data (5 min bins). Statistics for comparisons: (**c**) 2,554 frames (WT) vs. 1,455 frames (KD), P < 0.0001; (**e**) 2,552 frames (WT) vs. 1,455 frames (KD), P < 0.0001; (**i**) 1,670 frames (WT) vs. 1,182 frames (KD), P < 0.0001. In (**c**), (**e**) and (**i**), the central line represents the median, and the upper and lower lines represent the 75^th^ and 25^th^ percentiles, respectively. Data in (**b-e**) and (**h-i**)) were obtained from 9 cells across 5 experiments (WT) and 5 cells across 4 experiments (KD).

To quantify heterogeneity in the BB distribution, we employed two clustering coefficients: a global clustering coefficient that incorporates both angular and radial variation and a local clustering coefficient based on the ratio of nearest-neighbor distance to the mean inter-BB distance (*see methods section 7.3.12 in SI*). These metrics provided continuous and unbiased quantification of BB clustering throughout the apical domain, taking into account orientation and position. In WT cells, the global clustering coefficient (Fig. 7b) and the local clustering coefficient (Fig. 7d) followed the expected pattern, consistent with other spatial organization metrics (Fig. 1i, 1j): initially elevated, reflecting early-stage disorganization, followed by a gradual decrease as the BBs became more uniformly distributed.

Interestingly, *α*-actinin-1 KDs also showed a decreasing trend in both coefficients (Fig. 7b, 7d), suggesting a trend towards spatial organization. However, the variance of these metrics was greater and their absolute values were significantly elevated compared to WT (Fig. 7c, 7e), indicating persistent disorganization.

To investigate the mechanism underlying the observed BB clustering, we turned to our theoretical model. For simulations replicating *α*-actinin-1 KD, three key modifications were implemented to capture experimental perturbations: (i) BBs exhibited a strong spatial bias, with a higher probability of appearance within one region of the apical area compared to the rest of the domain (*see SI*); (ii) diffusion strength was reduced by a factor of 5 in non-preferred quadrants, creating localized confinement that restricts redistribution; and (iii) actin pocket thickness was reduced, weakening repulsive interactions and permitting closer BB approach (Movie 14). The temporal analysis of simulated clustering metrics revealed dynamics consistent with experimental observations (Fig. S10a, S10b). Both global and local clustering coefficients were initially elevated for wild-type (WT) and knockdown (KD) simulations, reflecting early-stage spatial disorganization when BB numbers were low. As time progressed, the global clustering coefficient decreased toward zero in WT simulations as BBs became scattered across the apical surface through stochastic peripheral appearance and anomalous diffusion. In the KD simulations, however, the global clustering coefficient remained saturated at relatively higher values even at later times, as BBs remained confined within the preferred quadrant. Similarly, local clustering decreased gradually in WT simulations but remained elevated in KD simulations due to reduced actin pocket sizes that permitted closer BB approach.

Notably, inducing clustering *in silico* required simultaneous tuning of both parameters, i.e., BB repulsion distance and spatial bias in BB appearance; manipulating either alone was insufficient to drive clustering. This suggests that BB clustering in the experimental *α*-actinin-1 KD arises from a combination of biased BB emergence and a lack of repulsive interactions, likely due to disrupted actin cross-linking, which normally enforces BB spacing. However, the origin of the preferential appearance could not be readily explained. Given that BB docking requires actin, we hypothesized that apical actin distribution may be involved in spatially biased BB docking.

Although clustering coefficients effectively capture heterogeneity in discrete BB distributions, they are not optimal for comparing spatially continuous signals such as actin intensity, which serves as a proxy for the underlying actin distribution. To address this, we developed a quadrant-based clumping factor that quantifies spatial heterogeneity for both discrete (BB positions) and continuous (actin intensity) distributions. The apical domain is divided into four equal angular sectors, and for each time frame, the number of BBs (or mean actin intensity) in each quadrant is determined. The clumping factor is then calculated as the normalized variance of these quadrant populations (*see methods section 7.3.13 in SI*). If BBs are uniformly distributed across all quadrants, the clumping factor approaches 1; if all BBs are confined within a single quadrant, it approaches 4. This metric enables direct comparison of spatial heterogeneity between BB density and actin intensity distributions within a unified analytical framework (Fig. S10c, S10d, Movie 15).

In the experimental data, clumping factors for both BB density (Fig. S10e, Movie 16) and actin intensity (Fig. 7f, Movie 16) followed the same trend observed in clustering coefficients: initially elevated, then declining towards 1 over time (Fig. 7h, S10g). This trend indicates a transition from spatial heterogeneity to homogeneity during normal apical expansion. In contrast, although the *α*-actinin-1 KDs showed a similar downward trend (Fig. 7h, S10g), their clumping factors remained significantly higher than those of WT, for both BBs (Fig. S10f, S10h, Movie 16) and actin (Fig. 7g, 7i, Movie 16), indicating persistent heterogeneity in both components. Consistent with the experimental observations, BB simulations incorporating preferential appearance and reduced repulsion exhibited higher clumping factors compared to simulations with regularly spaced BBs (Fig. S10i).

Together, these analyses demonstrate that the *α*-actinin-1 KDs consistently exhibit elevated spatial heterogeneity in the BB distribution, primarily in the form of BB clustering. These clusters tend to colocalize with regions of increased actin intensity, suggesting that BB disorganization reflects and is likely driven by the underlying heterogeneity of actin. These findings reinforce the critical role of actin cross-linking in the regulation of both BB dynamics and spatial patterning during apical expansion.

## Discussion

The actin cytoskeleton is a fundamental component and master regulator of cellular architecture and related functions^13,77,86^. It can rapidly reorganize by assembling^87^, disassembling^88^, and forming higher-order structures such as filamentous networks^12,82^ and bundles^89^. These structures propagate forces across scales^90,91^, facilitating processes ranging from intracellular transport^92^ and organelle positioning in the crowded cytoplasmic environment^93,94^, to cell migration^95^ and tissue morphogenesis^96^. For example, in plant cells, the actin cytoskeleton efficiently transports Golgi bodies, and their movement can be predicted from the global organization of the actin network ^97^.

Our quantitative analysis of the MCC apical domains in *X. laevis* embryos reveals that the dynamic organization of the apical actin meshwork plays a decisive role in the emergent mobility and patterning of BBs. Importantly, we capture changes in meshwork formation that are acutely reflected in BB dynamics and patterning. This reinforces the idea that the dynamics of the BBs are tailored to their final patterning, which is rigorously orchestrated by actin remodeling. These findings align with the known role of actin dynamics in intracellular organization and significantly extend this established paradigm.

Previous studies have shown a correlative relationship between the size of the apical area and the BB count^57,58^. Here, we demonstrate that this relationship is dynamic, and the BB count increases linearly with area. Interestingly, this correlation does not break down in the *α*-actinin-1 KDs. Although the rate of expansion and BB docking is slower compared to that of the WT, the BB density is still preserved. This argues for a mechanism involving apical actin, which balances the expansion of the apical domain with the number of BBs docked. We propose that cross-linking-based meshwork formation maintains this balance, patterning the BBs while simultaneously providing mechanical stability to the expanding apical domain. Apical expansion then creates a new space that allows the recruitment of BBs. Interestingly, while cross-linking coordinates apical expansion and BB patterning, it also keeps these processes largely independent. If the processes were interdependent, then the apical expansion should already passively distribute the BBs. In contrast, the movement of BB trajectories in the direction opposing expansion argues for a decoupling mechanism between apical expansion and BB distribution. The correlation between actin remodeling and the shift in BB dynamics from diffusive (or even superdiffusive) to subdiffusive behavior implies that actin is the decoupling element. While actin polymerization pushes the boundary of the apical domain, the cross-linked actin meshwork presumably stabilizes the expanded area and simultaneously patterns the BBs. Thus, the link between apical expansion and BB patterning appears to be mediated through actin remodeling, although the direction of causality remains unclear.

The apical actin appears to take control of the BBs as soon as they are incorporated into the apical domain. In WT conditions, new BBs emerge near the periphery, close to the newly extended area, but then start migrating in a tortuous fashion until they settle among their pre-existing neighbors. This in-plane trafficking reflects the local activity of actin, which leads to mechanical push-pull within the apical cortex. In contrast, in *α*-actinin-1 KDs with disrupted cross-linking, although new BBs appear peripherally, they wander randomly and ultimately accumulate into clusters rather than forming a structured arrangement. This is further confirmed by our minimum and pairwise distance metrics and clustering analysis, which show that actin integrity in the WT maintains uniform spacing. *α*-actinin-1 KDs, on the other hand, exhibit higher clustering coefficients and greater heterogeneity in distance metrics. Although *α*-actinin-1 KDs consistently show higher levels of heterogeneity compared to WT, a baseline level of heterogeneity is always present in the WT. This likely reflects intrinsic biological noise in the system. In this context, the actin meshwork can be viewed as a regulatory mechanism that refines the BB pattern against this inherent background noise. Disruption of actin cross-linking in *α*-actinin-1 KDs compromises this regulatory mechanism, allowing intrinsic noise to become amplified and leading to disorganized spatial patterning of BB. Thus, actin cross-linking likely facilitates the coordinated movement of BBs into their final patterned positions. This is consistent with the current understanding of how actin networks and cross-linkers give rise to dynamic patterns across scales in diverse biological contexts, as well as their functional relevance^14,91,98^.

Previous literature has suggested that BBs may bring actin polymerization factors, such as formins and Arp2/3, to the apical domain, thus promoting cross-linking^99,100^. However, blocking BB formation and apical docking does not impair the initial phase of apical expansion^57^. These studies focused on endpoint effects, not time-evolution dynamics. Therefore, BBs might be influencing the apical expansion by helping the apical domain reach the critical amounts of actin-related proteins within a defined time frame for the formation of the meshwork. The results of the *α*-actinin-1 KD in our study clearly show that the lack of actin cross-linking prevents the BB from docking to the apical domain, with no apparent effect on BB synthesis or migration. Specifically, the arrest of the apical expansion at ∼120 µm^2^ points to a mechanism other than cross-linking that may drive the expansion up to this range of the area. This is likely actin polymerization, which is then taken over by branching and cross-linking. If identified, this switch in branching and cross-linking could be considered a key developmental time point in MCC apical expansion. Thus, the results of this study further solidify the idea that the actin network provides both mechanical support and the dynamic rearrangements necessary for precise dynamics and organization of subcellular organelles, particularly in specialized cells such as MCCs that undergo extensive morphogenetic changes.

Perturbations in actin cross-linking significantly impair the spatiotemporal patterning of BBs, leading to observable defects in the expansion of the apical area, the count of BBs, the variance of the tessellation areas, and the minimum and pairwise distance metrics of different BB populations. These widespread disruptions highlight that actin cross-linking is not a passive process but an active organizer that enables the apical domain to facilitate BB arrangement. The fact that BB density, peripheral appearance, and trajectory direction remain similar to WT, while other parameters are disrupted, suggests a selective impairment of BB organization rather than a complete abolition of BB biogenesis or basal-to-apical migration. This implies that while initial BB appearance and general directional tendencies are preserved, the fine-tuning of their spatial relationships and overall patterning is highly dependent on an intact and cross-linked actin meshwork.

MSD analysis is a powerful tool for characterizing the nature of particle motion in complex biological environments^5,101^. When a particle exhibits standard diffusion with Brownian motion, MSD scales linearly with time and the power-law exponent (*α*) of the MSD-time relationship equals 1. However, deviations from this behavior indicate anomalous diffusion, including subdiffusion (*α <* 1), superdiffusion (*α >* 1), and ballistic motion (*α* ≈ 2)^5,101^. Our findings from MSD analysis provide critical insight into BB dynamics and their strong correlation with actin meshwork formation. The transition of cortical actin into a cross-linked meshwork occurs concurrently with a shift in the BB dynamics, from freely diffusing or superdiffusive motion in small apical domains to confined, subdiffusive motion in expanded domains. This suggests that the actin meshwork lattices of larger domains effectively cage the BBs, restricting long-range movement. In contrast, when actin cross-linking is disrupted in *α*-actinin-1 KDs, the meshwork fails to form, and BBs remain relatively unconstrained and diffusive throughout. This result is in line with studies that demonstrate that WDR5^62^ and Filamin-A^63^ contribute to an F-actin lattice essential for uniform BB spacing. We show that, in the absence of this caging effect, BBs exhibit clumping and enhanced mobility. Further, our data directly link actin meshwork formation to BB single-particle dynamics: the meshwork imposes subdiffusive constraints, while in its absence, BBs behave in a nearly Brownian fashion. Thus, while coarse-timed experiments highlighted the role of cross-linking in the organization of BBs, it is the fine-timed data that reveal the underlying mechanism. These findings suggest that actin remodeling progressively primes the BB dynamics to achieve a final ordered state.

This conclusion is supported by a large body of biophysical literature on diffusion in complex media. In polymer networks, particles often exhibit anomalous behavior due to physical obstacles and active interactions^4–8^. Depending on the dynamic state of the environment and the nature of particle–environment interactions, particles can exhibit complex combinations of anomalous and diffusive behaviors, separated by a characteristic timescale^7,17,102^. These types of behavior, including transient caging, have been observed in various systems, including nonbiological particles in colloidal fluids and glasses^103^, granular materials under cyclic shear^104^ or vertical shaking^105^, as well as biological contexts such as lipid bilayers^106,107^ and F-actin networks^108^.

Consistent with these findings, our MSD analysis of WT MCCs shows a transition from constrained to diffusive, or even superdiffusive, motion in BB trajectories. These transitions occur over a characteristic timescale (transition time), which increases from approximately 5 s to 15 s as the apical area expands. This observed transition time aligns with the timescale of actin remodeling reported in the literature^75–77^. The increase in transition time correlates with the formation of the meshwork, highlighting the role of the actin network in tuning the behavior of the BB trajectory. In contrast, in *α*-actinin-1 KDs, this transition is severely delayed, suggesting that BBs behave as if in a more fluid, unconstrained environment. This observation agrees with studies in reconstituted cytoskeletal systems, where actin-microtubule cross-linked networks lead to subdiffusive motion of embedded tracer particles^18^. Thus, in WT, the actin meshwork likely cages BBs during late apical expansion, reducing their mobility and resulting in subdiffusive motion. In its absence, as in *α*-actinin-1 KDs, BBs diffuse freely. The delay in the transition time in the *α*-actinin-1 KDs indicates that, without reinforcement via cross-linking, the actin network never matures into a phase capable of constraining the BB motion.

While transitions between anomalous and diffusive behavior have been reported, to our knowledge, this is the first demonstration of a transition from one anomalous regime to a combination of anomalous and diffusive behaviors, spanning subdiffusive and superdiffusive motion, in the context of development. This suggests that the physics governing subcellular dynamics in developmental processes, such as MCC apical expansion in *X. laevis*, may be more complex than currently appreciated. It also emphasizes the need for careful consideration in choosing appropriate models for intracellular trajectory behavior^79^.

Recent advances in active gel theory provide a compelling framework for understanding the role of cytoskeletal remodeling in generating subcellular patterns, such as the BB organization described in this work^12–16^. In active gels such as the actin cytoskeleton, cross-linkers transiently bind and unbind to polymerizing actin filaments, tuning network connectivity and modulating viscoelastic properties^12,84,109–115^. Our findings are consistent with this framework: the *α*-actinin-1 based meshwork emerges during apical expansion and constrains BB mobility. This suggests that the meshwork acts as a spatially extended mechanical scaffold, regulating the positioning of the BB on the apical surface.

The concurrent shift in actin meshwork formation and BB dynamics, from fast, diffusive behavior to slower, subdiffusive motion, suggests that the surrounding cytoskeletal environment becomes increasingly confining. Notably, both shifts occur around similar expansion range: 140 µm^2^ for BB dynamics and 170 µm^2^ for meshwork formation. These observations suggest a possible percolation transition ^12,84,109,113–115^ around this area range, where the meshwork becomes sufficiently connected across the apical surface to behave as a constraining solid-like medium over timescales relevant for BB motion. This interpretation is consistent with previous work^52^ that describes BBs as particles embedded in a viscoelastic medium. Our results further specify that this viscoelastic nature arises from actin polymerization, which drives activity, and *α*-actinin-1 cross-linking, which provides mechanical integrity.

Thus, the apical actin network in MCCs is not only a passive stabilizer of apical domain expansion but also functions as a viscoelastic gel whose mechanical properties, regulated by cross-linkers such as *α*-actinin-1, drive the emergent spatial organization of BBs. These findings are in line with observations in other systems: for example, apical actin organization is critical for anchoring hundreds of cilia in human airways^59^; perturbations of WDR5^62^ and filamin^63^ disrupt BB spacing in *Xenopus*; and actin-modulating drugs and actin-associated factors affect BB distribution and docking in *Xenopus* and zebrafish MCCs, respectively^54,116^. Together with our data, these studies suggest a conserved mechanism by which actin meshwork dynamics actively dictate BB organization.

In conclusion, through a combination of imaging, quantitative analysis and perturbation studies, we establish the cytoskeletal organization of actin as a central driver of BB spatial patterning in *X. laevis* MCCs. We anticipate that identifying this mechanistic link between cytoskeletal mechanics and BB dynamics will advance our understanding of how subcellular architecture emerges during development and how its dysregulation contributes to disease states.

## Methods

The sections below describe the experimental procedures, and all materials used in these experiments are listed in tabular form in the List of Materials section in the Supplementary information.

### 1 *Xenopus laevis* embryo preparation and manipulation

#### 1.1 Husbandry

Adult *Xenopus laevis* were obtained from NASCO (Fort Atkinson, WI, USA) or Xenopus 1 (Dexter, MI, USA). Animals were maintained in a centralized facility in accordance with guidelines provided by the National Xenopus Resource (Marine Biological Laboratory, Woods Hole, MA, USA), the European Xenopus Resource Center (University of Portsmouth, UK), and Xenbase (http://www.xenbase.org). All procedures were reviewed and approved by the Danish National Animal Ethics Committee (permit no. 2017-15-0201-01237).

#### 1.2 Embryo preparations

Ovulation in adult female *Xenopus laevis* was induced by injection of human chorionic gonadotropin (hCG; Chorulon; 500 IU per animal) on the day prior to experiments. Approximately 16 h later, eggs were collected by gentle abdominal squeezing of hormonally primed females. Eggs were fertilized in 1/3× Modified Ringer’s (MR) solution (33 mM NaCl, 0.6 mM KCl, 0.67 mM CaCl_2_, 0.33 mM MgCl_2_; pH 7.6) by addition of a small piece of crushed testis obtained from a sacrificed adult male. After ∼2 h at 20 °C, fertilized embryos were dejellied by incubation in 2.5–3% (w/v) Cysteine solution (pH 7.8) for 8 min, followed by several rinses in 1/3× MR. Morphologically healthy embryos were then selected and transferred to fresh 1/3× MR for subsequent manipulations.

#### 1.3 Microinjections of plasmids and morpholinos

For targeted fluorescent labeling of BBs and actin in multiciliated cells (MCCs), embryos were microinjected with the following plasmids driven by the *α*-tubulin promoter: Chibby-GFP/RFP and LifeAct-GFP/RFP, at 5 ng/µL per blastomere. Microinjections were performed into the ventral blastomeres at the (Nieuwkoop and Faber (NF)^85^) 2-cell or 4-cell stage, using either 1/3× MR solution or 2% Ficoll in 1/3× MR. For *α*-actinin-1 knockdown experiments, morpholino doses were carefully titrated to 15 ng per blastomere. Injection of morpholino at the 2–4-cell stage caused basal bodies to migrate from the basal to apical side, expansion of the apical domain, and docking of basal bodies into the expanding apical domain, while formation of the actin meshwork was impaired. Translation-blocking (ATG) or splice-blocking *α*-actinin-1 morpholinos (Gene Tools; morpholino sequences are listed in Table 1) were co-injected with Chibby-GFP/RFP and LifeAct-GFP/RFP constructs for knockdown experiments. Prior to injection, the morpholino solution was heated to 90 °C for 10 min, centrifuged, and the supernatant was subsequently mixed with the Chibby- and LifeAct-based constructs to obtain the desired concentrations. After injection, embryos were incubated at 18-23 °C in 1/3x MR until the desired stage (20-26) for live imaging or fixation.

#### 1.4 Immunostaining for high-resolution imaging

For high-resolution Airyscan imaging of the actin meshwork, embryos were injected at the 2–4 cell stage with the *α*-tubulin driven Chibby-GFP plasmid and fixed at the desired developmental stage for phalloidin staining. For knockdown experiments, *α*-actinin-1 morpholino was co-injected with Chibby-GFP. Embryos were fixed between stages 22–26 to visualize actin meshwork formation across different BB distributions and apical domain sizes. At the desired stage, vitelline membranes were carefully removed, and embryos were rinsed once in MEMFA (0.1 M 3-(N-morpholino)propanesulfonic acid (MOPS), pH 7.4; 2 mM Ethylene glycol-bis(*β*-aminoethyl ether)-N,N,*N ^′^*,*N ^′^*-tetraacetic acid (EGTA); 1 mM MgSO_4_; 3.7% formaldehyde), followed by fixation in fresh MEMFA on a rotator either for 2 h at room temperature (∼22 °C) or overnight at 4 °C. Following fixation, embryos were washed three times in Tris-buffered saline containing Triton X-100 (TBST; TBS: 155 mM NaCl, 10 mM Tris-HCl, pH 7.4, supplemented with 0.1% Triton X-100) for 5 min each. Blocking was performed by incubation in blocking buffer (10% fetal bovine serum (FBS) and 5% dimethyl sulfoxide (DMSO) in TBS) for 1 h, followed by washing in TBST for 1 h on a rotator. Embryos were then incubated with fluorescent phalloidin (555, 568, or 647; 1:500 dilution) for 2 h at room temperature or overnight at 4 °C. After staining, embryos were washed three times in TBST over 30 min and stored in TBS at 4 °C for up to 10 days. All fixed embryos were imaged within 10 days of staining.

#### 1.5 Embryo mounting for imaging

##### 1.5.1 Live imaging

Embryos at the desired developmental stage were manually removed from their vitelline membranes and briefly rinsed in drops (∼70 µL) of 1.5–2% low– or ultra-low–melting-point agarose preheated to 65 °C. This rinsing step was repeated twice using fresh agarose drops to dilute residual MR solution coming from the embryo transfer. A fresh drop (∼70 µL) of molten agarose was placed onto a coverslip mounted in the imaging chamber (Attofluor Cell Chamber for microscopy). Rinsed embryos (5–7 per drop) were transferred into this agarose drop and their orientations were adjusted so that the flank of the embryos faced the coverslip. Then excess agarose surrounding the embryos was carefully aspirated to slightly flatten the embryo surface against the coverslip. This increased the accessible surface area for imaging without excessive flattening. The agarose was allowed to set at 13 °C for 3 min. A small volume (∼20 µL) of agarose was then added to the top and sides of the set agarose to reinforce the mount and allowed to solidify for an additional 3 min at 13 °C. This reinforcement step was repeated twice before gently filling the imaging chamber with fresh 1/3× MR solution. All agarose mounting steps were performed at a reduced ambient room temperature (∼20 °C). Further, agarose was dispensed using pipettes with tips cooled by briefly holding them in hand prior to contact with embryos. These precautions minimized heat-induced damage to the embryonic surface. Agarose-mounted embryos were imaged for up to 5 h, after which a fresh batch of embryos was mounted.

##### 1.5.2 Fixed imaging

Phalloidin-stained embryos stored in TBS were placed directly onto a coverslip mounted in the imaging chamber. Excess TBS was gently aspirated to allow the embryos to flatten against the coverslip, maximizing the available surface area for imaging. A small coverslip fragment, with silicone grease applied to its four corners, was gently pressed from the top of the embryos to stabilize their position. The imaging chamber was then carefully flooded with TBS and transferred for imaging.

### 2 Microscopy

#### 2.1 Imaging setups

##### 2.1.1 Confocal microscopy

All coarse-timed imaging were performed using either a Zeiss LSM880 or a Leica Stellaris line-scanning confocal microscope. Detailed acquisition specifications for each system are provided below. *Zeiss LSM880*: Imaging was performed using 1–3% laser power from Argon 25 mW (488 nm, 514 nm), diode-pumped solid-state (DPSS) 10 mW (561 nm), and helium–neon (HeNe) 5 mW (633 nm) laser lines; a 40x C-Apochromat water immersion objective (NA 1.2, Korr M27, WD 0.22 mm) or 60x C-Plan-Apochromat oil immersion objective (NA 1.4, DIC M27, WD 0.14 mm); conventional photomultiplier tube (PMT) detectors; and ZEN Black acquisition software. *Leica Stellaris*: Imaging was performed using 1–3% laser power from a freely tunable white-light laser (485–685 nm); a 40x HC PL APO CS2 oil immersion objective (NA 1.3, WD 0.24 mm); highly sensitive HyD S detectors; and LAS X acquisition software. For both systems, the frame size and zoom were adjusted to ensure a minimum of 2x Nyquist sampling in the xy plane (pixel size ≤ Δ*_xy_/*2), where Δ*_xy_* denotes the lateral resolution, estimated as Δ*_xy_* ≈ 0.61*λ/NA* based on the acquisition settings. Full three-dimensional image stacks of entire MCC were acquired at 30 s or 60 s intervals with an axial resolution (optical section thickness) of Δ*_z_* ≈ 2*nλ/NA*^2^. Image stacks were sampled with a z-step size of ∼0.4 µm, corresponding to an optical section of ∼0.9 µm, thereby satisfying the Nyquist criterion for axial sampling (z-step size ≤ Δ*_z_/*2).

##### 2.1.2 High-speed imaging

To capture fine BB movements at the MCC apical domain, imaging was performed using a custom-built total internal reflection fluorescence (TIRF) microscope based on an Olympus IX83 inverted platform. Imaging was carried out using 0.5–2% laser power from Cobolt 200 mW laser lines (488 nm and 561 nm); a 150x universal apochromat Olympus TIRF oil-immersion objective (NA 1.45, WD 0.08 mm); a high-speed Hamamatsu ImagEM X2 electron-multiplying charge-coupled device (EMCCD) camera (106.667 nm per pixel for 150x 1.45 NA objective); and cellSens acquisition software. Using this setup, 30,000 images of BBs were acquired over 10.5 min at the rate of 1 frame every 21 ms. The frame size and zoom were adjusted to ensure 2x Nyquist sampling in the lateral (xy) plane for single-plane TIRF imaging, with all images acquired at a fixed focal plane throughout the time lapse acquisition. The excitation focus position was maintained by a motorized Olympus TIRF module, and axial drift was continuously corrected using an Olympus IX3-ZDC2 z-drift compensation system. Photobleaching was minimized using a real-time controller that synchronized camera exposure with an acousto-optic tunable filter (AOTF).

##### 2.1.3 High-resolution imaging

High-resolution imaging of the actin meshwork was performed using the Airyscan superresolution mode on a Zeiss LSM980 microscope. Imaging was carried out using 1–3% laser power from diode 30 mW (488 nm), DPSS 25 mW (561 nm), DPSS 8 mW (594 nm), and diode 25 mW (639 nm) laser lines; a 63x Plan-Apochromat oil-immersion objective (NA 1.4, WD 0.19 mm); an Airyscan2 detector with a 32-channel gallium arsenide phosphide (GaAsP) detector array; and ZEN Blue 3 acquisition software. The frame size and zoom were adjusted to ensure a minimum of 2× Nyquist sampling in the lateral (xy) plane. Z-stacks were acquired to cover the entire apical cortex of MCCs with a z-step size of ∼0.3 µm using a piezo stage, corresponding to an optical section of ∼0.9 µm, thereby satisfying the Nyquist criterion for axial sampling.

#### 2.2 Pre-acquisition considerations for coarse-timed imaging

During imaging of BB distribution and apical domain expansion, ongoing morphogenetic movements of the embryo caused continuous lateral (xy) drift of the region of interest (ROI). To facilitate downstream image analysis, the field of view (FOV) was continuously monitored throughout acquisition, and the ROI was manually repositioned by stage movement whenever drift was observed. Similarly, the z-stack range was adjusted as needed to compensate for axial drift, ensuring that the full height of the MCC remained within the defined imaging volume at all time points. Laser power and detector gain were adjusted during acquisition to maintain consistent visibility of BBs and apical actin throughout the imaging period.

#### 2.3 Pre-acquisition considerations for fine-timed imaging

Fine-timed imaging was performed using a TIRF setup, which requires a planar sample for proper visualization of BBs moving across the apical plane. Therefore, only cells with a flat apical profile were selected to avoid losing BBs moving out of the TIRF plane. Using this setup, we found that BB fluorescence (Chibby-GFP) remained stable and trackable for 10.5 min at 1 frame every 21 ms. Apical areas of different sizes were imaged for 10.5 min to characterize BB trajectories across the full apical area range. To measure the apical area, we first acquired 400 frames in the actin channel, followed by 30,000 frames in the BB channel. To assess microscope stability, including lateral (xy) and axial (z) drift, and to provide reference objects for localization noise correction during particle detection and tracking, we imaged 0.5 µm diameter beads (∼ BB size) attached to coverslips. These beads were imaged with the same settings (1 frame every 21 ms for 10.5 min), yielding 30,000 frames of bead data. Bead imaging was performed prior to BB experiments to validate the setup and provide reference data for downstream image analysis.

### 3 Softwares

All image and data analyses were performed using established plugins and custom-written scripts in ImageJ^117^, Fiji^118^, and MATLAB R2023b (MathWorks, Natick, MA, USA). Analyses were carried out on a Windows 11-based workstation equipped with 192 GB of RAM, an 8 GB graphics card, and an Intel Xeon W-125 CPU (2.5 GHz). Statistical analyses were conducted using GraphPad Prism 11. Figures and schematics were generated using Adobe Illustrator 27 and Inkscape 1.4. Detailed descriptions of all image analysis methods are provided in the Supplementary Information.

## Acknowledgments

We thank the John Wallingford group from the University of Texas at Austin for the alpha-Tubulin-LifeAct-GFP and alpha-Tubulin-LifeAct-RFP constructs and the Brian Mitchell group from the Northwestern University Feinberg School of Medicine for sharing the alpha-Tubulin-Chibby-GFP and alpha-Tubulin-Chibby-RFP constructs. We are grateful to the *Xenopus* facility caretakers at the University of Copenhagen; Jutta Bulkescher and the reNEW Imaging Platform, and the Core Facility for Integrated Bioimaging (Faculty of Health and Medical Sciences, University of Copenhagen) for technical support and assistance with imaging. We thank Lene Oddershede and Akbar Samadi, formerly a postdoctoral researcher in her group at the Niels Bohr Institute, for technical assistance and discussions on fine-timed experiments. We thank Guilherme Bastos Ventura for critical reading of the manuscript and for valuable comments. We also thank members of the Sedzinski group for their valuable comments and suggestions.

## Funding

This work was supported by grants from Novo Nordisk Fonden (NNF21CC0073729, J.S.), European Research Council Consolidator Grant (ERC CoG 101125803 MechanoFate, J.S.). J.S. acknowledges the support of the Novo Nordisk Foundation (NNF22OC0076414, NNF19OC0056962) and the Leo Foundation (LF-OC-19-000219). P.M.B and Y.F.B acknowledge Novo Nordisk Foundation grant number NNF20OC0061176.

## Author contributions

R.T. conceived and designed the study, performed all experiments, analyzed the data, and wrote the manuscript with contributions from all other authors. Y.F.B. performed TIRF imaging experiments and TIRF data analysis. P.B. supervised TIRF experiments. M.M.I. conceived and designed the study, generated the theoretical model description, and helped with data analysis. J.S. conceived and designed the study, supervised the project, and acquired funding.

## Use of Artificial Intelligence tools

The authors acknowledge the use of ChatGPT, Gemini, Perplexity, and Claude for brain-storming, editing, coding and technical assistance, and proofreading during the preparation of this manuscript. All content generated with the assistance of these tools was subsequently reviewed and edited by the authors, who take full responsibility for the final version of the manuscript.

## Data availability

The data and materials supporting the findings of this study are available within the article and its supplementary information files. Additional information and relevant raw data are available from the corresponding authors upon request.

## Code availability

The code used for data analysis and simulations is available in the GitHub repositories BB analysis and BB simulations, respectively.

## Competing interests

The authors declare no competing interests.

## Notes

### Competing Interest Statement

The authors have declared no competing interest.

