## Supplementary Information for "Actin cross-linking organizes basal body patterning through anomalous diffusion transitions"

### Contents

**Supplementary figures and Movies** 4

**1 Supplementary figures** 4

**2 Movie captions** 23

**Theoretical Model** 26

**3 Dynamics of Basal Body movements at fine-grained time-scales** 26

**4 Coarse-grained basal body dynamics** 28

**5 Results** 32

**6 Simulation parameters** 38

**Methods** 40

**7 Image analysis for coarse-timed experiments** 40

**11 Image analysis for figures and movies** **56**

**12 Statistical analysis** **56**

**List of materials** **58**

**References** **60**

Supplementary figures and Movies

### 1 Supplementary figures

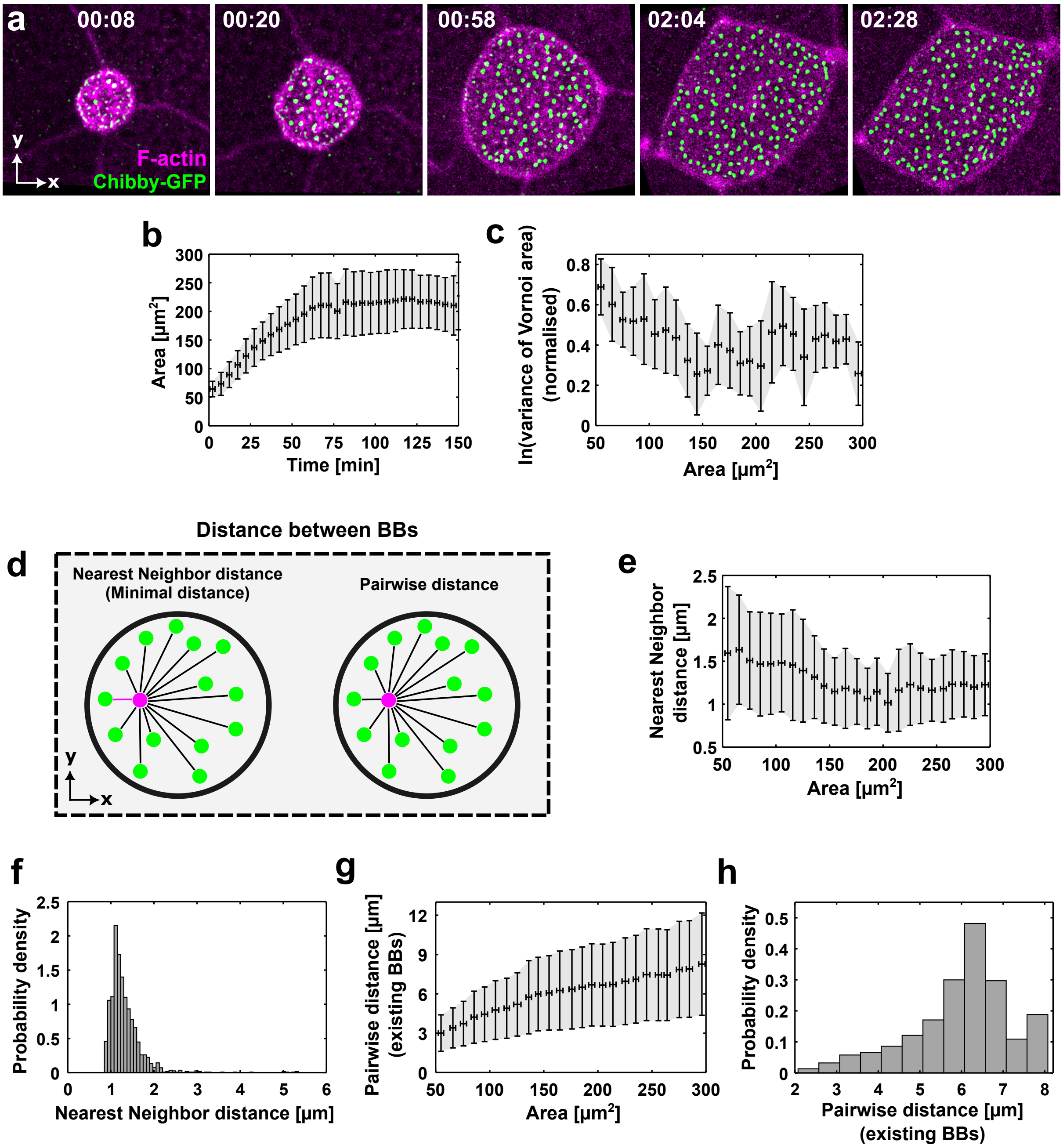

**Figure S1. Basal Body (BB) patterning is synchronized with apical surface expansion.** All data correspond to coarse-timed imaging (frame interval = 30–60 s; total duration = 2.5 h). (**a**) Time-lapse sequence of BBs (Chibby-GFP, green) embedded within the actin cortex (LifeAct-RFP, pink) of the expanding apical domain, corresponding to Fig. 1a (see Movie 1). Time is in hh:mm. (**b**) Apical area plotted against time. (**c**) Variance of Voronoi tessellation areas plotted against apical area. Variance values are natural log–transformed (ln) and min–max normalized. (**d**) Schematic explaining the calculation of minimal (nearest-neighbor) and pairwise distances between BBs. The reference BB (pink) is compared with cohabiting BBs (green); pink line marks the nearest neighbor and the black lines mark the pairwise distances *(see methods section 7.3.4 in SI).* (**e**) Nearest-neighbor distance plotted against apical area. (**f**) Probability density of nearest-neighbor distances with a peak at $1.23\,\mu m^2$. (**g**) Pairwise distance plotted against apical area. (**h**) Probability density of pairwise distances with a peak at $6.5\,\mu m$. Distances in (**f**) and (**h**) were obtained from 9 cells across 5 experiments, covering 1487 frames and $\sim 97,000$ BBs (f) and $\sim 1,048,000$ BBs (h). Distances were computed for all BBs in each frame, then frame-averaged and pooled over the full 2.5 h time-lapse sequence. In (**b**), (**c**), (**e**), and (**g**), error bars represent mean $\pm$ standard deviation (SD) of data averaged over 9 cells from 5 experiments, using 5 min (time) or $10\,\mu m^2$ (area) bins.

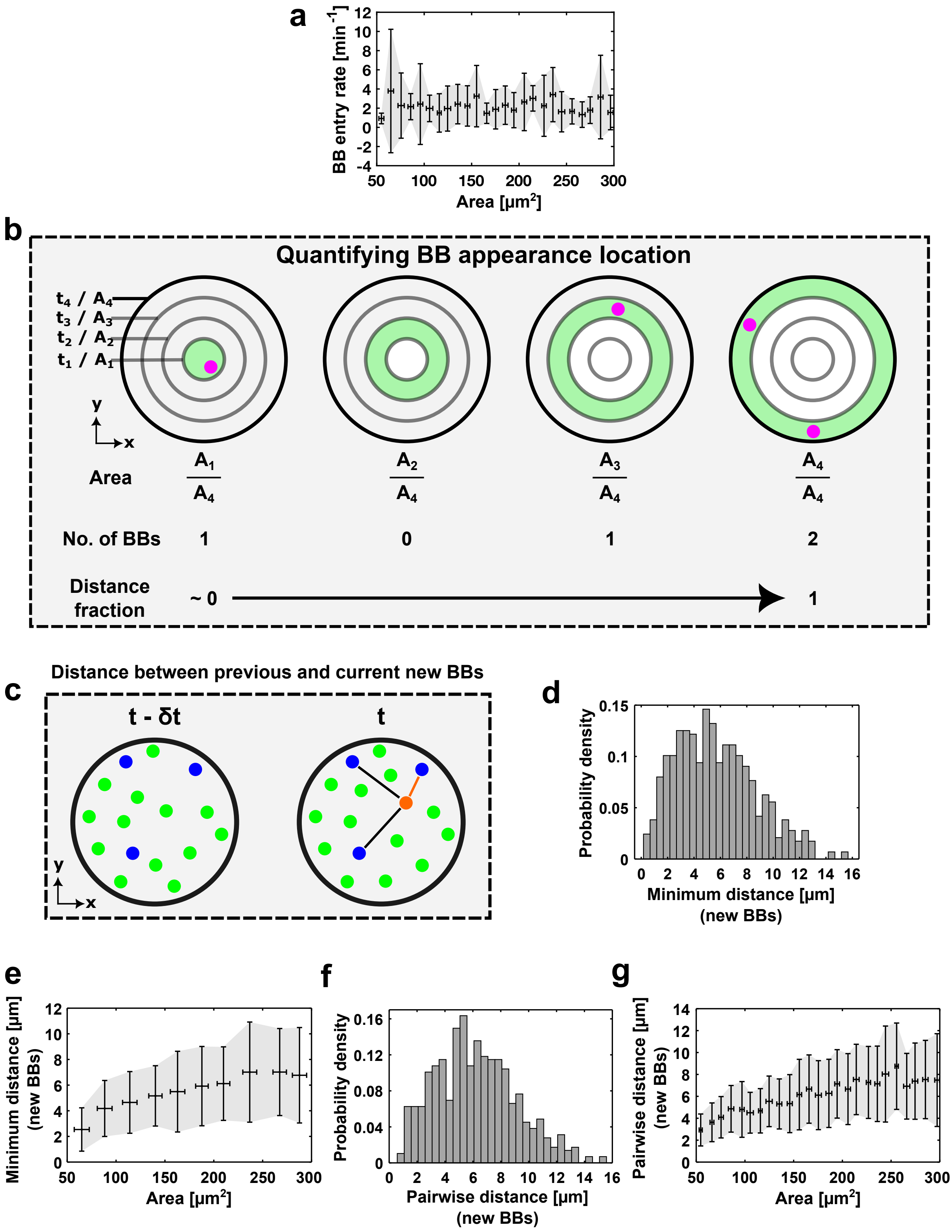
a
BB entry rate [min^-1]
Area [μm^2]
b
Quantifying BB appearance location
t4 / A4
t3 / A3
t2 / A2
t1 / A1
y
x
Area
A1/A4
A2/A4
A3/A4
A4/A4
No. of BBs
1
0
1
2
Distance fraction
~ 0
1
c
Distance between previous and current new BBs
t - δt
t
d
Probability density
Minimum distance [μm] (new BBs)
e
Minimum distance [μm] (new BBs)
Area [μm^2]
f
Probability density
Pairwise distance [μm] (new BBs)
g
Pairwise distance [μm] (new BBs)
Area [μm^2]

**Figure S2. Passive and active mechanisms shape BB distribution.** All data correspond to coarse-timed imaging (frame interval = 30–60 s; total duration = 2.5 h). (**a**) Entry rate of BBs plotted against apical area. (**b**) Schematic showing calculation of BB appearance location, corresponding to Fig. 2c. Black contours represent apical boundaries; the solid black contour marks the boundary at the current time point, whereas semi-transparent contours mark boundaries at previous time points. The green donut-shaped regions between successive contours indicate the newly formed apical area. Pink dots mark BBs. The area, BB count and distance fraction for the current time point are given below the schematic (*see methods section 7.3.6 in SI for description*). (**c-g**) Distances between newly appearing BBs at consecutive time points. (**c**) Schematic showing minimal and pairwise distances between new BB at time $t$ (orange) and at the previous time point $t - \delta t$ (blue). Lines (black and orange) indicate pairwise distances; orange highlights the minimal distance; green shows pre-existing BBs (*see methods section 7.3.4 in SI for description*). (**d**) Probability density of minimal distances with a peak at 5.3 µm. (**e**) Minimal distance plotted against apical area. (**f**) Probability density of pairwise distances with a peak at 6 µm. (**g**) Pairwise distance plotted against apical area. In (**a**) and (**d-g**), data are pooled from 9 cells across 5 experiments. In (**d-g**), distances were calculated for all BBs in each frame and then frame-averaged before pooling across the full 2.5 h time-lapse sequences of all cells. The plots in (**d**) and (**f**) were computed from 851 minimal and 1221 pairwise distances for 575 frames. In (**a**), (**e**) and (**g**), error bars represent mean ± SD of binned data (10 µm$^2$ bins).

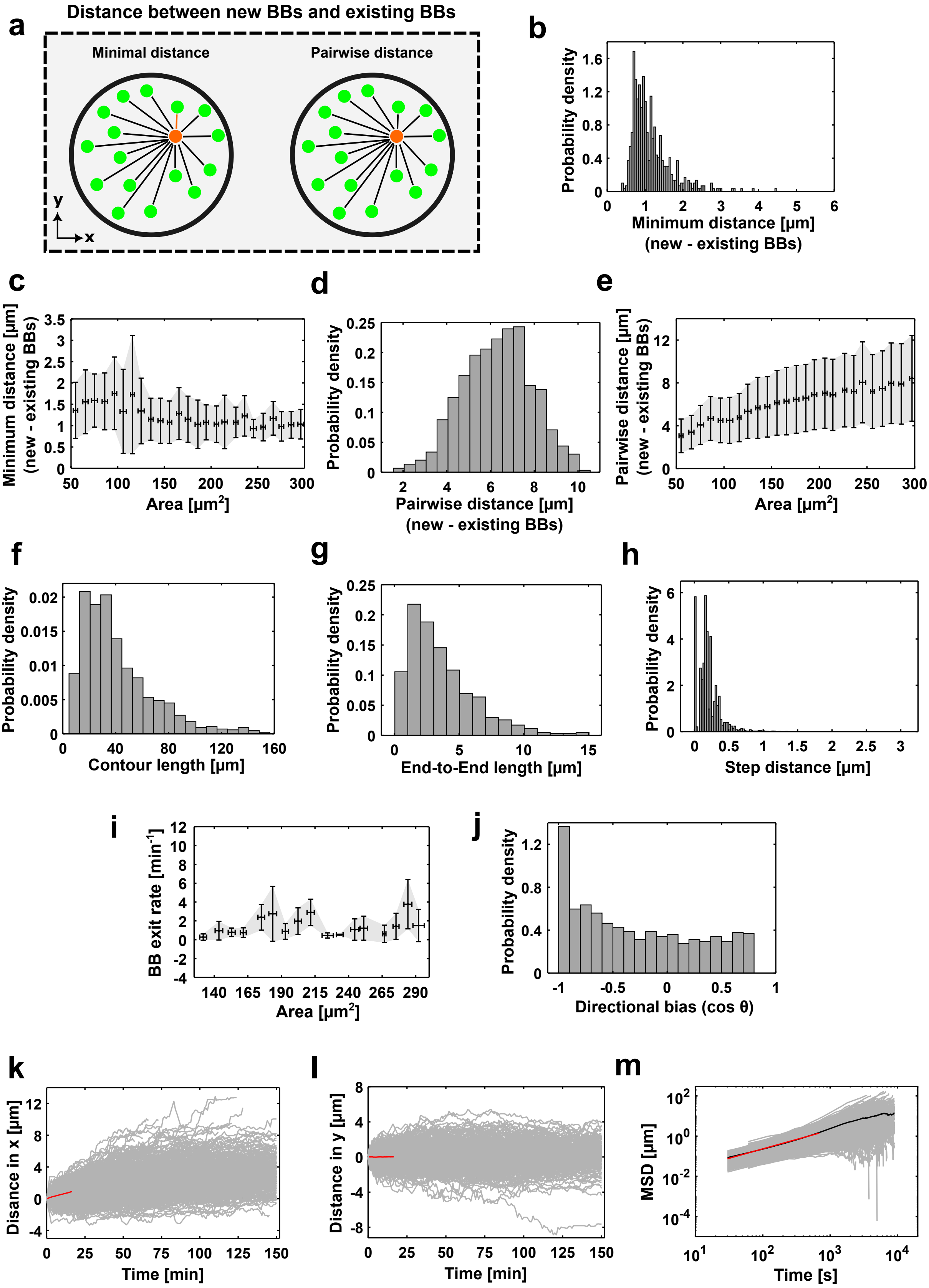
a
Distance between new BBs and existing BBs
Minimal distance
Pairwise distance
y
x
b
Probability density
Minimum distance [μm]
(new - existing BBs)
c
Minimum distance [μm]
(new - existing BBs)
Area [μm2]
d
Probability density
Pairwise distance [μm]
(new - existing BBs)
e
Pairwise distance [μm]
(new - existing BBs)
Area [μm2]
f
Probability density
Contour length [μm]
g
Probability density
End-to-End length [μm]
h
Probability density
Step distance [μm]
i
BB exit rate [min-1]
Area [μm2]
j
Probability density
Directional bias (cos θ)
k
Disance in x [μm]
Time [min]
l
Distance in y [μm]
Time [min]
m
MSD [μm]
Time [s]

**Figure S3. Passive and active mechanisms shape BB distribution.** All data correspond to coarse-timed imaging (frame interval = 30–60 s; total duration = 2.5 h). (**a-e**) Distances between new and pre-existing BBs. (**a**) Schematic showing distances between new BBs (orange) and pre-existing BBs (green). Black lines indicate pairwise distances, and orange highlights the minimal distance (*see methods section 7.3.4 in SI for description*). (**b**) Probability density of minimal distances with a peak at 1 µm. (**c**) Minimal distance plotted against apical area. (**d**) Probability density of pairwise distances with a peak at 6.6 µm. (**e**) Pairwise distance plotted against apical area. In (**b-e**), data are pooled from 9 cells across 5 experiments. The distances were calculated for all BBs in each frame and then frame-averaged before pooling across the full 2.5 h time-lapse sequences of all cells. The plots in (**b**) and (**d**) were computed from 898 minimal and 39326 pairwise distances for 593 frames. In (**c**) and (**e**), error bars represent mean ± SD of binned data (10 µm$^2$ bins). (**f**) Probability density of contour lengths of BB trajectories. (**g**) Probability density of end-to-end displacements of BB trajectories. (**h**) Probability density of BB step sizes. (**i**) Exit rate of BBs from the apical domain as a function of apical area. The error bars represent mean ± SD of data averaged over 9 cells from 5 experiments, using 10 µm$^2$ bins. (**j**) Probability density of the normalized dot product values quantifying trajectory direction relative to the periphery of the apical domain. (**k**) Mean x-displacement of all trajectories (grey: coherent drift component) and their ensemble average (red), plotted against time. (**l**) Mean y-displacements (grey: stochastic component; red: ensemble average), plotted against time. (**m**) Mean squared displacement (MSD) of trajectories against lag time. Grey curves: individual MSDs; black: ensemble average; red: fit line. A total of 1052 trajectories from 9 cells across 5 experiments were analyzed for (**f-h**) and (**j-m**).

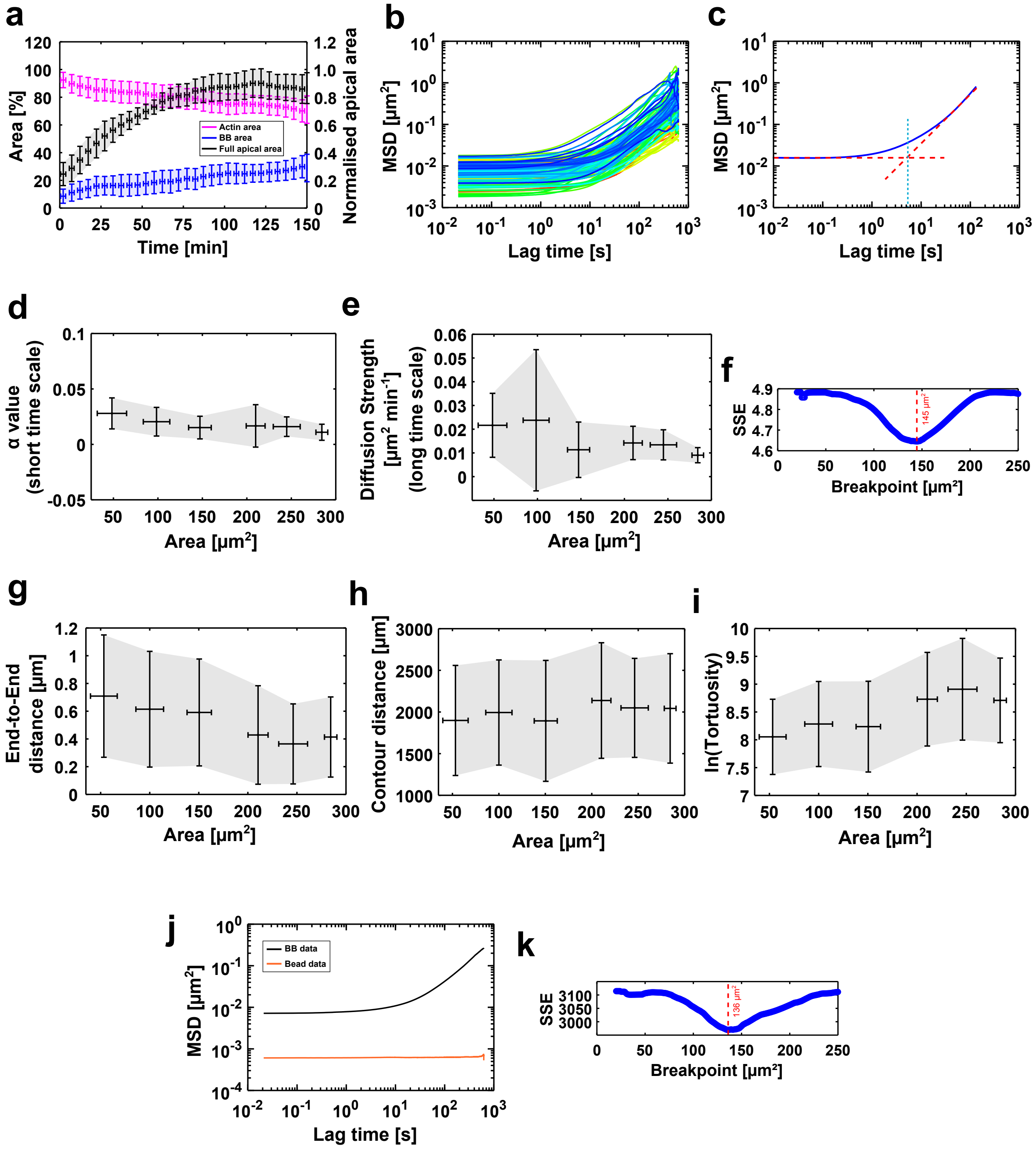
a
Area [%]
Normalised apical area
Time [min]
Actin area
BB area
Full apical area
b
MSD [μm²]
Lag time [s]
c
MSD [μm²]
Lag time [s]
d
α value
(short time scale)
Area [μm²]
e
Diffusion Strength
[μm² min⁻¹]
(long time scale)
Area [μm²]
f
SSE
145 μm²
Breakpoint [μm²]
g
End-to-End
distance [μm]
Area [μm²]
h
Contour distance [μm]
Area [μm²]
i
ln(Tortuosity)
Area [μm²]
j
MSD [μm²]
BB data
Bead data
Lag time [s]
k
SSE
136 μm²
Breakpoint [μm²]

**Figure S4. BB dynamics shift with apical domain expansion.** (**a**) Apical area percentages plotted against time. Black: total apical area (see Fig. S1b). Blue: area fraction occupied by BBs. Pink: area fraction occupied by actin. Data from coarse-timed imaging (frame interval = 30–60 s; total duration = 2.5 h). Error bars represent mean ± SD of data averaged over 9 cells across 5 experiments, using 5 min bins. (**b-k**) Data from fine-timed imaging (frame interval = 21 ms; 30,000 frames; total duration = 10.5 min). (**b**) Ensemble-averaged MSDs of BB trajectories from 100 cells across 17 experiments, plotted against lag time. (**c**) Schematic illustrating transition time extraction from MSD: blue - MSD curve; red dotted lines - short- and long-time fits; blue dotted line - intersection (transition time) (*see methods section 8.3.3 in SI*). (**d**) Short-time $\alpha$ values plotted against apical area. (**e**) Long-time diffusion strength plotted against apical area. (**f**) Sum of Squared Errors (SSE) of long-time $\alpha$ values fits (from Fig. 3c), plotted across apical area; the red dotted line marks the breakpoint area (lowest SSE). (**g**) End-to-end displacement plotted against area. (**h**) Contour length plotted against area. (**i**) Tortuosity (natural log–transformed) plotted against area. (**j**) Ensemble-averaged MSDs of BBs (black; 4,604 trajectories from 100 cells) and 0.5 µm beads (orange; 47 beads) (*see methods section 8.3.6 in SI*). (**k**) SSE of transition time fits (from Fig. 3e), plotted across apical area; the red dotted line marks the breakpoint area (lowest SSE). In (**d, e, g-i**), error bars represent mean ± SD of data averaged over 4,604 trajectories from 100 cells across 17 experiments, using 50 $\mu m^2$ bins.

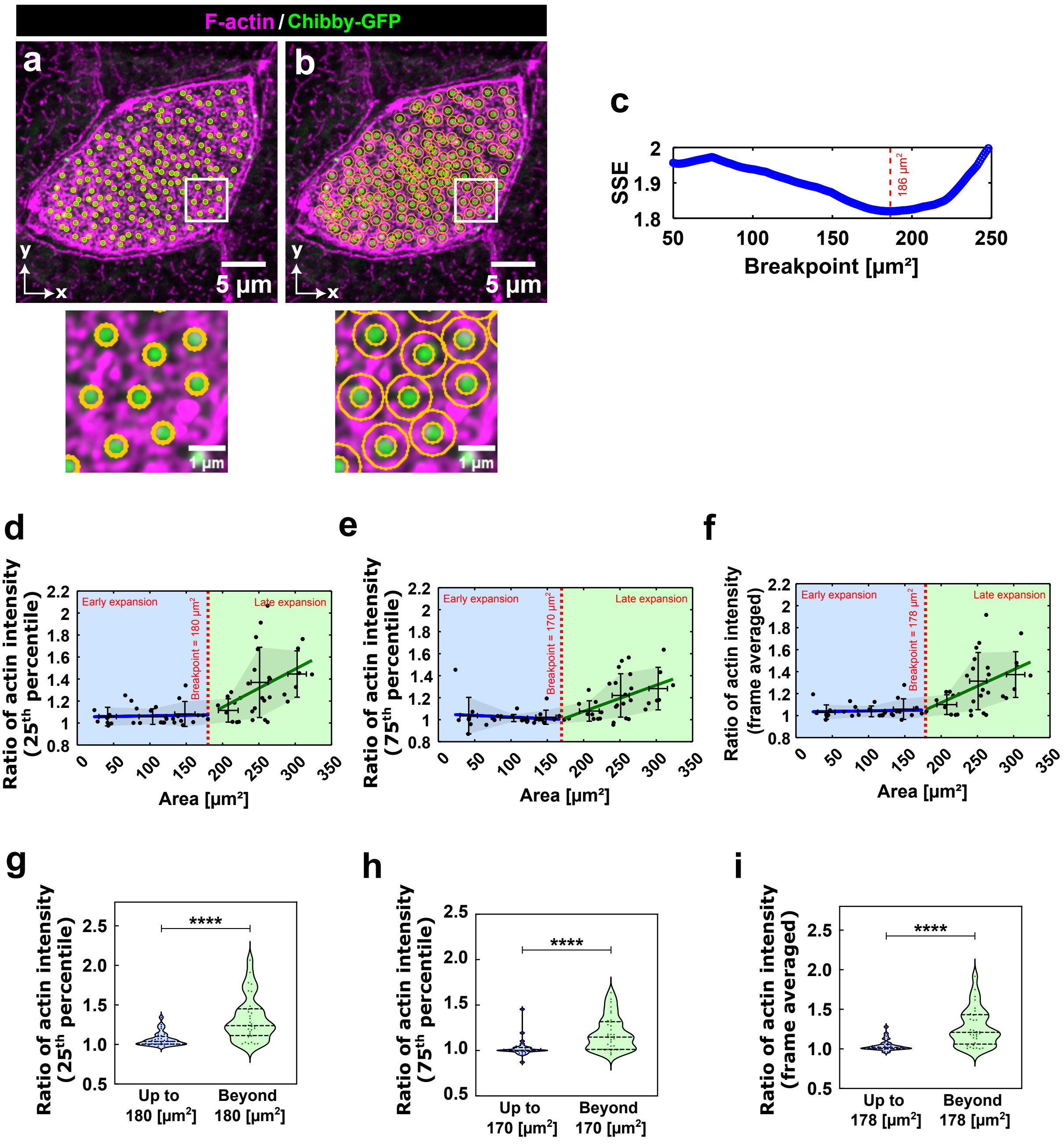

F-actin / Chibby-GFP
a
b
5 μm
5 μm
1 μm
1 μm
c
SSE
186 μm²
Breakpoint [μm²]
d
e
f
Early expansion
Late expansion
Breakpoint = 180 μm²
Breakpoint = 170 μm²
Breakpoint = 178 μm²
Ratio of actin intensity (25th percentile)
Ratio of actin intensity (75th percentile)
Ratio of actin intensity (frame averaged)
Area [μm²]
g
h
i
****
Up to 180 [μm²]
Beyond 180 [μm²]
Up to 170 [μm²]
Beyond 170 [μm²]
Up to 178 [μm²]
Beyond 178 [μm²]

**Figure S5. Progressive formation of an apical actin meshwork facilitates BB redistribution.** All data from high-resolution airyscan imaging. (**a-b**) Quantification of actin intensity at BB positions (**a**) and surrounding regions (**b**); zoomed-in ROIs from white areas are shown below. Orange circular ROI: position of the BB used for quantifying actin intensity at the BB; orange donut ROI: annular region used to quantify actin intensity in the surrounding regions. (**c**) SSE of actin intensity ratios at $50^{th}$ percentile of CDF (from Fig. 4d), across apical area. Red dotted line highlights the breakpoint area (lowest SSE). (**d-e**) Ratios of actin intensities (surrounding / BB positions) plotted against apical area at $25^{th}$ (**d**) and $75^{th}$ (**e**) percentiles of CDF. (**f**) Ratio of frame-averaged surrounding actin intensity to frame-averaged actin intensity at BB positions. (**d, e, f**) Each dot represents a single cell and error bars represent mean ± SD of data averaged over 64 cells from 4 experiments, using $50\,\mu m^2$ bins. Red dotted lines mark the breakpoints identified by piecewise linear regression: $25^{th}$ percentile of CDF - $180\,\mu m^2$; $75^{th}$ percentile of CDF - $170\,\mu m^2$; frame-averaged - $178\,\mu m^2$, while blue and green lines are regression fits before and after the breakpoints. The blue and green shaded regions denote data before and after the breakpoints and serve as visual guides for the early ($<150\,\mu m^2$) and late ($>150\,\mu m^2$) phases of apical expansion. (**g-i**) Plots comparing ratios before vs. after breakpoints, using data from 63 cells across 4 experiments. (**g**) $25^{th}$ percentile of CDF, corresponding to (d): $P < 0.0001$; 32 cells ($\leq 180\,\mu m^2$), 32 cells ($>180\,\mu m^2$). (**h**) $75^{th}$ percentile of CDF, corresponding to (e): $P < 0.0001$; 30 cells ($\leq 170\,\mu m^2$), 34 cells ($>170\,\mu m^2$). (**i**) frame-averaged, corresponding to (f): $P < 0.0001$; 31 cells ($\leq 178\,\mu m^2$), 33 cells ($>178\,\mu m^2$). In (**g-i**), the central line represents the median, and the upper and lower lines represent the $75^{th}$ and $25^{th}$ percentiles, respectively.

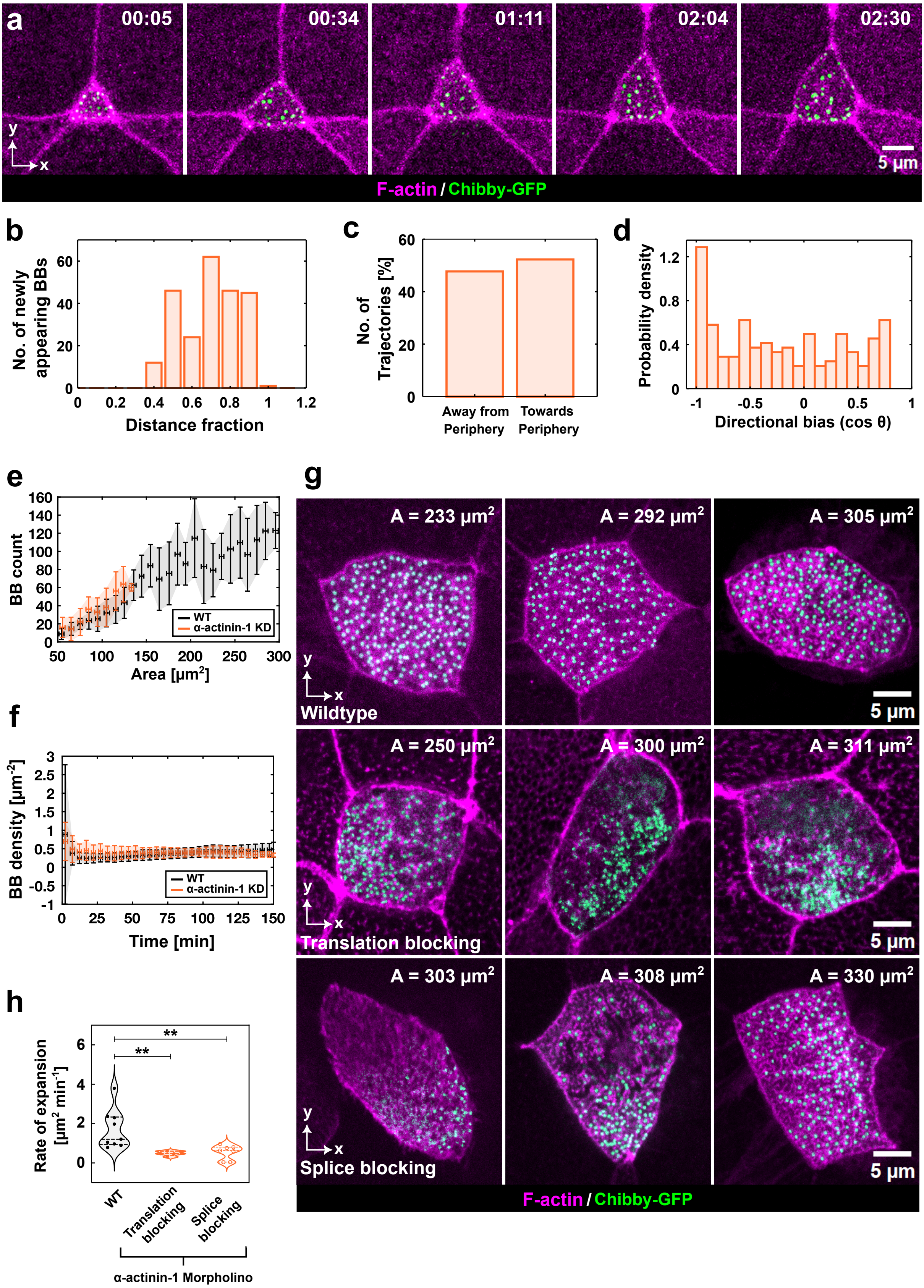

a
00:05
00:34
01:11
02:04
02:30
5 µm
F-actin / Chibby-GFP
b
No. of newly appearing BBs
Distance fraction
c
No. of Trajectories [%]
Away from Periphery
Towards Periphery
d
Probability density
Directional bias (cos θ)
e
BB count
Area [µm²]
WT
α-actinin-1 KD
f
BB density [µm⁻²]
Time [min]
WT
α-actinin-1 KD
g
A = 233 µm²
A = 292 µm²
A = 305 µm²
Wildtype
A = 250 µm²
A = 300 µm²
A = 311 µm²
Translation blocking
A = 303 µm²
A = 308 µm²
A = 330 µm²
Splice blocking
5 µm
F-actin / Chibby-GFP
h
Rate of expansion [µm² min⁻¹]
**
**
WT
Translation blocking
Splice blocking
α-actinin-1 Morpholino

**Figure S6. Disruption of apical actin crosslinking impairs BB organization.** Data in (**a-f**) and (**h**) correspond to coarse-timed imaging (frame interval = 30–60 s; total duration = 2.5 h). (**a-d**) Analyses of $\alpha$-actinin-1 translation-blocking morpholino injected cells. (**a**) Time-lapse sequence of BBs (Chibby-GFP, green) embedded in apical actin (LifeAct-RFP, pink), corresponding to Fig. 1a. Time is in hh:mm (see Movie 9). (**b**) Distribution of newly appearing BBs across distance fractions, where 0 corresponds to the center and 1 to the periphery of the apical domain. (**c**) Percentage of trajectories moving towards or away from the expanding periphery. (**d**) Probability density of normalized dot product values quantifying trajectory direction relative to the apical periphery. (**e-f**) Comparison of WT (black) and $\alpha$-actinin-1 translation-blocking morpholino (orange)–injected cells. Error bars represent mean ± SD of binned data: $10\,\mu m^2$ (area bins) or 5 min (time bins).(**e**) Number of BBs plotted against time. (**f**) BB density over the apical area plotted against time. (**g**) BB distributions in large apical domains for WT (top row), $\alpha$-actinin-1 translation-blocking morpholino (middle row), and $\alpha$-actinin-1 splice-blocking morpholino (bottom row) injected cells. (**h**) Apical expansion rates compared between WT, $\alpha$-actinin-1 translation-blocking morpholino (exact P = 0.0018), and $\alpha$-actinin-1 splice-blocking morpholino (exact P = 0.0097) conditions. The central line represents the median, and the upper and lower lines represent the $75^{th}$ and $25^{th}$ percentiles, respectively. In (**b-d**), data are from 5 cells across 4 experiments. In (**e-f**) and (**h**), data are from 9 cells across 5 experiments for WT, 5 cells across 4 experiments for $\alpha$-actinin-1 translation-blocking morpholino, and 7 cells across 3 experiments for $\alpha$-actinin-1 splice-blocking morpholino conditions.

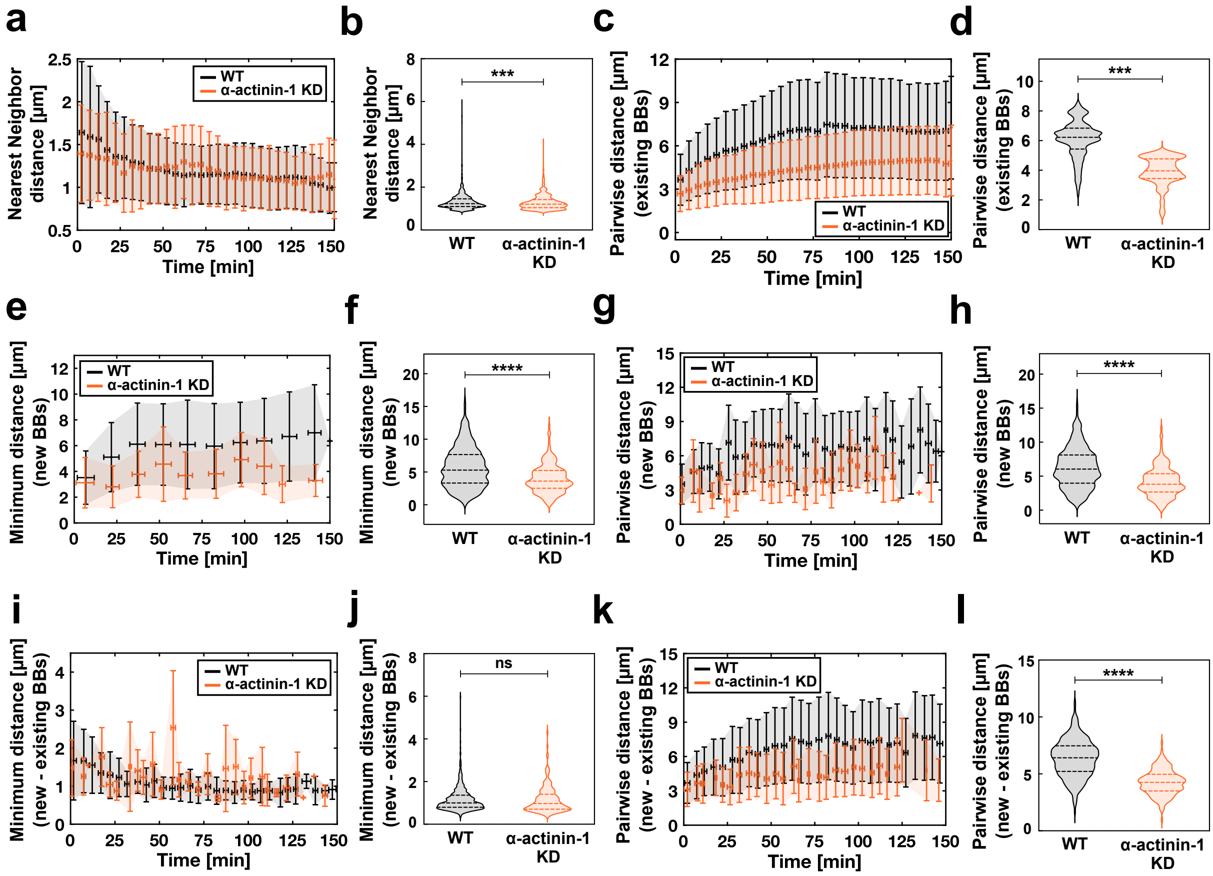
a
b
c
d
e
f
g
h
i
j
k
l
Nearest Neighbor distance [μm]
Pairwise distance [μm] (existing BBs)
Minimum distance [μm] (new BBs)
Pairwise distance [μm] (new BBs)
Minimum distance [μm] (new - existing BBs)
Pairwise distance [μm] (new - existing BBs)
Time [min]
WT
α-actinin-1 KD
***
****
ns

**Figure S7. Disruption of apical actin crosslinking impairs BB organization.** All data correspond to coarse-timed imaging (frame interval = 30–60 s; total duration = 2.5 h). WT (black) and $\alpha$-actinin-1 translation-blocking morpholino (orange)–injected cells are compared and plotted for an experimental time of 2.5 h. (**a-d**) Distances between BBs in the current frame: (**a**) nearest neighbor distance evolution; (**b**) frame-averaged minimal distances; (**c**) pairwise distance evolution; (**d**) frame-averaged pairwise distances. (**e-h**) Distances between newly appearing BBs at consecutive frames: (**e**) minimal distance evolution; (**f**) frame-averaged minimal distances; (**g**) pairwise distance evolution; (**h**) frame-averaged pairwise distances. (**i-l**) Distances between new and pre-existing BBs in the current frame: (**i**) minimal distance evolution; (**j**) frame-averaged minimal distances; (**k**) pairwise distance evolution; (**l**) frame-averaged pairwise distances. In (**a**), (**c**), (**e**), (**g**), (**i**) and (**k**), error bars represent mean ± SD of binned data (5 min bins). Statistics for violin plots: (**b**) $\sim 97,000$ distances from 1,487 frames (WT) vs. $\sim 12,000$ distances from 1,131 frames (KD), $P \approx 0.0004$; (**d**) $\sim 1,050,000$ distances from 1,487 frames (WT) vs. $\sim 670,000$ from 1,131 (KD), $P < 0.0001$; (**f**) 851 distances from 575 frames (WT) vs. 204 from 177 (KD), $P < 0.0001$; (**h**) 1,221 distances from 575 frames (WT) vs. 239 from 177 (KD), $P < 0.0001$; (**j**) 898 distances from 593 frames (WT) vs. 146 from 177 (KD), $P = 0.1964$; (**l**) 39,326 distances from 593 frames (WT) vs. 4,831 from 177 (KD), $P < 0.0001$. In all the violin plots, the central line represents the median, and the upper and lower lines represent the $75^{th}$ and $25^{th}$ percentiles, respectively. Data were obtained from 9 cells across 5 experiments (WT) and 5 cells across 4 experiments (KD).

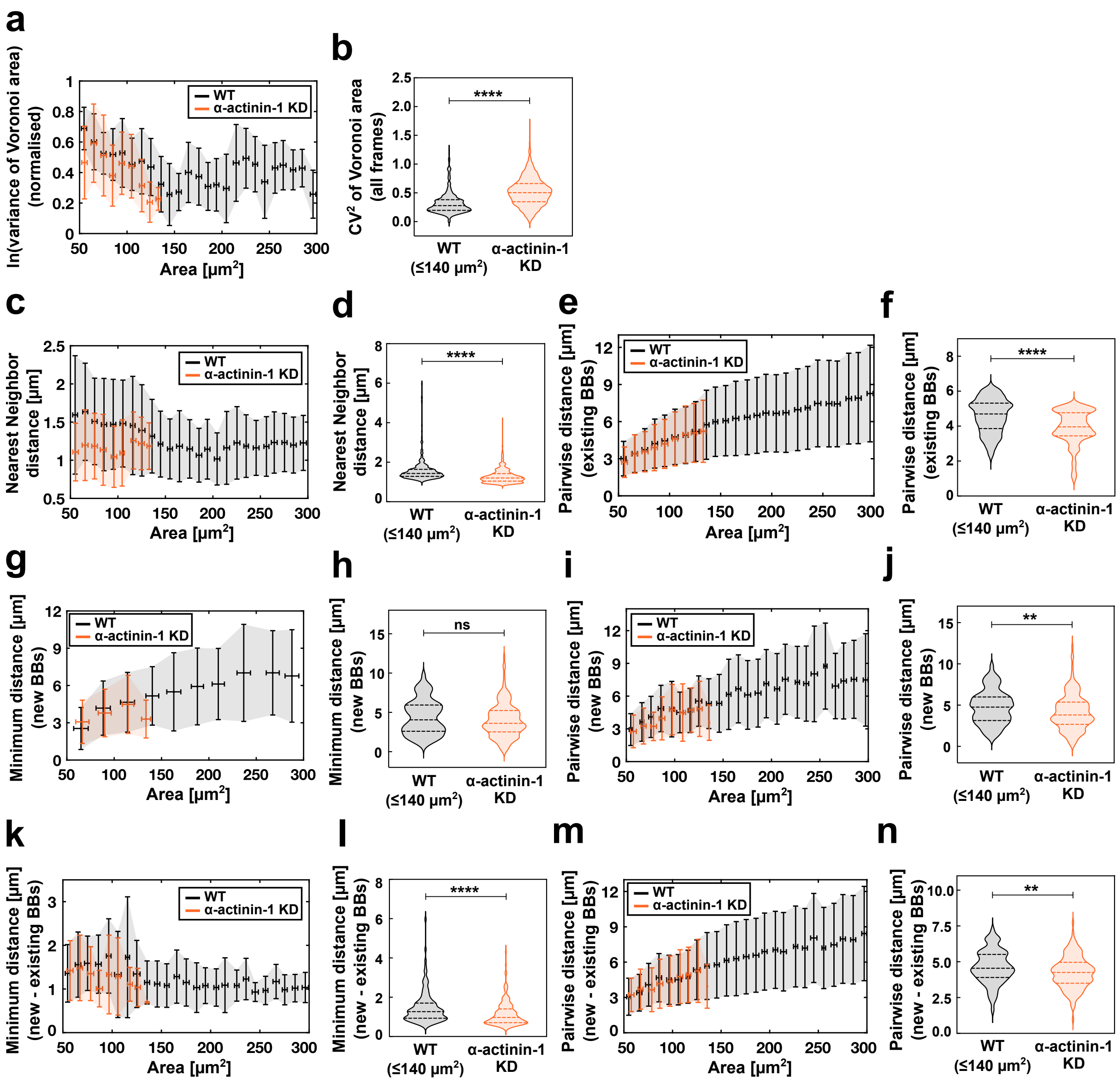
a
ln(variance of Voronoi area) (normalised)
Area [μm²]
WT
α-actinin-1 KD
b
CV² of Voronoi area (all frames)
****
WT (≤140 μm²)
α-actinin-1 KD
c
Nearest Neighbor distance [μm]
Area [μm²]
WT
α-actinin-1 KD
d
Nearest Neighbor distance [μm]
****
WT (≤140 μm²)
α-actinin-1 KD
e
Pairwise distance [μm] (existing BBs)
Area [μm²]
WT
α-actinin-1 KD
f
Pairwise distance [μm] (existing BBs)
****
WT (≤140 μm²)
α-actinin-1 KD
g
Minimum distance [μm] (new BBs)
Area [μm²]
WT
α-actinin-1 KD
h
Minimum distance [μm] (new BBs)
ns
WT (≤140 μm²)
α-actinin-1 KD
i
Pairwise distance [μm] (new BBs)
Area [μm²]
WT
α-actinin-1 KD
j
Pairwise distance [μm] (new BBs)
**
WT (≤140 μm²)
α-actinin-1 KD
k
Minimum distance [μm] (new - existing BBs)
Area [μm²]
WT
α-actinin-1 KD
l
Minimum distance [μm] (new - existing BBs)
****
WT (≤140 μm²)
α-actinin-1 KD
m
Pairwise distance [μm] (new - existing BBs)
Area [μm²]
WT
α-actinin-1 KD
n
Pairwise distance [μm] (new - existing BBs)
**
WT (≤140 μm²)
α-actinin-1 KD

**Figure S8. Disruption of apical actin crosslinking impairs BB organization.** All data correspond to coarse-timed imaging (frame interval = 30–60 s; total duration = 2.5 h). WT (black) and $\alpha$-actinin-1 translation-blocking morpholino (orange)–injected cells are compared as a function of morphogenetic time (apical area). Data for WT in the violin plots are plotted up to the maximum apical area reached in KD (140 µm$^2$). (**a**) Evolution of variance. Variance values are natural log–transformed (ln) and min–max normalized. (**b**) Comparison of variability in Voronoi tessellation areas pooled across all frames (2.5 h). Variability is quantified as $CV^2$, the squared coefficient of variation (variance divided by the square of the mean). (**c-f**) Distances between BBs in the current frame: (**c**) minimal distance evolution, (**d**) frame-averaged minimal distances, (**e**) pairwise distance evolution, (**f**) frame-averaged pairwise distances. (**g-j**) Distances between newly appearing BBs across consecutive frames: (**g**) minimal distance evolution, (**h**) frame-averaged minimal distances, (**i**) pairwise distance evolution, (**j**) frame-averaged pairwise distances. (**k-n**) Distances between new and pre-existing BBs in the current frame: (**k**) minimal distance evolution, (**l**) frame-averaged minimal distances, (m) pairwise distance evolution, (**n**) frame-averaged pairwise distances. In (**a**), (**c**), (**e**), (**g**), (**i**), (**k**), and (**m**), error bars represent mean $\pm$ SD of binned data (10 µm$^2$ bins). Statistics for violin plots: (**b**) variances from 428 frames (WT) and 1176 frames (KD). (**d**) $\sim 7,000$ distances from 414 frames (WT) vs. $\sim$ 12,000 distances from 1129 frames (KD), P $<$ 0.0001; (**f**) $\sim 148,000$ distances from 414 frames (WT) vs. $\sim 670,000$ from 1129 (KD), P $<$ 0.0001; (**h**) 178 distances from 137 frames (WT) vs. 204 from 177 (KD), P $\approx$ 0.0897; (**j**) 294 distances from 137 frames (WT) vs. 239 from 177 (KD), P $\approx$ 0.0049; (**l**) 184 distances from 137 frames (WT) vs. 146 from 177 (KD), P $<$ 0.0001; (**n**) 2992 distances from 137 frames (WT) vs. 4831 from 177 (KD), P = 0.0025. In all the violin plots, the central line represents the median, and the upper and lower lines represent the 75$^{\text{th}}$ and 25$^{\text{th}}$ percentiles, respectively. Data were obtained from 9 cells across 5 experiments (WT) and 5 cells across 4 experiments (KD).

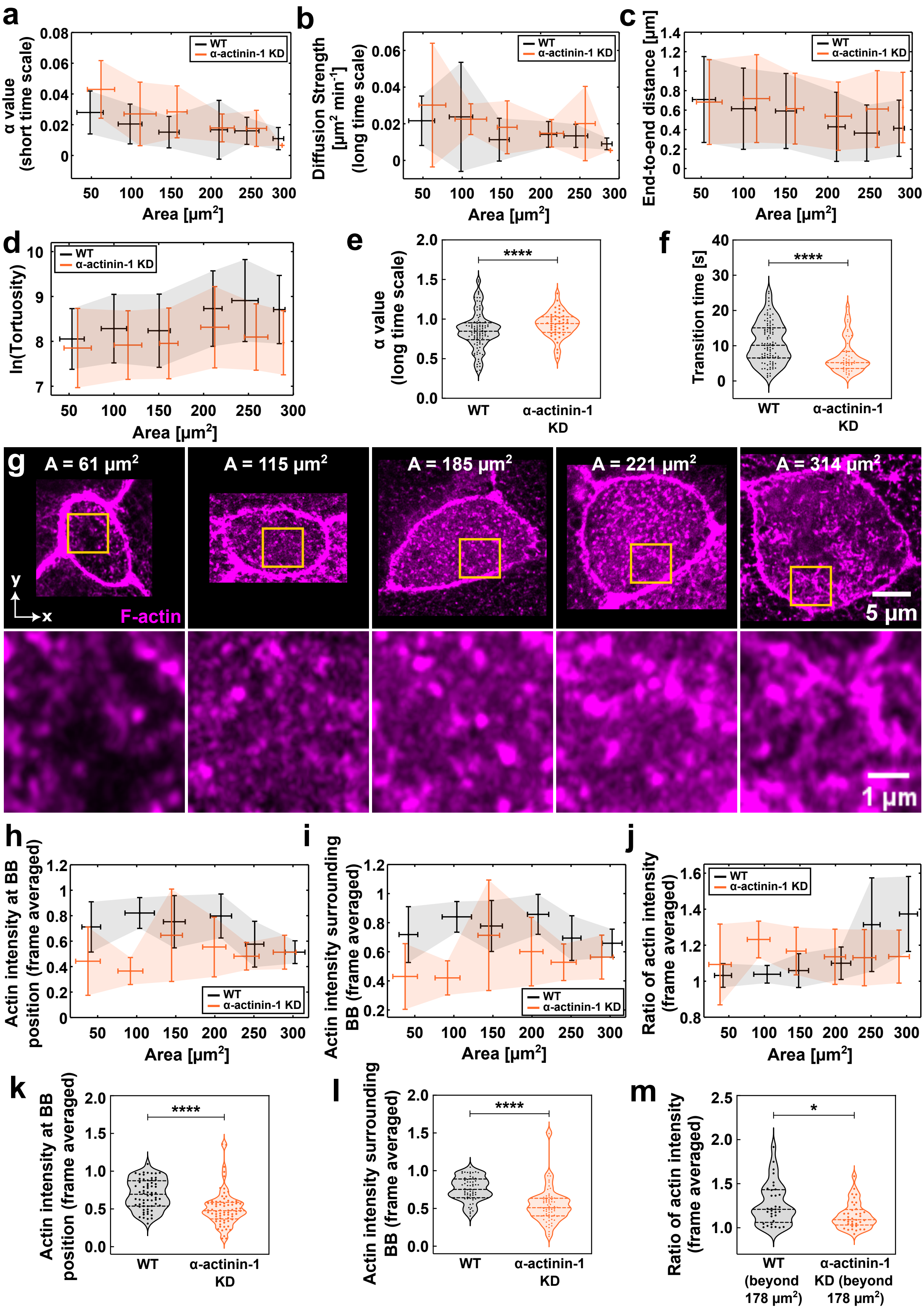
a
α value (short time scale)
WT
α-actinin-1 KD
Area [μm$^2$]
b
Diffusion Strength [μm$^2$ min$^{-1}$] (long time scale)
WT
α-actinin-1 KD
Area [μm$^2$]
c
End-to-end distance [μm]
WT
α-actinin-1 KD
Area [μm$^2$]
d
ln(Tortuosity)
WT
α-actinin-1 KD
Area [μm$^2$]
e
α value (long time scale)
****
WT
α-actinin-1 KD
f
Transition time [s]
****
WT
α-actinin-1 KD
g
A = 61 μm$^2$
A = 115 μm$^2$
A = 185 μm$^2$
A = 221 μm$^2$
A = 314 μm$^2$
y
x
F-actin
5 μm
1 μm
h
Actin intensity at BB position (frame averaged)
WT
α-actinin-1 KD
Area [μm$^2$]
i
Actin intensity surrounding BB (frame averaged)
WT
α-actinin-1 KD
Area [μm$^2$]
j
Ratio of actin intensity (frame averaged)
WT
α-actinin-1 KD
Area [μm$^2$]
k
Actin intensity at BB position (frame averaged)
****
WT
α-actinin-1 KD
l
Actin intensity surrounding BB (frame averaged)
****
WT
α-actinin-1 KD
m
Ratio of actin intensity (frame averaged)
*
WT (beyond 178 μm$^2$)
α-actinin-1 KD (beyond 178 μm$^2$)

**Figure S9. $\alpha$-actinin-1 perturbation impairs transition in BB dynamics and actin meshwork formation.** Data in (**a-f**) and (**h-m**) compare WT (black) and $\alpha$-actinin-1 translation-blocking morpholino (orange) conditions. Data in (**a-f**) correspond to fine-timed imaging (frame interval = 21 ms; 30,000 frames; total duration = 10.5 min). (**a**) Short-time $\alpha$ values versus apical area. (**b**) Long-time diffusion strength versus apical area. (**c**) End-to-end displacement versus apical area. (**d**) Tortuosity (natural log–transformed) versus apical area. (**e**) Comparison of long-time $\alpha$ values, corresponding to Fig. 6c; exact $P = 0.0071$. (**f**) Comparison of transition times, corresponding to Fig. 6d; $P < 0.0001$. Error bars in (**a-d**) represent mean $\pm$ SD of binned data ($50\,\mu m^2$ bins). Data in (**a-f**) include 4,604 trajectories from 100 cells across 17 experiments (WT) and 2,029 trajectories from 46 cells across 5 experiments (KD). Data in (**g-m**) correspond to high-resolution Airyscan imaging. (**g**) Images of apical domains of increasing size showing the lack of actin meshwork formation, corresponding to Fig. 6e, 6f. Top row: actin (Phalloidin). Bottom row: zoom of regions highlighted in orange squares in the top row. (**h-m**) compare the frame-averaged actin intensity at BB position (**h, k**), frame-averaged surrounding actin intensity (**i, l**), and ratio of frame-averaged actin intensity between surrounding and BB position (**j, m**). Data are from 64 cells across 4 experiments (WT) and 72 cells across 3 experiments (KD). (**h-j**) Compare the evolution across apical area and error bars represent mean $\pm$ SD of binned data ($50\,\mu m^2$ bins). (**k-m**) Violin plots for statistical comparison of: (**k**) frame-averaged actin intensities at BB position ($P < 0.0001$); (**l**) in the surrounding ($P < 0.0001$); and (**m**) their ratios beyond the breakpoint ($178\,\mu m^2$, see Fig. S5f), (exact $P = 0.0123$); data from 33 cells across 4 experiments (WT) and 37 cells from 3 experiments (KD). In all the violin plots, the central line represents the median, and the upper and lower lines represent the $75^{th}$ and $25^{th}$ percentiles, respectively.

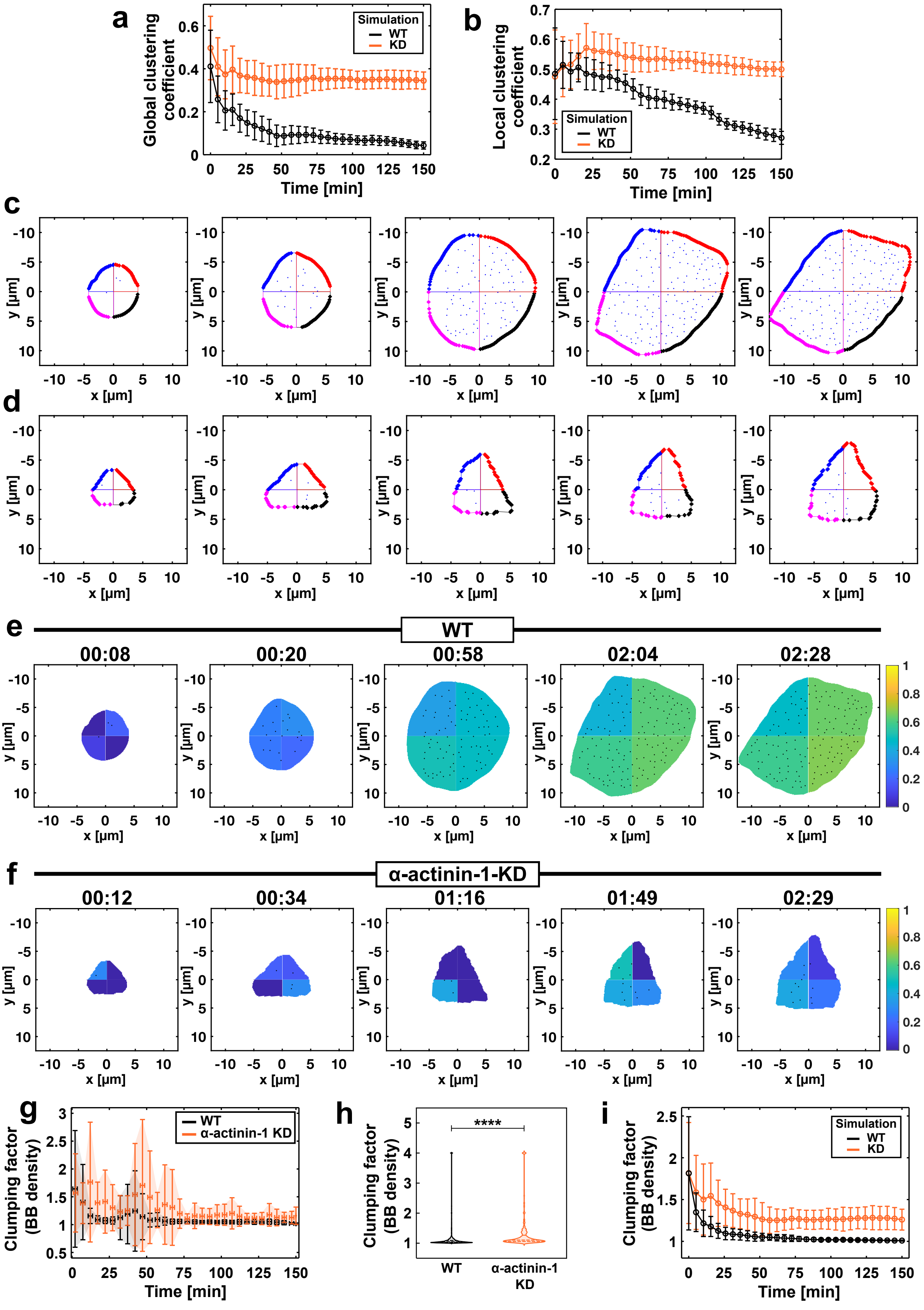
a
Global clustering coefficient
Simulation
WT
KD
Time [min]
b
Local clustering coefficient
Simulation
WT
KD
Time [min]
c
d
y [μm]
x [μm]
e
WT
00:08
00:20
00:58
02:04
02:28
f
α-actinin-1-KD
00:12
00:34
01:16
01:49
02:29
g
Clumping factor (BB density)
WT
α-actinin-1 KD
Time [min]
h
Clumping factor (BB density)
****
WT
α-actinin-1 KD
i
Clumping factor (BB density)
Simulation
WT
KD
Time [min]

**Figure S10. Heterogeneous distribution of the apical actin network leads to BB clumping.** (**a,b,i**) Data from the model, comparing WT (black) and KD (orange) simulations. (**c-h**) Data from coarse-timed imaging (frame interval = 30–60 s; total duration = 2.5 h), comparing WT and $\alpha$-actinin-1 translation-blocking morpholino conditions. (**a-b**) Time evolution for global (**a**) and local (**b**) clustering simulations. (**c-d**) Time-lapse sequences of apical domains showing quadrant discretization and BBs within each quadrant for WT (**c**) and KD conditions (**d**), corresponding to (Fig. S10e, 7f) (WT) and (Fig. S10f, 7g) (KD) (see Movie 15). (**e-f**) Time-lapse sequences of apical domains discretized into four quadrants and color-coded for BB density: WT (**e**) and KD (**f**) (see Movie 16). (**g**) Time evolution of the clumping factor for BBs. Error bars represent mean ± SD of binned data (5 min bins). (**h**) Comparisons of clumping factors between WT and KD for BBs; data obtained from 1,704 frames (WT) and 1,182 frames (KD), $P < 0.0001$. The central line represents the median, and the upper and lower lines represent the 75$^{\text{th}}$ and 25$^{\text{th}}$ percentiles, respectively. Data in (**g**) and (**h**) were obtained from 9 cells across 5 experiments (WT) and 5 cells across 4 experiments (KD).(**i**) Time evolution of the clumping factor for BBs from the simulations. In (**a**), (**b**) and (**i**), the error bars represent mean ± SD of data averaged over 20 simulations, using 5 min bins.

### 2 Movie captions

**Movie 1.** Left: BB (Chibby-GFP, green); Right: actin (LA-RFP, pink) and BB, showing the BB docking and distribution during the apical domain expansion in a WT MCC. Time in hh:mm.

**Movie 2.** 3D visualization of BB (Chibby-GFP, green) ascent from basal to apical side of a WT MCC, coupled with docking to the apical domain during expansion. Time in hh:mm.

**Movie 3.** Voronoi tessellations of BBs on the apical surface of a WT MCC. Left: data showing BBs (Chibby-GFP, green) used for constructing tessellations; the apical periphery is shown as a white contour. Right: extracted tessellations in pink with BBs as green dots. Time in hh:mm.

**Movie 4.** BB tracking and trajectory evolution in a WT MCC. The apical periphery is shown as a white contour. Left: data showing BBs (Chibby-GFP, green) used for tracking. Right: white dots represent BBs, and trajectories are shown in diverse colors. Time in hh:mm.

**Movie 5.** Spatial position of newly appearing BBs in a WT MCC. The apical periphery is shown as a white contour. Left: data showing BBs (Chibby-GFP, green). Right: circles mark the positions where new BBs appear. Time in hh:mm.

**Movie 6.** Exit and entry events of a BB (Chibby-GFP, green) in a WT apical domain shown from three different views: xy (top), xz (bottom), and yz (right). Yellow arrows in the xz and yz views continuously track the BB both while docked at the apical surface and after exiting beneath the apical domain. The yellow arrow in the xy view appears only when the BB is docked at the apical surface. The animated plot below shows the corresponding Z-position variation of the BB; '0' on the y-axis marks the apical surface. Time in mm:ss.

**Movie 7.** Fine-timed imaging sequences (frame interval = 21 ms; 30,000 frames; total duration = 10.5 min) of BBs (Chibby-GFP, green) overlaid with trajectories for four different apical domain sizes (WT). For visualization and file-size compatibility, maximum intensity projection was applied to every 30 frames, and the movie was downsized to 1000 frames, resulting in an apparent frame interval of 630 ms over a total duration of 10.5 min. Time in mm:ss.

**Movie 8.** WT model simulation of basal body distribution during apical domain expansion. BBs are shown as green dots within the expanding circular apical domain boundary (magenta circle). Simulation units are converted to experimental time and spatial scales to enable direct comparison with experiments. Time is shown in min.

**Movie 9.** Left: BB (Chibby-GFP, green); Right: actin (LA-RFP, pink) and BB, showing the BB docking and distribution during the apical domain expansion of an MCC injected with $\alpha$-actinin-1 translation-blocking morpholino. Time in hh:mm.

**Movie 10.** 3D visualization of BB (Chibby-GFP, green) ascent from basal to apical side, coupled with docking to the apical domain during expansion in an MCC injected with $\alpha$-actinin-1 translation-blocking morpholino. Time in hh:mm.

**Movie 11.** BB tracking and trajectory evolution in an MCC injected with $\alpha$-actinin-1 translation-blocking morpholino. The apical periphery is shown as a white contour. Left: data showing BBs (Chibby-GFP, green) used for tracking. Right: white dots represent BBs, and trajectories are shown in diverse colors. Time in hh:mm.

**Movie 12.** Voronoi tessellations of BBs on the apical surface of an MCC injected with $\alpha$-actinin-1 translation-blocking morpholino. Left: data showing BBs (Chibby-GFP, green) used for constructing tessellations; the apical periphery is shown as a white contour. Right: extracted tessellations in pink with BBs as green dots. Time in hh:mm.

**Movie 13.** Fine-timed imaging sequences (frame interval = 21 ms; 30,000 frames; total duration = 10.5 min) of BBs (Chibby-GFP, green) overlaid with trajectories for four different apical domain sizes ($\alpha$-actinin-1 translation-blocking morpholino condition). For visualization and file-size compatibility, maximum intensity projection was applied to every 30 frames, and the movie was downsized to 1000 frames, resulting in an apparent frame interval of 630 ms over a total duration of 10.5 min. Time in mm:ss.

**Movie 14.** KD model simulation of basal body distribution during apical domain expansion. BBs are shown as green dots within the expanding circular apical domain boundary (magenta circle). Simulation units are converted to experimental time and spatial scales to enable direct comparison with experiments. Time is shown in min.

**Movie 15.** Quadrant discretization of apical domains during expansion, showing BB docking and distribution for WT and $\alpha$-actinin-1 translation-blocking morpholino conditions. Each quadrant is marked by a distinct color, and BBs are shown as blue dots.

Time in hh:mm.

**Movie 16.** Discretized quadrants of apical domains color-coded for BB density and mean actin intensity. Top and bottom rows correspond to WT and KD conditions, while left and right columns correspond to BB density and mean actin intensity, respectively. BBs are shown as black dots. Time in hh:mm.

### Theoretical Model

#### 3 Dynamics of Basal Body movements at fine-grained time-scales

The basal bodies (BB) exhibit stochastic movements as per experimental tracking. The tracking was experimentally done in two modes. In the fine-grained mode, the time-interval between two frames was $\Delta t_{\mathrm{fg}} = 20$ ms and the total duration of measurement was 20 min. In the coarse-grained mode, the time-interval was $\Delta t_{\mathrm{cg}} = 30$ s and the measurement for done for the entire duration $\approx 150$ min of a typical experiment.

In the fine-mode, the following features of BB movements were observed. As a function of time-lag $\Delta t$, the mean square displacement of a particular BB could be approximately represented as:

$$\mathrm{MSD}(\Delta t) \approx \begin{cases} \Delta R_0^2, & \Delta t < \tau, \\ 4D\Delta t^{\alpha}, & \Delta t > \tau. \end{cases} \tag{1}$$

This indicated that the basal bodies are caged within a region of size $\Delta R_0$ for time-lags $\Delta t$ smaller than $\tau$, and they undergo anomalous diffusion with exponent $\alpha$ on longer time-scales. The typical time-scales for $\tau$ were experimentally found to be of the order $5 - 15$ s and the exponent $\alpha$ approximately in the range $0.4 - 1.4$, thus exhibiting both subdiffusion and superdiffusion.

To model this process, we adapted the framework developed by Metzner et al.[1]. We assumed that a BB $i$ is caged inside a harmonic trap with a diffusivity $D_w$ of the BB within the cage with a trap relaxation time of $\tau_w$. The trap itself is modeled to undergo anomalous diffusion with strength $D$ and exponent $\alpha$ (Figure M1). The equations of motion for the position $\mathbf{r}_{bi}$ and $\mathbf{r}_{ti}$ of the BB and the center of the harmonic trap are, respectively, given as

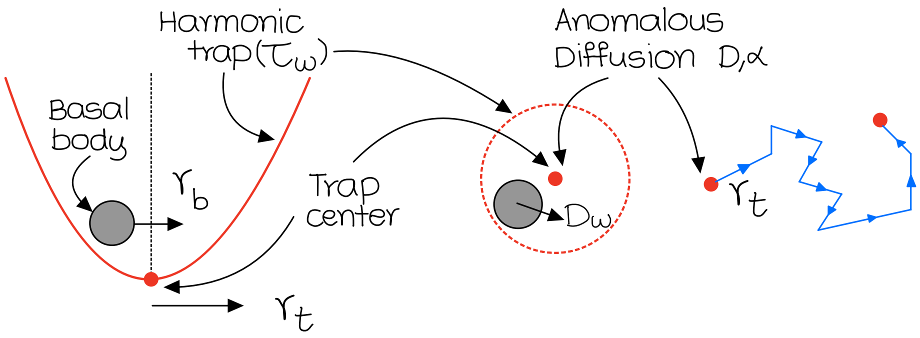

**Figure M1.** In the fine-grained model, the early time caged movements of the basal body (BB), with position $\mathbf{r}_b$, within the actin pocket is modeled as a harmonic trap of relaxation time $\tau_w$. The BB diffuses within the trap with diffusivity $D_w$ . The trap center, with position $\mathbf{r}_t$, itself undergoes anomalous diffusion with exponent $\alpha$ and strength $D$.

$$\frac{d}{dt}\mathbf{r}_{bi} = -\frac{1}{\tau_w}(\mathbf{r}_{bi} - \mathbf{r}_{ti}) + \boldsymbol{\xi}_i(t), \tag{2}$$

$$\frac{d}{dt}\mathbf{r}_{ti} = \boldsymbol{\eta}_i(t), \tag{3}$$

where $\boldsymbol{\xi}_i(t)$ is standard Gaussian noise with zero mean $\langle \boldsymbol{\xi}_i \rangle = \mathbf{0}$ and delta correlation $\langle \boldsymbol{\xi}_i(t) \cdot \boldsymbol{\xi}_i(t') \rangle = 4D_w\delta(t - t')$ and $\boldsymbol{\eta}_i$ is non-Gaussian noise with zero mean $\langle \boldsymbol{\eta}_i \rangle = \mathbf{0}$ and power-law correlation $\langle \boldsymbol{\eta}_i(t) \cdot \boldsymbol{\eta}_i(t') \rangle = 4D|t - t'|^{\alpha}$. We use standard Euler-Maruyama algorithm to integrate these equations as

$$\mathbf{r}_{bi}^{t+dt} = \mathbf{r}_{bi}^{t} - \frac{1}{\tau_w}(\mathbf{r}_{bi}^{t} - \mathbf{r}_{ti}^{t}) + \sqrt{2D_w dt}\Delta G(t), \tag{4}$$

$$\mathbf{r}_{ti}^{t+dt} = \mathbf{r}_{ti}^{t} + \sqrt{2D}dt^{H}\Delta N(t), \tag{5}$$

where $H = \alpha/2$ is the so-called Hurst exponent and $dt$ is the time-step of the simulation[2]. $\Delta G$ is a sequence of random numbers from Gaussian distribution with zero mean and unit standard deviation. $\Delta N(t)$ are the increments obtained from the Davis-Harte process, corresponding to fractional Brownian motion (fBM) that is used to model the anomalous diffusion of the actin trap. Increments for fBM were generated using the circulant embedding method[3,4], as implemented via the `fbm` Python package[5] and adapted for MATLAB.

To capture the experimentally observed BB-to-BB heterogeneity in anomalous dynamics, the exponent $\alpha$ is sampled independently for each realisation from a Gaussian distribution with mean $\mu = 0.85$ and standard deviation $\sigma = 0.2$, clipped to the range $[0.4, 1.4]$ consistent with the experimentally observed spread. Since $D$ carries units $\mu\mathrm{m}^2\,\mathrm{s}^{-\alpha}$ that depend on $\alpha$, using a fixed numerical value across realisations with different $\alpha$ would be physically inconsistent. We therefore normalise $D$ per realisation as $D = D_{\mathrm{raw}} \cdot t_{\mathrm{ref}}^{1-\alpha}$, where $D_{\mathrm{raw}} = 0.0161$ $\mu\mathrm{m}^2/\mathrm{min}$ is the mean diffusion strength fitted from fine-timed experimental trajectories and $t_{\mathrm{ref}} = 10$ s is the mean elbow transition time measured experimentally. This ensures $\mathrm{MSD}(t_{\mathrm{ref}}) = 4D_{\mathrm{raw}}\,t_{\mathrm{ref}}$ is identical across all realisations regardless of $\alpha$. A sample mean square displacement (MSD) obtained using this model is shown in Figure M2.

The model admits simple analytical insights. In the short-time limit $\Delta t \ll \tau_w$, the BB displacement within the trap saturates, giving a plateau $\Delta R_0^2 = 4D_w\tau_w$. Equating this with the long-time anomalous diffusion MSD $4D\tau^{\alpha}$ gives the elbow time

$$\tau = \left(\frac{D_w\tau_w}{D}\right)^{1/\alpha}, \tag{6}$$

at which the BB transitions from caged to anomalous diffusion. The numerical values of these quantities are discussed in the caption of Figure M2.

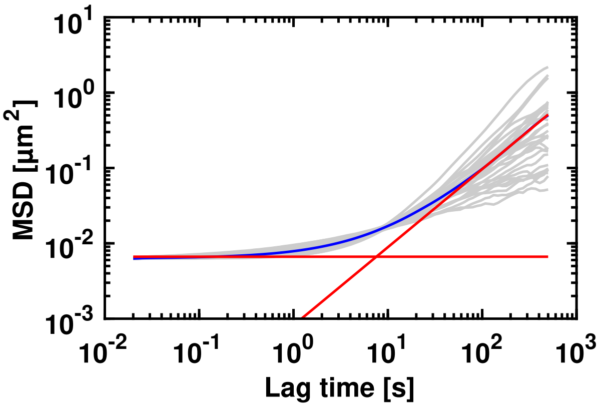

**Figure M2.** MSDs of simulated BBs: grey, individual particles; blue, ensemble average; red lines, fitted regimes; their intersection defines the transition time. Data obtained from 30 independent runs of the single-particle simulation. The moving trap model described in the text and shown in Figure M1 exhibits commensurate MSD with an initial caging regime followed by anomalous diffusion. The fitted ensemble-averaged MSD yields a plateau $\Delta R_0^2 \sim 10^{-2}$ $\mu$m$^2$, consistent with $4D_w\tau_w$, and a power-law regime with $D_{\text{eff}} \approx 0.014$ $\mu\text{m}^2\,\text{min}^{-\alpha}$ and $\alpha \approx 1.04$, close to $D_{\text{raw}} \cdot t_{\text{ref}}^{1-\alpha} \approx 0.017$ $\mu\text{m}^2\,\text{min}^{-\alpha}$. The intersection of the two fitted red lines gives the caging or elbow time $\tau \approx 7.6$ s $\sim 10$ s, consistent with the analytical estimate $\tau = (D_w\tau_w/D_{\text{eff}})^{1/\alpha} \approx 6.9$ s and the experimentally observed range 5–15 s. See Table 1 for the full list of model parameters.

The mechanism described in Figure M1 is applicable to our system. This is supported by our experimental observations showing that basal bodies are surrounded by an actin network that can effectively trap them. Moreover, the actin network undergoes remodeling as the apical surface evolves and can be effectively thought to undergo stochastic movements that statistically show signature of anomalous diffusion.

### 4 Coarse-grained basal body dynamics

In the coarse-grained experimental mode of BB tracking, the time-interval between two frames, $\Delta t_{\text{cg}} = 30$ s $> \tau \approx 5 - 15$ s, the trapping time-scale observed experimentally. Hence, at this experimental resolution, we do not observe the initial trapping. However, in this case the MSD fits reasonably well with

$$\text{MSD}(\Delta t) = 4D\Delta t^{\alpha}, \tag{7}$$

where all the terms have identical interpretation as earlier. A majority of the phenomena reported in our experiments was quantified using coarse-grained imaging in which the initial trapping behavior is not captured. Hence, while modeling the complete BB dynamics, in conjunction with the apical surface growth, we do not incorporate the initial trapping dynamics, but instead focus on the anomalous diffusion BB dynamics. The complete model is detailed below.

Our computational model simulates the spatiotemporal dynamics of basal body organization in ciliated epithelial cells. The model incorporates several key biological features that govern basal body positioning and clustering behavior during apical emergence. We model that the apical surface of the cell grows dynamically while maintaining its circular geometry throughout the simulation. The surface area increases according to a prescribed time-dependent function that reflects the natural expansion patterns observed experimentally. Figure M3 demonstrates this growth pattern, showing how the apical surface area evolves over time for wild-type (WT) and knockdown (KD) simulations.

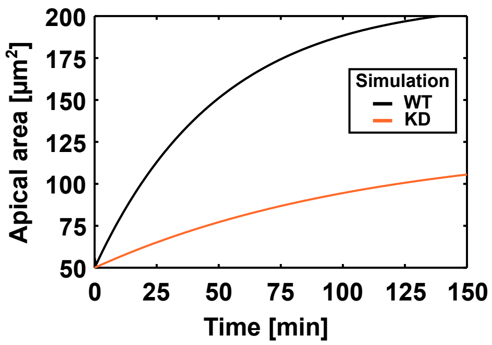

**Figure M3.** The growth of apical surface area ($\mu$m$^2$) as a function of time (in mins) for the WT (black) and KD (orange).

Basal bodies appear at the apical surface following a steady recruitment process that occurs at a constant rate throughout the simulation period. This continuous appearance of new basal bodies mimics the temporal dynamics of basal body insertion observed experimentally. As shown in Figure M4, the cumulative number of basal bodies increases linearly with time, reflecting the steady-state recruitment mechanism implemented in our model. The basal bodies do not appear uniformly, but their appearance is biased as observed experimentally.

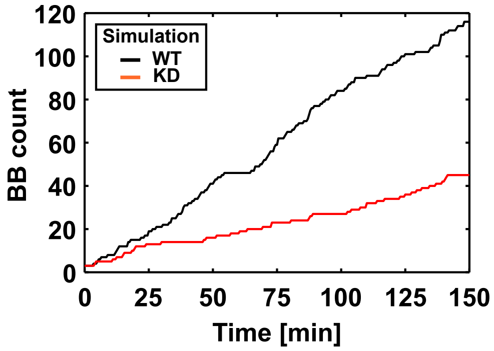

**Figure M4.** The basal bodies steadily appear in the apical surface from the basal surface. The black and red curves, respectively, corresponds to WT and KD.

The appearance of basal bodies follows a spatially dependent probability distribution that varies with the normalized distance from the cell center. Figure M5 shows the probability density functions for basal body appearance, revealing the spatial preferences that govern where new basal bodies are most likely to emerge during the simulation.

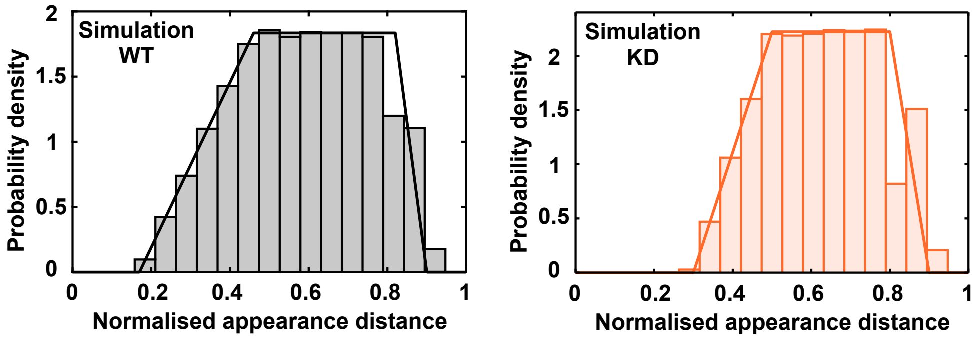

**Figure M5.** The probability of appearance of basal bodies in the apical surface as a function of normalized radius of the apical surface for WT (left) and KD (right) simulations using a model that closely resembles the experimentally observed basal body appearance. The distribution is trapezoidal: zero for $\rho < a$, rising linearly from $\rho = a$ to $\rho = b$, uniform between $\rho = b$ and $\rho = c$, falling linearly to zero at $\rho = d$, and zero for $\rho > d$. For WT, $(a, b, c, d) = (0.17, 0.46, 0.82, 0.90)$ and for KD, $(a, b, c, d) = (0.30, 0.50, 0.80, 0.90)$, where $\rho$ is the normalised radial distance from the cell centre.

After appearance, the basal bodies undergo anomalous diffusion as observed in the experiments. To mimic this, each basal body in the simulation is characterized by an anomalous exponent $\alpha$ that is randomly sampled from a Gaussian distribution with mean $\mu = 0.85$ and standard deviation $\sigma = 0.2$ for WT and $\mu = 0.72$ and $\sigma = 0.2$ for KD simulations. This parameter controls the dynamics of movement of basal bodies within the apical surface. The fBM increments are generated using the Davis-Harte method as described above for the fine-grained case. The raw diffusion strength $D_r$ is normalised using a physical reference time $t_{\text{ref}} = 30$ s (the coarse-grained frame interval) such that $\text{MSD}(t_{\text{ref}}) = 4D_r\, t_{\text{ref}}$ is consistent across all realisations, i.e., $D = D_r \cdot t_{\text{ref}}^{1-\alpha}$. Note that $t_{\text{ref}} = 30s$ here is the coarse-grained frame interval, distinct from $t_{\text{ref}} = 10s$ used in the fine-grained model which was the mean elbow transition time. Both serve the same normalisation purpose but are chosen to match the characteristic timescale of each model. When the time-dependent reduction in anomalous dynamics is used, each BB trajectory is generated as three consecutive Davis–Harte segments, each with a fixed Hurst exponent $H = \alpha/2$. The Hurst exponent is reduced between successive segments by a factor of 3/4, with a lower bound $\alpha_{\min} = 0.4$, and the diffusion-strength normalisation is recomputed separately for each segment using the same reference time $t_{\text{ref}}$. This provides a piecewise approximation to the progressive confinement observed experimentally, rather than a continuously varying fractional process.

The model incorporates explicit basal body interactions through a self-repulsion mechanism mediated by deformable actin pockets surrounding each basal body. These actin-rich regions create localized zones of altered mechanical properties that influence the positioning and movement of neighboring basal bodies. Figure M6 illustrates the geometric configuration of these interactions, showing how the actin pocket radius and basal body separation distance determine the strength of repulsive forces between adjacent structures. Denoting by $d = |\mathbf{r}_{ij}|$ the center-to-center distance between basal bodies $i$ and $j$, and by $r_o = r_b + r_s$ the outer radius of the actin pocket around a single basal body, the force exerted by basal body $j$ on basal body $i$ is calculated as:

$$\mathbf{F}_{ij} = \begin{cases} \mathbf{0} & \text{if } d \geq 2r_o \\ k_s(2r_o - d)\dfrac{\mathbf{r}_{ij}}{|\mathbf{r}_{ij}|} & \text{if } 2r_b \leq d < 2r_o \\ [2k_s r_s + k_b(2r_b - d)]\,\dfrac{\mathbf{r}_{ij}}{|\mathbf{r}_{ij}|} & \text{if } 0 \leq d < 2r_b \end{cases}$$

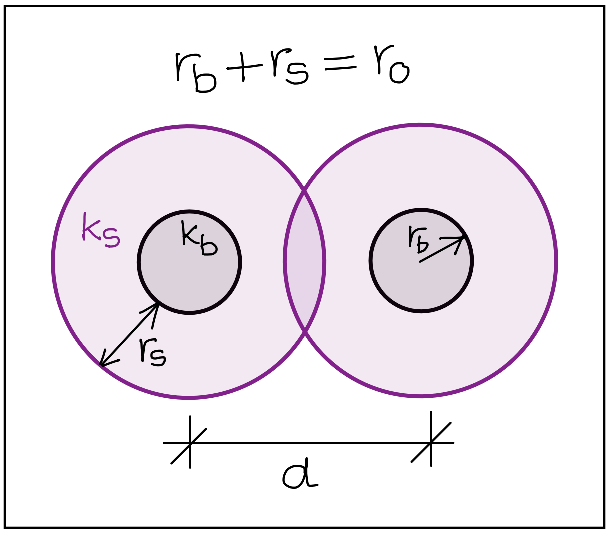

**Figure M6.** Mechanical interaction between two basal bodies. The radius of the dark basal body is $r_b$. The effective thickness of the surrounding actin pocket is $r_s$, so that the outer radius of the pocket is $r_o = r_b + r_s$. The basal bodies interact mechanically with each other when the center-to-center spacing satisfies $d < 2r_o$. The stiffness of the softer actin pocket and the stiffer basal body is $k_s$ and $k_b$, respectively.

The idea is that when $2r_b \leq d < 2r_o$, the softer actin pocket first resists the approach of the basal bodies. When the actin pocket of thickness $r_s$ is fully compressed, the stronger self-exclusion resistance between the stiffer basal bodies is activated, with the hard-core branch containing the maximum soft-shell force plus the additional hard-core contribution. Hence, the equation of motion of basal body $i$ is given as:

$$\frac{d}{dt}\mathbf{r}_i = \sum_j \mathbf{F}_{ij} + \boldsymbol{\eta}_i(t), \tag{8}$$

where the anomalous noise is implemented to be the same as in Eq. 3, but here used for the BB directly instead of the trap as was done earlier. $\mathbf{F}_{ij}$ is the force between basal bodies $i$ and $j$ due to interaction shown in Figure M6 and described in Eq. 4. We note that without the actin pockets, the basal bodies can approach each other at distances corresponding to $r_b \approx 0.25$ $\mu$m. This would lead to the observed spacing between the basal bodies to be much lesser than that experimentally observed indicating the presence of an additional steric cushion that naturally maps to actin pockets in our model.

Basal bodies are constrained to remain within the apical surface. If a basal body strays outside the apical surface of radius $R(t)$, we project it back just inside the current domain, at a small radial offset from the boundary along the local radial direction.

While most of the simulation procedures for the KD condition are similar, the experimental condition introduces a few key modifications to the base model that reflect the altered cellular environment created by $\alpha$-actinin-1 morpholino treatment. These changes capture the essential features of $\alpha$-actinin-1 morpholino induced disruptions to normal basal body organization and positioning.

Firstly, in the KD simulation, basal bodies exhibit a strong spatial bias in their appearance pattern, preferentially emerging within a single quadrant of the apical surface rather than distributing uniformly across the entire area. This quadrant-specific recruitment pattern reflects the polarized disruption of cellular organization that occurs following $\alpha$-actinin-1 morpholino treatment. Figure M7 illustrates this spatial bias by dividing the circular apical surface into four equal sectors and highlighting the preferred region-1 of basal body emergence.

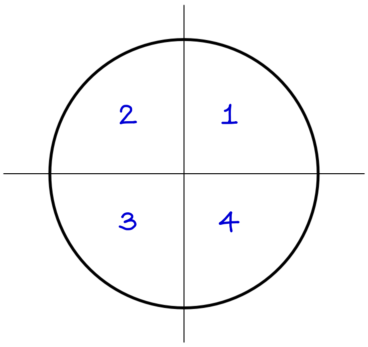

**Figure M7.** For the morpholino case, we model that the basal bodies appear in quadrant 1 at any instant with a probability of 0.7, and with a probability of 0.5 in the rest of the three quadrants combined.

Similarly, to accurately capture the localization of basal bodies observed in $\alpha$-actinin-1 morpholino treated cells, the diffusion strength parameter for the basal bodies is reduced by a factor of 5 throughout the rest of the apical surface, with the exception of the preferred quadrant-1. This modification creates a form of mini-confinement that restricts basal body mobility and prevents the normal redistribution processes that would otherwise occur in WT conditions.

Finally, the thickness of actin pockets surrounding individual basal bodies is reduced in the KD model compared to the WT case. This reduction reflects the altered cytoskeletal organization that accompanies $\alpha$-actinin-1 morpholino treatment and affects the local mechanical environment experienced by each basal body and allow the basal bodies to cluster more near each other as compared to the WT. The modified actin pocket geometry influences the strength of inter-basal body interactions. The full list of coarse-grained simulation parameters is given in Table 2.

### 5 Results

Our analysis focuses on several key metrics that capture different aspects of basal body organization and dynamics as was done in the experiments. These measurements provide quantitative insights into the differences between the wild-type (WT) and knockdown (KD) conditions. The quantification of various quantities from the simulations closely follows their experimental counterparts.

First, we analyze the organization of the basal bodies within the apical surface as a function of time. For example, we would like to quantify if the basal bodies are uniformly organized or more randomly scattered within the apical surface. One of the simplest approaches for that is to quantify the variation in apical area $\tilde{a}$ associated with individual basal bodies – $\tilde{a}$ is easiest obtained from the Voronoi tessellation of basal body spatial collection and delimited with the apical surface boundary. If $\tilde{a}$ is similar across the basal bodies, it is indicative of their uniform surface distribution. On the other hand, variation in this quantity is indicative of more scattered spatial distribution of the basal bodies. In Figure M8, as done for the experimental data, we plot the normalized statistical variance in $\tilde{a}$ as a function of time for WT and KD. As expected, the variance is higher at earlier times, indicating that the basal bodies are scattered in the apical surface due to their low numbers. However, as the number of basal bodies increase over time, the density assisted mechanical interactions between the basal bodies leads to their getting organized in the apical surface, as reflected in the decreasing normalized variance. Finally, we note that the variance in $\tilde{a}$ for KD case is mostly comparable to that for WT.

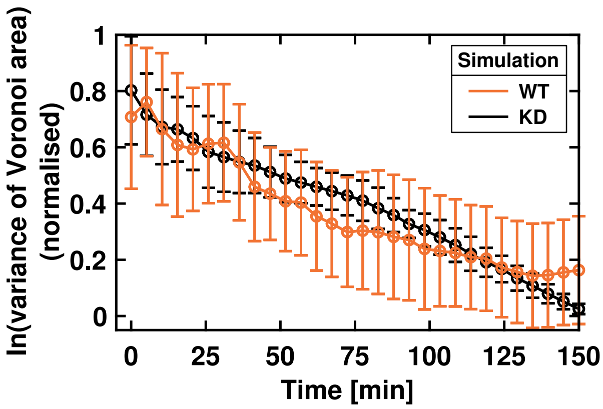

**Figure M8.** Normalized variance in area associated with basal bodies in the apical surface as a function of time for WT and KD cases. The error bars represent mean ± SD of data averaged over 20 simulations, using 5 min bins.

Another way to quantify the organization of basal bodies in the simulations is by calculating $d_{\mathrm{min}}$, i.e., the minimum pairwise distance between basal bodies. The average minimum distance between neighboring basal bodies provides insight into the typical spacing that develops within the population. Figure M9 shows how this average minimum distance evolves over time for both conditions and also the full distribution of minimum distances to capture the variability in local spacing patterns. We find that, similar to the experiments, the average value of $d_{\mathrm{min}}$ decreases with time and saturates to a value greater than 1.0 $\mu$m for the WT case and lesser than 1.0 $\mu$m for the KD case. In addition to calculating $d_{\mathrm{min}}$, we also quantified the probability distribution for $d_{\mathrm{min}}$ across all basal bodies at all times and over multiple ($N = 20$) simulation realisations. As expected, we find that for the KD cases, lower basal body spacings are favored since the actin pockets in KD that resist the pairwise approach of basal bodies towards each other are less developed as compared to the WT. We also note here that the specifics of area variance (Figure M8) are dependent on the kinetics of appearance of basal bodies in the apical surface (Figure M4) and the associated apical surface dynamics (Figure M3). Similarly, the mechanics of interaction between basal bodies via actin pockets is also instrumental in dictating the spacing between the basal bodies and hence their overall arrangement in the apical surface.

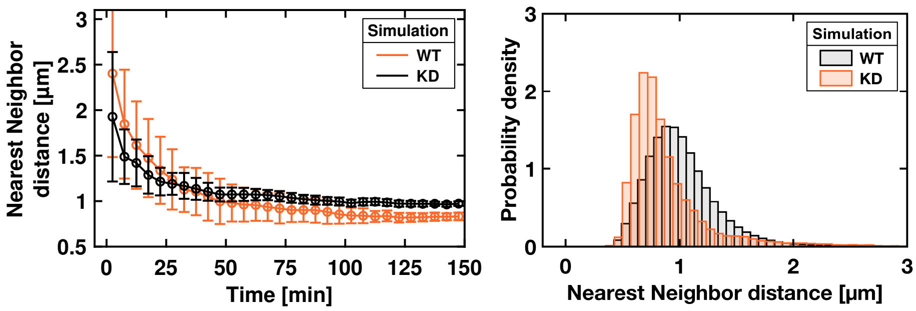

**Figure M9.** (Left) Average minimum spacing $d_{\min}$ between basal bodies as a function of time for WT and KD simulations. The error bars represent mean ± SD of data averaged over 20 simulations, using 5 min bins. (Right) Histogram for $d_{\min}$. As expected, $d_{\min}$ decreases with time and mimics the experimental observations. Data were obtained from 20 simulation runs.

The average pairwise distance $d_{\text{avg}}$ between all basal bodies offers a complementary perspective on the global organization of the basal body population. This metric reflects the overall dispersion and clustering tendencies within the system. Figure M10 tracks the temporal evolution of average inter-basal body distances and also show the complete distribution of pairwise distances at specific time points. We find that, in general, $d_{\text{avg}}$ increases with increasing area. However, the probability distribution for $d_{\text{avg}}$ is not uniform, but instead shows a peak at $\approx 6$ $\mu$m – the final radius of the apical surface is $\approx 8$ $\mu$m, for reference. The location of this peak is influenced by the basal body appearance probabilities (Figure M5). In the case of KD, the peak appears at $\approx 3.5$ $\mu$m – the final radius of the apical surface in the case of KD is $\approx 5.6$ $\mu$m (Figure M3). In this case, the lower value of the peak, as compared to WT, is indicative of the smaller apical area, preferred clustering in one quadrant, and preferential appearance of basal bodies.

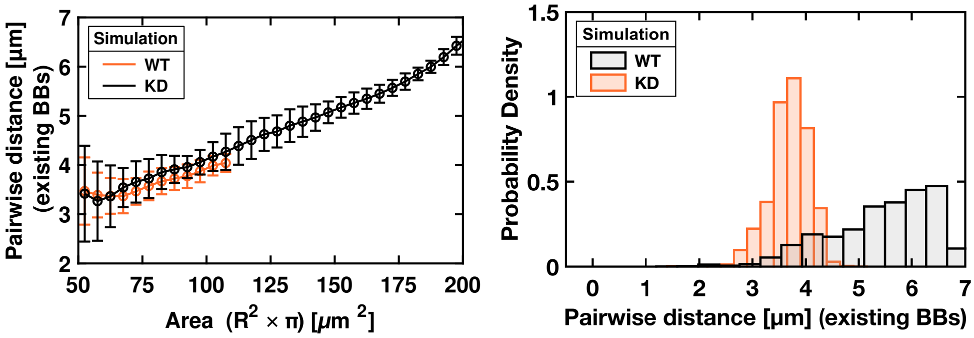

**Figure M10.** (Left) The mean pairwise spacing between the basal bodies at any given instant for WT (black) and KD (orange) based simulations. The error bars represent mean ± SD of data averaged over 20 simulations, using 5 $\mu$m$^2$ area bins. (Right) Histogram for the pairwise spacing. Data were obtained from 20 simulation runs.

We do yet another independent analysis to quantify how well-spread globally the basal bodies in the apical surface and if they form any local clusters. Such clustering analysis would provide insight into both local and global organization patterns within the basal body population. We quantify clustering tendencies using appropriate statistical measures that capture the degree to which basal bodies aggregate into distinct groups rather than maintaining uniform spatial distributions. The global clustering coefficient (GC) is calculated as follows

$$GC = \sqrt{\frac{(\text{Clust}_\theta)^2 + (1 - \text{Disp}_{r,\text{norm}})^2}{2}}, \tag{9}$$

where

$$\text{Clust}_\theta = \left| \frac{1}{N} \sum_{k=1}^{N} e^{i\theta_k} \right| = \frac{1}{N} \sqrt{\left( \sum_{k=1}^{N} \cos\theta_k \right)^2 + \left( \sum_{k=1}^{N} \sin\theta_k \right)^2}, \tag{10}$$

$$\text{Disp}_{r,\text{norm}} = \min\left(1, \frac{\sigma_r}{R/\sqrt{18}}\right) \tag{11}$$

The idea is to calculate if the basal bodies are uniformly spread across the apical surface both azimuthally and radially. The azimuthal coefficient $\text{Clust}_\theta$ (Eq. 10) would be zero if all the basal bodies are uniformly distributed. Similarly, if the basal bodies are radially uniformly distributed in a circle of radius $R$, then the ideal standard deviation of their distribution would evaluate to $R/\sqrt{18}$. Hence, for basal bodies uniformly scattered across the apical surface $\text{Disp}_\text{r} \approx 1$. For such case $GC \approx 0$ indicate the absence of global clustering. In the extreme limit in which all the basal bodies are gathered in a narrow region, then $GC \approx 1$, indicating extreme clustering. On the other hand, even if $GC = 0$, i.e., no global clustering, it could still be possible that there are patches in which the basal bodies form local clusters. Hence, in order to quantify local clustering, we calculate

$$LC = \max\left(0, 1 - \frac{d_\text{obs}}{d_\text{hex}}\right) \tag{12}$$

where,

$$d_\text{obs} = \frac{1}{N} \sum_{i=1}^{N} \left( \min_{j \neq i} \sqrt{(x_i - x_j)^2 + (y_i - y_j)^2} \right) \tag{13}$$

$$d_\text{hex} = R\sqrt{\frac{2\pi}{N\sqrt{3}}} \tag{14}$$

Here, the average value of $d_\text{min}$ is compared with the value expected for hexagonal packing. For regular arrangement, $LC \ll 1$, whereas for clustered patches $d_\text{obs} \ll d_\text{hex}$ due to which $LC \approx 1$ (Eq. 12). Figure M11 shows the temporal evolution of both global and local clustering coefficients for WT and KD simulations. We find that, as expected, at early times, local and global clustering coefficients are both large for WT as well as KD case. As time-progresses, the global clustering decreases and tends to 0 for WT as the basal bodies get scattered across the apical surface due to stochastic appearance in the apical surface followed by anomalous diffusion. In the case of KD, since the basal bodies remain confined in one quadrant, the global clustering coefficient remains saturated at a relatively higher value even at later times.

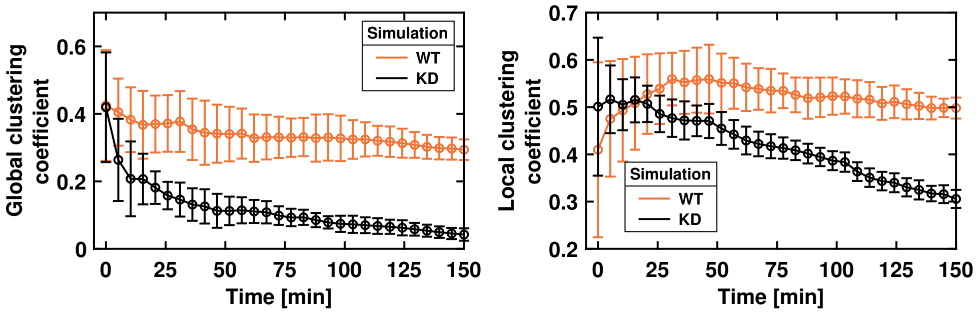

**Figure M11.** (Left) Global clustering coefficient for WT and KD based simulations. (Right) Local clustering coefficient for WT and KD based simulations. The error bars represent mean ± SD of data averaged over 20 simulations, using 5 min bins.

The local clustering, too is larger at early times for the WT, and decreases slowly to a lower but non-zero value at later times, indicating that the distribution of the basal bodies is getting more ordered but not reached hexagonal distribution, at least within the observed time-window of 150 min. For the KD case, the local clustering too is higher when compared to the WT since the actin pockets are smaller thus facilitating closer approach of basal bodies to each other.

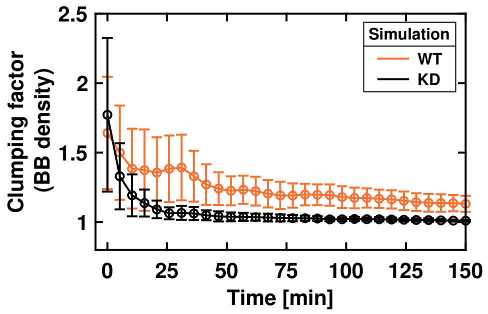

**Figure M12.** Clumping of basal bodies in one quadrant from the model for WT and KD case. The error bars represent mean ± SD of data averaged over 20 simulations, using 5 min bins.

The last aspect of our basal body configurational analysis examines clumping behavior to understand how basal body segregation patterns differ between WT and KD simulations. In this analysis, for any time-frame, we obtain $n_1, n_2, n_3, n_4$, the number of basal bodies in quadrants $1, 2, 3, 4$ and then calculate the clumping factor

$$\mathrm{CF} = 4\frac{n_1^2 + n_2^2 + n_3^2 + n_4^2}{(n_1 + n_2 + n_3 + n_4)^2} \tag{15}$$

averaged over multiple repeats of simulations $N_{\mathrm{rep}} = 20$ and within a time-window $t, t+\Delta t$. Here, if the basal bodies are uniformly distributed across all quadrants, $CF \approx 1$. If all the basal bodies are confined within one quadrant then $CF \approx 4$. Figure M12 quantifies the degree of clumping over time, showing clear differences in segregation dynamics between the WT and KD simulations.

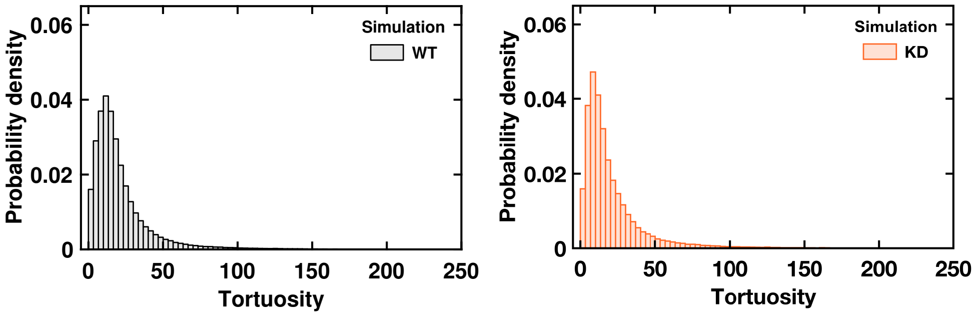

**Figure M13.** Tortuosity of WT (left) and KD (right) probability distribution. Data were obtained from 20 simulation runs.

Finally, we also examine the trajectory evolution of individual basal bodies by calculating the tortuosity of their movement paths over time. Tortuosity quantifies the degree to which basal body trajectories deviate from straight-line motion. For a given trajectory

$$\mathrm{To}(t) = \frac{\sum_t |\mathbf{r}(t+\Delta t) - \mathbf{r}(t)|}{|\mathbf{r}(t) - \mathbf{r}(0)|} \tag{16}$$

providing insight into the local constraints and interactions that influence movement patterns. The combined probability distribution of tortuosity values across the population reveals the heterogeneity in movement behavior. Figure M13 presents the probability distribution of tortuosity values. Interestingly, we find that that the tortuosity distribution for both WT and KD simulations show similar behavior indicating that the differences between WT and KD is likely not governed by basal body movements within the apical surface, but by confinement within domains of apical surface and mechanical interactions between the basal bodies.

All simulations and postprocessing was performed in MATLAB and the code is publicly available on GitHub[6].

### 6 Simulation parameters

**Table 1:** Fine-grained model simulation parameters

| Parameter | Value | Physical meaning |
|---|---|---|
| $D_w$ | 0.3 $\mu$m$^2$/s | Diffusivity of the BB within the harmonic trap; sets the short-time caging MSD plateau $\Delta R_0^2 \approx 4D_w\tau_w$. The Stokes–Einstein value for a sphere (BB) of radius $r_b = 0.25$ $\mu$m in plain water is $k_BT/6\pi\eta r_b \approx 1$ $\mu$m$^2$/s at room temperature; $D_w = 0.3$ $\mu$m$^2$/s is $\approx 30\%$ of this, consistent with the BB moving within a viscous actin-rich pocket rather than free solution. |
| $\tau_w$ | 0.005 s | Trap relaxation time; sets the short-time plateau via $\Delta R_0^2 = 4D_w\tau_w \sim 10^{-2}$ $\mu$m$^2$. Chosen so that the plateau is flat well before the earliest plotted lag time ($t = 0.02$ s). Note: $\tau_w$ governs short-time caging and is distinct from the elbow time $\tau \sim 10$ s at which the crossover to anomalous diffusion occurs. |
| $D_{\mathrm{raw}}$ | 0.0161 $\mu$m$^2$/min | Mean anomalous diffusion strength of the trap before normalisation; fitted from fine-timed experimental trajectories. |
| $t_{\mathrm{ref}}$ | 10 s | Reference time for normalising $D$; set to the mean elbow transition time $\tau \sim 10$ s from experiments. The normalisation $D = D_{\mathrm{raw}} \cdot t_{\mathrm{ref}}^{1-\alpha}$ ensures $\mathrm{MSD}(t_{\mathrm{ref}}) = 4D_{\mathrm{raw}}\, t_{\mathrm{ref}}$ is identical across realisations with different $\alpha$. |
| $\alpha$ | $\mu = 0.85$, $\sigma = 0.2$ | Anomalous exponent of the trap (MSD $\propto \Delta t^\alpha$, $H = \alpha/2$); sampled per realisation from a Gaussian, clipped to $[0.4, 1.4]$. |
| $T_{\mathrm{max}}$ | 25 min | Total simulation duration, matching fine-grained experimental imaging window. |
| $dt$ | 0.0005 s | Integration time step; $dt/\tau_w = 0.0005/0.005 = 0.1 \ll 1$ ensures Euler–Maruyama accuracy for the harmonic trap. |
| $N_{\mathrm{fun}}$ | 30 | Number of independent realisations over which the MSD is ensemble-averaged. |

**Table 2:** Parameters for coarse-grained simulations

| Parameter | WT | KD | Physical meaning |
|---|---|---|---|
| $k_b$ | 1.0 | 1.0 | BB-core repulsion stiffness. Drag coefficient set to unity in the simulation; effective unit upon dimensionalisation is $s^{-1}$. |
| $k_s^{\text{fac}}$ | 1000 | 1000 | Dimensionless ratio; soft actin-shell stiffness $k_s = k_b/k_s^{\text{fac}}$, i.e. $10^3\times$ softer than the BB core. |
| $r_o$ | 0.8 $\mu$m | 0.4 $\mu$m | Outer actin-pocket radius ($r_o = r_b + r_s$); sets the range of soft-shell repulsion. Reduced in KD, reflecting weaker actin pockets after $\alpha$-actinin-1 knockdown. |
| $r_b$ | 0.25 $\mu$m | 0.25 $\mu$m | BB hard-core radius; minimum realisable centre-to-centre separation is $2r_b = 0.5$ $\mu$m. |
| $D_r$ | 0.0285 $\mu m^2$/min | 0.0317 $\mu m^2$/min | Anomalous diffusion strength, normalised so $\text{MSD}(t_{\text{ref}}) = 4D_r t_{\text{ref}}$ at $t_{\text{ref}} =$ 30 s. Fitted to experimental BB displacement distributions. |
| $\alpha$ | $\mu = 0.85$, $\sigma = 0.2$ | $\mu = 0.72$, $\sigma = 0.2$ | Anomalous exponent ($\text{MSD} \propto \Delta t^{\alpha}$); sampled per BB from a Gaussian. Lower mean in KD reflects reduced mobility due to weakened actin cross-linking. |
| $N_{\text{bb}}$ | 120 | 45 | Final BB count at saturation, set by experimentally observed values at end of apical expansion. |
| $T_{\text{sat}}^{\text{BB}}$ | 140 min | 130 min | Time at which BB number reaches $N_{\text{bb}}$ from starting value $N_{\text{bs}}$; appearance rate $k_{\text{on}} = (N_{\text{bb}} - N_{\text{bs}})/T_{\text{sat}}^{\text{BB}}$. |
| $dt$ | 0.0625 s | 0.0625 s | Integration time step; chosen so that $k_{\text{bb}} \cdot dt \ll 1$, ensuring numerical stability of the stiff BB-core repulsion. |

### Methods

The sections below describe the image-processing workflows and data-analysis methods. All image processing and analyses were performed using ImageJ macros and MATLAB, and the corresponding code is publicly available on GitHub[7].

### 7 Image analysis for coarse-timed experiments

In this section of methods, image analysis performed on coarse-timed datasets acquired by confocal microscopy at 30–60 s intervals over a total duration of 2.5–3 h are described. After the completion of acquisition, only cells meeting the following criteria were selected for analysis: (i) an initial apical surface area of $\leq 75\,\mu m^2$, (ii) complete migration of BBs from the basal to the apical surface by the end of the acquisition, and (iii) sustained cellular viability for at least 2.5 h of imaging.

#### 7.1 Preprocessing of image stacks

##### 7.1.1 3D drift correction

The first preprocessing step consisted of correcting three-dimensional drift across time to ensure spatial alignment of all z-slices. This was performed using the ImageJ plugin Poorman3Dreg (developed by Michael Liebling, University of California Santa Barbara), applying a rigid body transformation with maximum intensity projection based registration.

##### 7.1.2 Z registration

Following 3D drift correction, image stacks were cropped in xy to isolate the MCC and remove surrounding tissue. Z registration was then performed to correct for misalignment of z-slices across time points arising from repeated manual adjustments of the z-stack range during acquisition. Such misalignment can result in the apical plane appearing at different z positions over time, leading to inconsistent visualization of BBs during apical expansion. Z registration was achieved using a custom made script that exploits the high actin signal at the apical cortex as a reference. For each time point, the script identifies the z-slice with the highest actin intensity, corresponding to the apical plane, and assigns it to a fixed z index across the entire time series. All other z-slices are repositioned accordingly. This transformation is then applied identically to the BB channel, resulting in actin and BB image stacks that are registered in z and aligned to a common apical reference plane across time. Consequently, both actin and BB channels were required for this registration step.

##### 7.1.3 Z projection of the apical plane

After z registration, a z-projection of the apical plane was performed to account for curvature of the apical surface during expansion. In some cells, the apical cortex exhibited slight curvature along the z-axis, with peripheral regions appearing in adjacent z-slices relative to the center of the apical domain. This curvature can introduce variability in BB fluorescence intensity across the apical surface and impair reliable BB detection and tracking. To correct for this effect, a custom made script was used to perform maximum-intensity z-projection over those slices spanning the curved apical plane. The extent of curvature was assessed manually by scrolling carefully through the z-stack, and typically encompassed 3–5 z-slices (corresponding to ∼1.2 µm–2 µm). These slices were selectively projected while leaving all other z-slices unchanged. The resulting projected slice defined a flattened representation of the apical surface. This procedure was applied identically to both actin and BB channels, and the projected apical slice was isolated to generate a 2D time series.

##### 7.1.4 2D registration of apical surface stacks

The resulting 2D apical surface time stacks were further registered in xy to remove any residual lateral drift remaining after earlier corrections. Registration was performed jointly on actin and BB channels using the Poorman3Dreg plugin with rigid-body transformation and maximum-intensity projection options.

Unless stated otherwise, all subsequent image processing, quantification, and analysis were performed exclusively on these isolated, maximum-intensity–projected, and 2D-registered apical surface time stacks. Throughout the Methods and main text, references to the "apical surface" or "apical domain" correspond to this processed dataset.

#### 7.2 Image processing

##### 7.2.1 Apical area contour extraction

The first step of image processing was the identification of the apical boundary, defined as the periphery of the apical surface of MCCs. This boundary was extracted from the actin channel, which clearly delineates the apical surface. To achieve this, contrast across the apical surface was first normalized using the *Enhance Contrast* function in Fiji, applied uniformly across the entire time series. The images were then smoothed using a Gaussian blur with a high sigma value ($\sim 8 - 12$) to homogenize intensity across the apical surface. The processed images were subsequently binarized to distinguish the apical surface from the background. These steps were iteratively repeated using different contrast enhancement parameters, Gaussian blur sigma values, and binarization algorithms to obtain an optimally binarized stack with a uniform apical surface. The resulting binary stacks were carefully inspected for defects within the apical domain, which were manually corrected where necessary. ROIs corresponding to the apical domain at each time point were then generated using the *Analyze Particles* function and added to the *ROI Manager* in Fiji. All extracted ROIs were verified against the original actin image stacks used for binarization. Any discrepancies were corrected either by repeating the above processing steps or through manual adjustment until the ROIs were well matched with the actin-defined apical surface.

##### 7.2.2 Identification of the apical peripheral cortex

The next step involved extracting ROIs corresponding to the actin rich peripheral cortex within the apical domain. Along the z-axis, the apical side of MCCs is enriched in actin and is referred to as the apical cortex. In contrast, within the plane of the apical surface, the cell periphery exhibits higher actin intensity than the interior of the apical area. This actin enriched region is referred here as the apical peripheral cortex. Throughout this study, intensity measurements within the apical domain, particularly for high-resolution imaging data, were normalized to the actin intensity of the apical peripheral cortex. For simplicity, this normalization is referred to as "normalized by cortex intensity". To generate ROIs corresponding to the apical peripheral cortex, the previously obtained apical domain contour ROIs were radially shifted inward by varying distances to identify the appropriate inner boundary of the peripheral cortex. Once the optimal shift value matching the thickness of the apical peripheral cortex was determined, peripheral cortex ROIs were generated for the entire time series.

The ROIs of the apical domain contour and apical peripheral cortex were applied to the original actin image stacks to extract geometric and intensity-based parameters for each time point, including apical surface area, centroid position, major and minor axis lengths, mean apical actin intensity, as well as the area and actin intensity of the apical peripheral cortex. All analyses, except for contrast enhancement, Gaussian blurring, and binarization, were performed using custom-made scripts.

##### 7.2.3 BB detection

BB detection was performed on the apical surface BB channel to enable reliable tracking. A constant intensity offset was first subtracted from the BB channel using the *Subtract* operation under *Process→Math* in Fiji, followed by intensity normalization using *Enhance Contrast*. BBs were then enhanced using either Difference of Gaussians (DoG) or Laplacian of Gaussian (LoG) filtering implemented via the GDSC plugin[8] to improve spot definition. BB positions were identified using the *Find Maxima* function with an empirically determined prominence value and the output set to single-point detections. This resulted in a binarized image stack in which BBs were represented as discrete dots across time. The binarized BB stack was manually verified against the original images, and detection parameters were iteratively refined until maximum correspondence was achieved.

##### 7.2.4 BB tracking

BB tracking was performed using a two-step approach consisting of manual tracking followed by automated tracking, with manual tracking used to guide parameter selection for automated analysis.

**7.2.4.1 Manual tracking:** Approximately 5–10 BBs with the longest visible lifetimes were manually tracked in the original BB image stack using the Manual Tracking plugin (by Fabrice Cordelières) in ImageJ. Track overlays and trajectory files containing pixel coordinates were exported. From these data, average step distances were calculated and used to inform automated tracking parameters.

**7.2.4.2 Automated tracking:** Automated tracking was performed on the binarized BB image stack using the Particle Tracker plugin[9] in Fiji. The particle radius (typically 1–3 pixels) was chosen such that each BB was detected as an individual particle in the preview. The link range was set to 3, and the displacement field was defined using the average step distance obtained from manual tracking. Particle dynamics were modeled as Brownian motion. Following tracking, trajectories were visualized using the *Visualize All Trajectories* option. Trajectories were filtered based on minimum duration (number of frames) to retain long-lived trajectories corresponding to persistent BBs. Filter thresholds were empirically chosen such that the number of retained trajectories matched the number of BBs present at the end of apical expansion. Typically, this threshold corresponded to approximately one-third to one-fifth of the total number of frames. Trajectories retained after filtering were verified for close visual agreement with manually tracked BB trajectories. At the end of automated tracking, inflated trajectory numbers were observed and multiple short trajectories corresponding to a single BB were manifesting as individual trajectories. This effect was attributed primarily to fragmentation of long trajectories by the tracking algorithm rather than to spurious particle detection, as BB identification was based on a binarized image stack containing only BB signals. Therefore, to correct this inflation from the inclusion of shorter trajectories, downstream analyses were restricted to those trajectories with sufficient tracking confidence, as determined by their persistence over a minimum number of frames. From the filtered trajectories, BB dynamics were quantified, including directionality, entry and exit behavior, step size distributions, contour length, end-to-end displacement, drift velocity, mean squared displacement (MSD), pairwise distance distributions, and clustering metrics, which are described in the following subsections.

#### 7.3 Parameters calculated for the coarse-timed data

Image-derived quantities were used for downstream quantitative analysis. All downstream analyses were performed using custom-written scripts in MATLAB[7].

##### 7.3.1 Morphogenetic and experimental time axes

For each cell, the apical surface area was measured at every time point using the extracted contour ROIs over the entire apical expansion period ($\sim$ 2.5 h). The experimental acquisition time provided the time axis, whereas the instantaneous apical area was used as a morphogenetic time variable. All parameters were analyzed and plotted as functions of both experimental time and morphogenetic time.

When plotting parameters against apical area, the analysis was restricted to the range between $50\,\mu m^2$ and $300\,\mu m^2$. The lower limit of $50\,\mu m^2$ was chosen for two reasons: (1) MCCs that had been inserted but had not yet begun expanding could remain static for extended periods, so starting the image acquisition with cells that had roughly $50\,\mu m^2$ ensured higher possibility of apical expansion; (2) in most MCCs, the first BBs reached the apical surface around $50\,\mu m^2$ . The upper limit of $300\,\mu m^2$ corresponds to the approximate maximum area reached during the plateau of apical surface area expansion in MCCs, although fluctuations were observed during this plateau phase.

To prepare the parameter data for plotting against apical area, the following procedure was applied:

1. Parameter data corresponding to apical areas $\geq 50\,\mu m^2$ were isolated.
2. The time corresponding to $50\,\mu m^2$ was normalized to zero by subtracting it from all subsequent time points.
3. Parameter data were clipped at the maximum apical area of $300\,\mu m^2$.
4. Data were further clipped at the maximum experimental time of 2.5 h.

This procedure ensured that all parameter data corresponding to apical areas between $50\,\mu m^2$ and $300\,\mu m^2$, and within the total experimental duration of 2.5 h, were included.

It also accounted for fluctuations in apical area during the plateau phase around 300 µm$^2$. The resulting adjusted area and time axes were then used for plotting all parameters as functions of apical area and time.

##### 7.3.2 BB density

At each time point, the number of BBs was obtained by counting the binarised dots of BBs in the apical surface image. BB density was defined as the ratio of the number of BBs to the corresponding apical area measured from the contour ROI.

##### 7.3.3 Voronoi tessellation analysis

Voronoi tessellations were generated from the binarized BB positions using the Voronoi function in Fiji (*Process→Binary→Voronoi*). Tessellation edges extending beyond the apical domain were removed by applying the apical area contour ROI and clearing regions outside the contour. The areas of individual Voronoi cells were then measured using the *Analyze Particles* function in Fiji. To characterize temporal changes in BB spatial organization within a single cell, Voronoi cell areas were grouped into bins corresponding to 10% increments of apical area expansion and plotted as cumulative distribution functions (CDFs). To quantify changes in BB distribution uniformity over time, the variance of Voronoi cell areas was computed for each frame, yielding one variance value per time point. The temporal evolution of this variance was then averaged across cells. For visualization of temporal trends, Voronoi area variance was log-transformed to reduce skewness and enhance the decreasing trend over time. The log-transformed variance was subsequently normalized using min–max normalization to generate averaged plots with standard deviation. For statistical comparisons between experimental conditions, raw (non-transformed) variance values were used to preserve interpretability of absolute variability. However, variance was normalized by dividing by the square of the mean Voronoi area, equivalent to the squared coefficient of variation. This normalization yields a dimensionless measure, as variance has units of µm$^4$, which are cancelled by the squared mean Voronoi area.

##### 7.3.4 Pairwise and Minimum distance calculations

To characterize BB spatial organization, three categories of distances were analyzed:

1. Distances between BBs within the same frame (Fig. S1d);
2. Distances between newly appearing BBs in the current frame and newly appearing BBs in the preceding frame (Fig. S2c);
3. Distances between newly appearing BBs and pre-existing BBs within the same frame (Fig. S3a).

For each category, two distance metrics were computed:

- Minimum distance, defined as the shortest Euclidean distance from each BB in one group to its nearest neighbor in the other group;
- Pairwise distance, defined as the set of all Euclidean distances between BBs in one group and all BBs in the other group.

Distances were computed frame-by-frame. Here, a frame corresponds to a time point, and a group refers to the set of BBs between which distances were calculated. For example, in case (i), both groups consist of BBs within the same frame; in case (ii), the two groups correspond to newly appearing BBs in subsequent frames; and in case (iii), the two groups correspond to newly appearing and pre-existing BBs within the same frame.

##### 7.3.5 BB entry rate

BB trajectories were obtained from tracking data, where initiation of a trajectory corresponds to the appearance of a newly docked BB at the apical surface. For each trajectory, the frame of first appearance and spatial position within the apical contour were recorded. The number of newly appearing BBs in each frame was counted, and BB entry rates were quantified by binning these counts either by apical area increments of $10\,\mu m^2$ or by time intervals of 5 min, allowing visualization of the temporal evolution of BB entry.

##### 7.3.6 Spatial bias in BB appearance

To assess spatial biases in BB appearance during apical expansion, all frames containing newly appearing BBs were identified, along with the corresponding BB coordinates. For each such frame, two quantities were computed (Fig. S2b):

1. *Area fraction*, defined as the ratio of the apical contour area at a given earlier frame to the apical contour area of the current frame. This ratio corresponds to a concentric annular ("donut") region representing newly added apical area during expansion.
2. *BB count within area fraction*, defined as the number of newly appearing BBs located within each area fraction.

Because the apical surface expands over time, the contour from each previous frame defines a nested region within the current frame. Thus, for a given frame $n$, all contours from frames 1 to $n$ are retained. These nested contours partition the current apical surface into concentric regions (annular zones), each corresponding to the incremental area added between successive frames.

To illustrate the procedure, consider a frame in which four new BBs appear (e.g., Frame 4), as illustrated in Fig. S2b, and suppose these new BBs are appearing as follows:

- 1 BB within the innermost region defined by contour $A_1$,
- 0 BBs in the region between contours $A_1$ and $A_2$,
- 1 BB in the region between contours $A_2$ and $A_3$,
- 2 BBs in the outermost region between contours $A_3$ and $A_4$.

In this frame, the cumulative apical areas are $A_1$, $A_2$, $A_3$, and $A_4$, where $A_4$ represents the total apical area at that time point. The corresponding area fractions are therefore defined as $A_1/A_4$, $A_2/A_4$, $A_3/A_4$, $A_4/A_4$, with associated BB counts: 1, 0, 1, 2, respectively. Notably, the final area fraction ($A_4/A_4 = 1$) always corresponds to the newest region added during expansion at that frame. This procedure is repeated for every frame containing newly appearing BBs. As the frame number increases, the number of nested contours also increases, since all prior contours are retained. For example, in a 300 frame acquisition, when analyzing BB appearance in frame 300, the contours from frames 1 to 299 define the concentric regions within which newly appearing BBs in frame 300 are assigned. Consequently, the number of evaluated area fractions progressively increases with frame number. After processing all frames of a given cell, all pairs of $A_i/A_n$, and

*BB count in corresponding annular region* are pooled and binned according to area fraction. The binned area fractions are plotted on the x-axis and the corresponding BB counts on the y-axis (Fig. 2c). The resulting distribution is then interpreted as preferential appearance towards peripheral regions if the area fraction is closer to 1 and towards apical center if the area fraction is closer to 0. For spatial interpretability, the area fraction was additionally transformed into *distance fraction* given by $\sqrt{A_i/A_n}$. Because apical area scales with the square of radial distance, this transformation converts the area fraction into a normalized radial coordinate, where 0 corresponds to the apical center and 1 corresponds to the periphery. Plots of BB count versus distance fraction therefore directly quantify radial bias in BB appearance across the expanding apical domain.

##### 7.3.7 BB exit rate calculation

BB exit events were identified as the termination points of BB trajectories. The number of BB exits per frame was counted and binned either by apical area increments of 10 µm$^2$ or by 5 min time intervals. Averaged BB exit rates were then plotted to visualize the temporal evolution of BB removal from the apical surface.

##### 7.3.8 Directionality analysis using dot product

To quantify the directional bias of BB trajectories, the dot product between the displacement vector $\vec{u}$ and the positional vector $\vec{r}$ was computed for each trajectory:

$$\vec{u} \cdot \vec{r} = |\vec{u}|\,|\vec{r}|\ \cos\theta$$

Here, the displacement vector is defined as $\vec{u} = (u_x, u_y) = (x_{\text{end}} - x_{\text{start}}, y_{\text{end}} - y_{\text{start}})$, and the positional vector is defined as $\vec{r} = (r_x, r_y) = (x_{\text{start}} - x_{\text{centroid}}, y_{\text{start}} - y_{\text{centroid}})$. The resulting $\cos\theta$ values range from -1 to 1, where negative values indicate trajectories oriented away from the apical periphery and positive values indicate trajectories oriented toward the periphery. The magnitude of $|\cos\theta|$ reflects the strength of directional alignment. Cumulative counts of trajectories with positive and negative $\cos\theta$ values were used to quantify directional biases over time.

##### 7.3.9 Drift estimation

To quantify the advective (drift) component of BB motion, the mean velocity vector was computed for each trajectory. The angle of this vector relative to the image x-axis was used to define the trajectory's principal direction of motion. Each trajectory was then rotated such that its mean direction aligned with the x-axis. This procedure was applied independently to each trajectory, followed by ensemble averaging across all trajectories. The mean displacement along the aligned x-axis captures the coherent drift component, whereas displacement along the orthogonal y-axis reflects stochastic fluctuations around the drift direction.

##### 7.3.10 Mean squared displacement analysis

The mean squared displacement (MSD) was computed for each trajectory at multiple lag times $\tau$ as:

$$\text{MSD}(\tau) = \frac{1}{N_\tau} \sum_i \left[x(t_i + \tau) - x(t_i)\right]^2$$

where $x(t)$ denotes the BB position, and $N_\tau$ is the number of displacement pairs contributing to lag time $\tau$. For each trajectory, log–log plots of MSD versus lag time were fitted using linear regression over lag times up to 700 s to avoid noise from low-statistics at longer times. The MSD was assumed to scale as $\text{MSD}(\tau) \sim \tau^\alpha$, and the scaling exponent $\alpha$ and intercept c were extracted from $\log(\text{MSD}) = \alpha \log(\tau) + c$. For visualization, MSDs from all trajectories were plotted together with their ensemble average, and the same fitting procedure was applied to the ensemble-averaged MSD.

##### 7.3.11 Actin and BB area estimation

BB tracking data were used to generate circular ROIs centered at BB positions for all time points. ROIs were drawn with a radius of 0.25 $\mu$m, determined from manual measurements of BB diameters across multiple experiments. These ROIs defined BB occupied regions within the apical surface. Using ROI exclusion, the remaining apical area, predominantly containing actin signal, was identified. BB area and actin-dominated area were computed separately at each time point to quantify relative spatial occupancy during apical expansion.

##### 7.3.12 Clustering coefficients

Clustering coefficients were used to quantify BB spatial organization at local and global scales. Both coefficients are dimensionless and range from 0 (uniform distribution) to 1 (maximal clustering).

**7.3.12.1 Local clustering (LC):** Local clustering quantifies how closely BBs are positioned relative to their immediate neighbors within a single frame. For each BB, the shortest distance to all neighboring BBs which is referred to as the nearest neighbor distance, is calculated. The local clustering coefficients for a frame is then defined as:

$$\text{LC} = 1 - \frac{\langle d_{\text{nn}} \rangle}{1/\sqrt{\rho_{BB}}}$$

where $\langle d_{\text{nn}} \rangle$ is the mean nearest-neighbor distance and $\rho_{BB}$ is the BB density. Here, BB density is defined as the number of BBs divided by the apical area. By normalizing to the inter-BB distance ($1/\sqrt{BB\ density}$), LC accounts for differences in cell size and BB number. Values of LC close to 0 indicate a uniform, well-spaced local distribution of BBs, whereas values approaching 1 indicate tight local clustering of BBs.

**7.3.12.2 Global clustering (GC):** Global clustering quantifies the overall spatial organization of BBs within the apical domain, specifically the large scale asymmetry in BB distribution, integrating both angular and radial distributions. For a given frame, GC is calculated as a normalized Euclidean distance combining angular clustering ($Clust_\theta$) and radial dispersion ($Disp_{r,norm}$):

$$\text{GC} = \sqrt{\frac{{Clust_\theta}^2 + (1 - Disp_{r,norm})^2}{2}}$$

Angular clustering ($Clust_\theta$) is computed from the polar coordinates of the BB positions relative to the apical centroid. For each BB, the angle $\theta$ is calculated, and the angular clustering is defined as:

$$\text{Clust}_\theta = \left| \frac{1}{N} \sum_{k=1}^{N} e^{i\theta_k} \right|$$

where values range from 0 (uniform angular distribution) to 1 (all BBs aligned in the same angular direction).

Radial dispersion normalization ($Disp_{r,norm}$) accounts for differences in apical domain size and shape. For each frame, the periphery of the apical area contour is used to generate a mesh grid. Simulated particle positions are randomly distributed within this contour, and their radial distances from the centroid are computed. The standard deviation of these simulated radial distances is used to normalize the actual BB radial distances, producing a frame-specific $Disp_{r,norm}$ value. This ensures that radial dispersion is comparable across frames and cells, regardless of changes in apical area. Thus, GC integrates the normalized angular and radial components, giving equal weight to angular asymmetry and radial clustering. Higher GC values indicate stronger global clustering and asymmetry, whereas lower values indicate a more uniform distribution of BBs across the apical domain.

##### 7.3.13 Clumping factor

To compare BB spatial organization with actin intensity distributions, a quadrant-based clumping factor (CF) was computed. The clumping factor was defined as:

$$\text{CF} = \frac{\langle \rho^2 \rangle}{\langle \rho \rangle^2}$$

which is computed across four angular quadrants of the apical domain. This metric, commonly used to quantify spatial inhomogeneity, was applied both to BB density and to actin intensity, enabling direct comparison of heterogeneity across matched spatial regions.

### 8 Image analysis for fine-timed experiments

Fine-timed datasets consisted of 30,000 frames acquired over 10.5 min at 1 frame every 21 ms. Tracking of BBs across this large dataset was performed in two stages:

1. **Averaged stack generation**: Maximum intensity projections were performed for every 30 frames of the original stack, producing an averaged stack of 1000 frames.
2. **Tracking strategy**: BBs were first tracked in the averaged stack to obtain robust initial tracks. These tracks were then used as references to construct BB trajectories in the original stack.

All subsequent processing involves both the averaged and original stacks until tracking step is complete.

#### 8.1 Preprocessing of image stacks

##### 8.1.1 Drift correction

Lateral (xy) drift was corrected to ensure that measured BB movements reflect true dynamics and not stage or sample drift. This is especially important at small time intervals, where even minor drift can introduce apparent BB motion. Drift correction was performed using the StackReg plugin[10] in Fiji with rigid body transformation. The drift-corrected stack was then maximum intensity projected every 30 frames to generate the averaged stack of 1,000 frames. Both the averaged and original stacks were saved as separate files.

##### 8.1.2 BB detection

BBs in both the averaged and original stacks were detected using the ThunderSTORM plugin[11] in Fiji. Camera parameters were set as follows: pixel size = 106.667 nm, photoelectrons per A/D count = 5, base level A/D counts = 5, EM gain = 100. The resulting detected BB positions were saved separately for both stacks.

#### 8.2 Image processing

##### 8.2.1 BB tracking

Tracking was performed using a nearest-neighbor algorithm for both averaged and original stacks.

1. **Averaged stack**: Tracks were constructed by searching for the subsequent BB position in consecutive frames within a radius of 300 nm. If no particle was found, the track was advanced using the previous BB position. This process generated trajectories for all detected BBs.
2. **Original stack**: BB tracks from the averaged stack were used as references. For each BB in a track from the averaged stack, a corresponding BB was searched in the original stack within a 300 nm radius. If found, this position was used to extend the track; otherwise, the averaged stack position was used. Particles not found for $>5$ consecutive frames were excluded. Completed tracks for the original stack were saved for downstream analysis.

#### 8.3 Parameters calculated for the fine-timed data

##### 8.3.1 MSD calculation

Mean squared displacement (MSD) was calculated as described for coarse-timed data, using only trajectories with $\geq 20,000$ steps.

##### 8.3.2 Alpha value extraction

MSDs were ensemble averaged across all BBs within a single apical area over 10.5 min. Assuming MSD behaves as $MSD(\tau) \sim \tau^{\alpha}$, linear fits of $log(MSD)$ versus $log(\tau)$ were performed to extract the slope ($\alpha$) and intercept (c). fine-timed MSDs exhibited an initial constrained diffusion phase followed by a generalized diffusion phase (sub-, normal-, or superdiffusive). To capture this, two $\alpha$ values were calculated:

1. **Short-time $\boldsymbol{\alpha}$**: Linear fitting was iteratively extended from the smallest lag time until $R^2 > 0.9$ or RMSE $< 0.02$.
2. **Long-time $\boldsymbol{\alpha}$**: Fitting began immediately after the short-time regime, with maximum lag time set to $\sim$ *trajectory length*/4 to avoid noisy tails, iteratively extended until $R^2 > 0.99$ or RMSE $< 0.02$.

Both short- and long-time $\alpha$ values were extracted from ensemble averaged MSDs for each cell and plotted across the apical area range.

##### 8.3.3 Transition time extraction

The lag time at which BB motion transitions from short time constrained diffusion to a longer time generalized diffusive regime was defined as the transition time. This timescale captures the point at which BB trajectories escape local confinement and begin to exhibit sustained transport or diffusion. To extract this transition time, we used the two power law regimes identified in the MSD analysis. Linear fits to the ensemble averaged MSD in log–log space were performed separately for the short time regime and the long time regime, yielding slopes $(\alpha_s, \alpha_l)$ and intercepts $(y_s, y_l)$, respectively. The transition time $(\tau_{transition})$ was then defined as the lag time at which these two fitted lines intersect (Fig. S4c). The intersection point along the lag time axis was calculated as:

$$\tau_{transition} = \frac{y_l - y_s}{\alpha_s - \alpha_l},$$

where $y_s$ and $\alpha_s$ correspond to the short-time fit and $y_l$ and $\alpha_l$ to the long-time fit. This intersection provides an objective estimate of the timescale at which BB dynamics change regime.

##### 8.3.4 Diffusion strength calculation

The long-time MSD behavior of BB trajectories exhibited a broad range of dynamical regimes in an area dependent manner, including subdiffusive, diffusive, and superdiffusive motion, with $\alpha$ values spanning approximately 0.2–1.6 in both fine- and coarse-timed data (see figure below). Under such conditions, the conventional diffusion coefficient D, obtained from the generalized MSD relation

$$MSD(\tau) = 4D\tau^{\alpha},$$

is not directly comparable across trajectories. Specifically, when $\alpha \neq 1$, the units of D become $\mu m^2\,s^{-\alpha}$, rather than the standard $\mu m^2\,s^{-1}$. As a result, diffusion coefficients extracted from trajectories with different $\alpha$ values cannot be pooled or compared directly without violating dimensional consistency. To enable meaningful comparison across trajectories and across apical areas, we therefore defined a normalized diffusion strength by rescaling the diffusion coefficient using the transition time as a characteristic timescale. For each trajectory, the diffusion strength was calculated as:

$$D_{strength} = D \cdot \tau_{transition}^{\alpha-1}.$$

This normalization restores consistent units ($\mu m^2\,s^{-1}$) while preserving information about the underlying dynamical regime. The diffusion strength thus provides a physically interpretable measure of BB mobility that can be compared across trajectories with different $\alpha$ values.

Using this approach, the diffusion strength measured from the fine-timed data peaked at approximately 0.02 µm$^2$ min$^{-1}$. A similar normalization strategy was applied to the coarse-timed data, where the frame interval was used as the characteristic timescale, yielding a comparable diffusion strength of approximately 0.03 µm$^2$ min$^{-1}$ (see figure below).

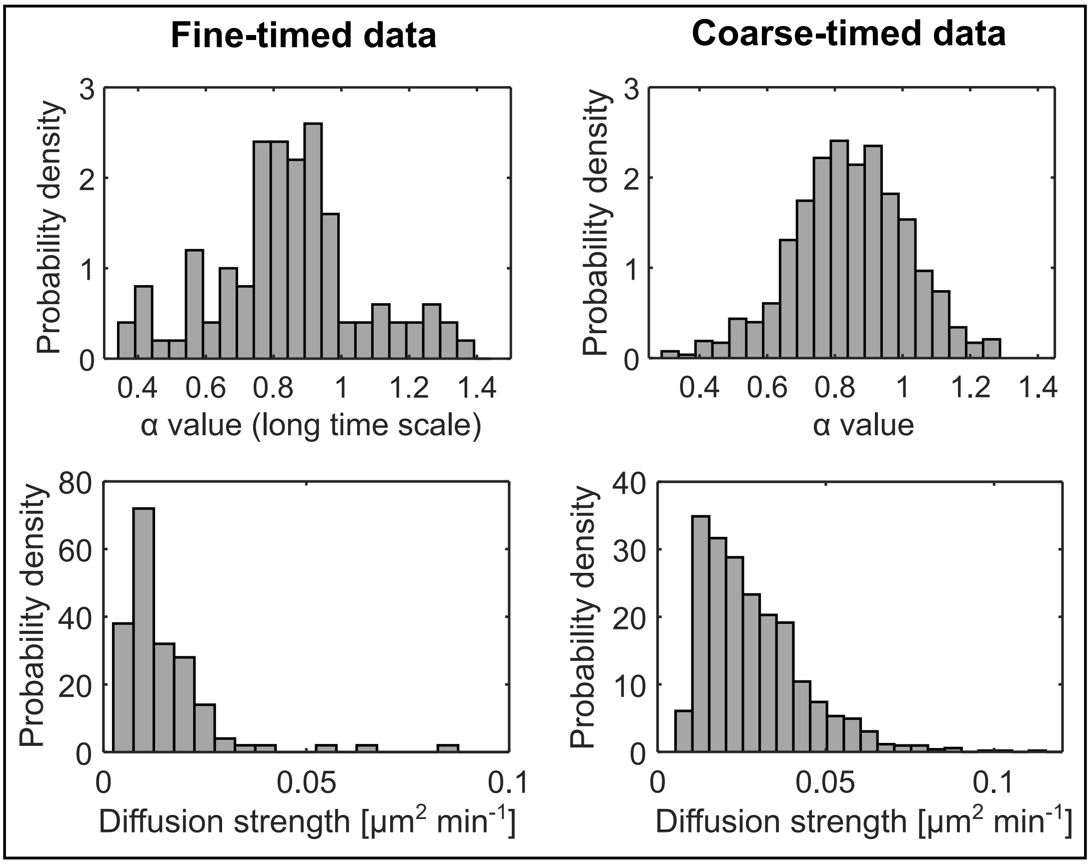

**Figure:** All data correspond to WT experiments. Plots in the first column correspond to fine-timed data, and those in the second column to coarse-timed data. The first and second rows show the probability densities of the $\alpha$ values and diffusion strengths, respectively.

##### 8.3.5 Breakpoint estimation

To identify potential changes in the relationship between apical area and dynamical parameters such as $\alpha$ values and transition times, we performed a piecewise linear regression analysis with a variable breakpoint. This analysis tests whether the data are better described by two linear regimes separated by a characteristic apical area. Breakpoint values were allowed to vary between approximately 70 and 250 µm$^2$ in steps of 1 µm$^2$. For each candidate breakpoint, the dataset was split into two subsets (below and above the breakpoint), and independent linear regressions were fit to each subset. The goodness of fit was quantified using the sum of squared errors (SSE) across both segments. The breakpoint yielding the minimum SSE was selected as the best-fit breakpoint. The resulting breakpoint locations were consistently found within the range of $\sim$ 150–200 µm$^2$

across parameters. To assess whether the identified breakpoint corresponded to a genuine change in behavior, values before and after the breakpoint were compared using appropriate statistical tests.

##### 8.3.6 Assessment of localization noise

To assess whether the features observed in the short-timescale MSDs of BBs could arise from localization noise or instrumental drift, we performed MSD analysis on immobilized fluorescent beads imaged under identical conditions. Fluorescent beads of approximately 0.5 µm diameter, comparable to the size of BBs, were attached to the coverslip and imaged using the same frame rate and acquisition duration as the fine-timed BB experiments. MSDs were calculated for individual bead trajectories and ensemble averaged using the same analysis pipeline applied to BBs. The ensemble-averaged bead MSD at the smallest lag times provides an upper bound on localization noise and residual mechanical drift in the system. We found that the ensemble-averaged MSD of the beads was approximately an order of magnitude lower than the corresponding MSD values of BBs (Fig. S4j).

To further evaluate the legitimacy of BB displacements at short timescales, we compared the root mean squared displacement (RMSD) at the first lag time for ensemble-averaged BB and bead trajectories (Fig. S4j). The RMSDs of BBs and beads were 85 nm and 25 nm, respectively. Since the BB displacement at the first lag time was more than threefold larger than that of the beads, this strongly supports that the observed BB motion reflects genuine biological displacement rather than localization noise alone.

However, immobilized beads are expected to exhibit higher fluorescence intensity than BBs, enabling more accurate localization. In contrast, the lower fluorescence signal of BBs introduces additional localization uncertainty, which contributes an additive offset to the MSD according to Martin et al.[12] $MSD = 4Dt + 2\sigma^2$, where $\sigma$ represents the localization uncertainty. Since the beads were immobilized on the surface, we used the bead RMSD of 25 nm as an empirical estimate of the experimental noise floor. To estimate the contribution of this noise floor to the measured BB displacement, we subtracted the squared bead RMSD from the squared BB RMSD and subsequently calculated the square root of the resulting value. This yielded a noise-corrected BB displacement of 77 nm, which remains approximately threefold larger than the bead displacement (25 nm). These results indicate that localization noise alone cannot account for the measured BB displacement at the first lag time.

Notably, the measured BB displacement (85 nm) is smaller than the physical pixel size (106.667 nm), necessitating verification that the localization algorithm achieves reliable sub-pixel precision and that the theoretical localization uncertainty agrees with the empirical estimate obtained from the bead measurements. BB localization in the fine-timed experiments was performed using the ThunderSTORM plugin[11] with parameters optimized for sub-pixel localization (*see Methods section 8.1.2*). The plugin reports individual $\sigma$ values for each localization event. These values were independently validated by manually calculating $\sigma$ using the expression described by Quan et al.[13], yielding consistent results. Across all BB detections in all cells, the mean and mode of the localization uncertainty distribution were approximately 20–25 nm, consistent with the experimental noise floor estimated from the immobilized beads. Together, these analyses confirm that the localization algorithm achieves reliable sub-pixel precision, that the bead displacement provides a robust empirical estimate of the localization noise floor, and that the noise-corrected BB displacement of 77 nm at the first lag time represents a legitimate biological displacement.

Taken together, these findings indicate that localization noise does not dominate the observed BB dynamics, and that the short-timescale constrained diffusion and subsequent dynamical regime transitions reflect genuine BB motion rather than measurement artifacts.

### 9 Common parameters for coarse- and fine-timed analysis

##### 9.0.1 Step distance

The BB step distance was defined as the Euclidean distance between two consecutive positions of a BB along its trajectory. For the coarse-timed dataset (frame interval: 30–60 s; total duration: $\sim 2.5$ h), step distances from all trajectories across all cells were pooled, and their probability density distributions were computed.

##### 9.0.2 Contour length

The contour length of a BB trajectory was calculated as the cumulative sum of all step distances along that trajectory. For the coarse-timed data, contour lengths from all BB trajectories across all cells were pooled and plotted as probability density distributions. For the fine-timed data, contour lengths were computed for trajectories spanning the entire apical area range. These values were binned in $50\,\mu\text{m}^2$ apical area intervals and plotted as mean $\pm$ SD to visualize how contour length evolves with increasing apical area.

##### 9.0.3 End-to-end length

The end-to-end length was defined as the direct Euclidean distance between the starting and ending coordinates of a BB trajectory. For the coarse-timed data, end-to-end lengths from all trajectories across all cells were pooled and plotted as probability density distributions. For the fine-timed data, end-to-end lengths were calculated for trajectories across the full apical area range, binned in $50\,\mu\text{m}^2$ intervals, and plotted as mean $\pm$ SD to examine their evolution during apical expansion.

##### 9.0.4 Tortuosity

Tortuosity of BB trajectories was quantified as the ratio of contour length ($d_l$) to end-to-end length ($d_r$), i.e., Tortuosity $= d_l/d_r$, and was calculated for both coarse-timed and fine-timed datasets. For the coarse-timed data, tortuosity was computed for each individual trajectory, and the resulting values were pooled and plotted as probability density distributions. For the fine-timed data, trajectories are substantially shorter in duration but sampled at much higher temporal resolution. As a result, tortuosity values were computed for trajectories grouped by apical area, binned in $50\,\mu\text{m}^2$ intervals, and plotted to assess how trajectory complexity evolves across the apical expansion. Notably, tortuosity values obtained from the fine-timed data were substantially larger than those from the coarse-timed data. This arises because the high temporal resolution captures thousands of very small BB displacements, leading to large cumulative contour lengths, while the corresponding end-to-end displacements remain small due to the short observation window. Consequently, the ratio $d_l/d_r$ becomes large. To facilitate visualization and comparison across apical area bins, tortuosity values from the fine-timed dataset were therefore natural log-transformed prior to plotting.

### 10 Image analysis for high-resolution experiments

#### 10.1 Airyscan processing

The first step in the high resolution imaging analysis pipeline was the processing of raw Airyscan data. Airyscan raw images were processed using ZEN 3 software, where the raw data were deconvolved using the Super Resolution filter, where the value was automatically estimated by the algorithm to generate the corresponding high resolution image. The default filter setting was chosen after testing a range of filter values across multiple images, during which the default value was found to consistently provide optimal deconvolution performance without introducing artifacts.

#### 10.2 Image processing

Subsequent analysis steps like the apical plane isolation, apical area contour identification, BB detection, and BB position extraction, closely follow the analysis pipeline described for the coarse-timed experiments. Therefore, only those steps that were specifically adapted or tuned for high resolution imaging are described in detail below. For a complete description of shared steps, refer to the corresponding sections in the coarse-timed analysis.

##### 10.2.1 Apical plane isolation

To isolate the apical plane containing the BBs and the associated actin meshwork, the acquired z-stack spanning the entire apical cortex was carefully examined. The apical cortex was manually identified and isolated. In cases where curvature of the apical surface was observed, the z-slices encompassing the curved apical plane were projected along the z-axis. The resulting projected image was used as a representative apical surface for subsequent analysis of the actin meshwork surrounding BBs.

##### 10.2.2 Apical area contour extraction and BB detection

The apical area contour was manually annotated using the actin channel. From this contour, the apical peripheral cortex was defined by inwardly shifting the apical boundary by a specified distance using a custom written script. This inward shifted contour defined the ROI for the peripheral cortex, which was used to quantify actin intensity at the apical periphery. Actin intensities measured within this peripheral cortex ROI were used for normalization.

BBs were detected in the BB channel by transforming the signal into binarized, discrete puncta using either a Difference of Gaussians (DoG) or Laplacian of Gaussian (LoG) filter implemented via the GDSC plugin[8], followed by identification of central maxima using the *Find Maxima* function in Fiji.

##### 10.2.3 BB position identification

BB positions were extracted from the binarized images using the Particle Tracker plugin [9]. The same workflow described in the Automated tracking section (7.2.4.2) of the coarse-timed analysis was followed, with the exception that all parameters were left at their default values. As these datasets consist of single frame images rather than time stacks, the plugin returns BB positions only for the first frame.

##### 10.2.4 Definition of BB and surrounding regions

Using the BB positions obtained above, circular ROIs were drawn around each BB with a radius of 0.25 µm. A second concentric circular ROI with a radius of 0.5 µm was then generated. The inner ROI (0.25 µm radius) was used to quantify actin intensity at the BB position, while the annular region between the 0.25 µm and 0.5 µm radii was used to quantify actin intensity in the surrounding region. The choice of 0.25 µm and 0.5 µm radii was guided by repeated manual measurements of BB diameters and the spatial extent of high intensity actin enrichment surrounding BBs. These ROI definitions enabled automated and consistent extraction of actin intensities at BB positions and in their immediate surroundings for all BBs within the apical area. All ROI generation and intensity extraction steps were implemented using a custom written script.

#### 10.3 Parameters calculated for high-resolution imaging analysis

##### 10.3.1 Actin intensity ratios at different CDF percentiles

For each apical domain, actin intensities measured at BB positions and in the surrounding regions were plotted as cumulative distribution functions (CDFs). These CDFs were used to visualize systematic differences in actin enrichment between the two regions as a function of apical area. From each CDF, the corresponding actin intensity at $25^{th}$, $50^{th}$, and $75^{th}$ percentiles were extracted. For each percentile, the ratio of actin intensity in the surrounding region to that at the BB position was calculated. These ratios were then plotted across the entire apical area range to assess how the relative actin enrichment evolves during apical expansion, across all cells.

##### 10.3.2 Ratio of ensemble-averaged actin intensity

While the CDF based analysis rigorously incorporates actin intensity measurements from individual BBs, it may introduce pseudoreplication by treating multiple BBs within the same apical domain as independent data points. To address this, a complementary ensemble averaged analysis was performed. For each apical domain, actin intensities at BB positions and in surrounding regions were averaged across all BBs within that apical domain, yielding one mean value per domain for each region. The ratio of the ensemble averaged actin intensity in the surrounding region to that at the BB position was then computed, resulting in a single ratio per apical domain. These ratios were plotted across the full apical area range and used as a complementary measure to the CDF based ratios.

##### 10.3.3 Breakpoint estimation

Changes in the relationship between apical area and (i) the CDF based actin intensity ratios and (ii) the ratio of ensemble averaged actin intensities were assessed using breakpoint estimation, following the same procedure described for the fine-timed experiments. Following breakpoint identification, data points before and after the breakpoint were statistically compared to assess the nature and significance of changes in the relationship between apical area and relative actin enrichment around BBs.

### 11 Image analysis for figures and movies

**Fig.** 1b, **2a, 5a, S1a, S6a**: For visualization purposes, the contrast of the BB channel in these figures was enhanced by subtracting a constant intensity offset from the images using the *Subtract* operation under the Math functions in Fiji.

**Fig.** 1c, **5b**: Orthogonal views shown in these figures were generated using the ClearVolume plugin[14] in Fiji.

**Fig.** 2d **and Movie 6**: BB entry and exit events shown in this figure and movie were obtained as follows. A BB of interest was first identified in the xy plane. The *Orthogonal Views* function in Fiji was then used to locate the same BB in the corresponding xz and yz planes. Images corresponding to the xy, xz, and yz planes were saved separately. This procedure was repeated for each successive step of BB movement observed in the xy plane. This approach allowed capture of BB entry and exit events at the apical plane in the xy view, while simultaneously tracking BB motion below the apical surface in the xz and yz views. The process was continued until the BB ceased its entry–exit cycles and remained stably associated with the apical domain. Following this, BB movement along the z-axis during the entry–exit cycles was quantified by manually tracking the BB in the xz plane using the Manual Tracking plugin (by Fabrice Cordelières) in Fiji, yielding a measure of BB displacement along the apico–basal axis.

### 12 Statistical analysis

Data are presented as the mean of binned values (corresponding to the intersection of the vertical and horizontal error bars), with error bars representing the standard deviation (SD), for the time- and area-evolution plots. For violin plots, individual data points are shown as dotted markers within the distribution when sample sizes are moderate. For plots with very large sample sizes, individual points are not displayed to avoid visual overcrowding. In all violin plots, the central line indicates the median, and the upper and lower lines represent the $75^{\text{th}}$ and $25^{\text{th}}$ percentiles, respectively.

For plots showing probability distributions, the peak of unimodal histogram distributions was estimated by fitting a normalized Gaussian function to the binned data. The fitted mean was used as an estimate of the peak. This procedure was used solely to obtain a smooth and consistent peak estimate and was not intended to infer the underlying distribution shape. Across all unimodal distributions analyzed, this approach yielded peak estimates that were visually consistent with the data.

For statistical comparisons, normality of the data was assessed using the Shapiro–Wilk,

D'Agostino–Pearson, Kolmogorov–Smirnov, and Anderson–Darling tests implemented in GraphPad Prism. The tests generally yielded consistent conclusions regarding normality. When minor discrepancies occurred, the overall assessment was based on the majority of tests.

For unpaired data with two groups, normally distributed data with large sample sizes were assessed using an unpaired t-test, whereas non-normally distributed data or data with small sample sizes were analyzed using the Mann–Whitney test. For unpaired data involving more than two groups, a Kruskal–Wallis test was performed, followed by Dunn's post-hoc test for pairwise comparisons.

A significance threshold of $p < 0.05$ was applied. Exact $p$ values are reported in the figure legends, and significance is denoted as follows: $p < 0.05(*)$, $p < 0.01(**)$, $p < 0.001(***)$, $p < 0.0001(****)$. All statistical analyses were performed using GraphPad Prism 11. Each experiment was repeated at least three times, and the number of cells, trajectories, and independent experiments is reported in the figure legends.

### List of materials

**Chemicals and staining reagents**

| Product | Source | Identifier |
|---|---|---|
| Chorulon | MSD Animal Health | 422741 |
| Nacl | Fisher Scientific | BP358 |
| KCL | Fisher Scientific | BP366 |
| $CaCl_2$ | Scharlab | CA01941000 |
| $MgCl_2$ | Sigma-Aldrich | M2670 |
| Cysteine | Sigma-Aldrich | C7880 |
| Ficoll | Sigma-Aldrich | F4375 |
| 3-(N-Morpholino)propanesulfonic acid (MOPS) | Sigma-Aldrich | M1254 |
| Ethylene Glycol-bis(beta-aminoethyl ether) - N,N,N',N' - tetraacetic acid tetrasodium salt (EGTA) | Sigma-Aldrich | E8145 |
| $MgSO_4$ | Fisher Scientific | BP213 |
| Formaldehyde | Sigma-Aldrich | 47608 |
| Tris-HCl (Trizma® Hydrochloride) | Sigma-Aldrich | T5941 |
| Triton X-100 | Sigma-Aldrich | T8787 |
| Fetal bovine serum (FBS) | Sigma-Aldrich | F2442 |
| Dimethyl Sulfoxide (DMSO) | Sigma-Aldrich | D2650 |
| Phalloidin 555 | Invitrogen | A34055 |
| Phalloidin Atto-594 | Sigma-Aldrich | 51927 |
| Phalloidin 647 | Invitrogen | A22287 |
| Ultra low melting point agarose | Sigma-Aldrich | A2576 |
| Low melting point agarose | Promega | V2111 |
| Cover slips (Thickness 1.5 H; 25 mm) | Marienfeld | 0117650 |
| Imaging chambers (Attofluor Cell chamber™ for microscopy) | Invitrogen | A7816 |
| High vacuum Silicone grease | Sigma-Aldrich | Z273554 |
| Fluorescent beads (0.5 µm; Yellow) | Spherotech | FP-0552-2 |

**Plasmids**

| Plasmids | Source |
|---|---|
| alpha-Tubulin-LifeAct-GFP | John Wallingford group |
| alpha-Tubulin-LifeAct-RFP | |
| alpha-Tubulin-Chibby-GFP | Brian Mitchell group |
| alpha-Tubulin-Chibby-RFP | |

| Oligomorpholinos | | |
|---|---|---|
| **Morpholino** | **Company** | **Sequence (5'-3')** |
| $\alpha$-Actinin-1 (ACTN1.L) Translation blocking | Gene Tools USA | CATAATGATCCATCC TGAGCTGCTG |
| $\alpha$-Actinin-1 (ACTN1.L) Splice blocking | Gene Tools USA | CCTGTGGCTCAAAAT CTTACCTTCA |

| Softwares | | |
|---|---|---|
| **Product** | **Company** | **Link** |
| ImageJ | - | https://imagej.net/ij/ |
| Fiji | - | https://fiji.sc |
| MATLAB R2023b | Mathworks | https://se.mathworks.com/products/matlab.html |
| GraphPad Prism 11 | GraphPad Software, LLC | www.graphpad.com |
| Inkscape 1.4 | - | https://inkscape.org |
| Adobe Illustrator 27 | Adobe | https://www.adobe.com/products/illustrator.html |
